# scRBP infers cell-specific RNA-binding protein regulons from single-cell transcriptomes at gene and isoform resolution

**DOI:** 10.64898/2026.10.13.758934

**Authors:** Yunlong Ma, Miao Tang, Vanshika Sriram, Jingjing Li, Connor Jops, Jade England, Daniel Vo, Yundan Liao, Rachel L. Smith, Michael Margolis, Ophir Shalem, Michael J. Ziller, Celine K. Vuong, Luis de la Torre Ubieta, Michael J. Gandal

## Abstract

Single-cell regulatory network inference remains dominated by transcription factors, despite RNA-binding proteins (RBPs) orchestrating major aspects of transcriptional regulation. We present scRBP, a framework that infers motif-supported RBP regulons from short- and long-read single-cell transcriptomes at gene and isoform resolution, resolves targets by transcript region (5′UTR, CDS, intron, 3′UTR), scores regulon activity in individual cells, and tests common- and rare-variant association against matched empirical nulls. From 20,746 motifs for 616 RBPs, scRBP recovered experimentally defined targets across 223 ENCODE eCLIP datasets at gene and isoform level. Across 30 tissues, transcript-region preferences separated 3′UTR-associated stability factors from intron-dominant splicing regulators. In published genome-scale CRISPRi Perturb-seq data, scRBP-inferred targets were over-represented among genes responding to RBP knockdown (1.4-2.5-fold across 161 RBPs; FDR<2×10^−4^). Across 6.2 million cells and 50 traits, scRBP prioritized cell-specific post-transcriptional programs linking human genetic risk to development and disease.

## INTRODUCTION

RNA-binding proteins (RBPs) are central regulators of gene expression, orchestrating multiple layers of RNA metabolism via co- and post-transcriptional regulation. Major established roles for RBPs include coordination of pre-mRNA splicing, RNA localization, RNA stability, cleavage and polyadenylation, and regulation of translation, among others^1,2^. Many of the 1,542 currently annotated RBPs act across multiple regulatory steps^1^. For example, the neuronal splicing factor *NOVA1* coordinates both alternative splicing and poly(A) site usage^3^. Likewise, *FMR1* is well known for repressing translation but also exerts nuclear functions in chromatin regulation^2^. These processes are essential for cellular homeostasis and organismal physiology. RBP-mediated regulation is governed in part by *cis*-regulatory RNA elements^4,5^, often with characteristic binding motif sequences inferred from experimental mapping of RBP-RNA interactions. RBP regulation also reflects dynamic RNA-protein interactions that are highly dependent on tissue- and cell type-specific expression contexts^6,7^. However, experimental maps of RBP-RNA interactions remain largely restricted to a small number of cell lines^3,8,9^, leaving the cellular specificity of RBP regulatory architectures poorly resolved.

Consistent with this central role in RNA metabolism, RBPs also exhibited strong genetic associations with human diseases^2,10^. Rare deleterious variants in RBP genes cause a spectrum of neurodevelopmental and neurological disorders, including fragile X syndrome (FXS) resulting from mutations in *FMR1*^*11*^, and spinal muscular atrophy caused by loss-of-function mutations in *SMN1*^*12*^. In addition, mutations in RBP genes such as *TARDBP* and *FUS*^*13,14*^ contribute to neurodegenerative disorders including amyotrophic lateral sclerosis. Beyond rare variants, genome-wide association studies (GWAS) increasingly implicate RBPs in diverse common polygenic diseases^1,2,15–19^, including neuropsychiatric, autoimmune and metabolic traits. These findings indicate that perturbation of RBP mediated post-transcriptional regulatory programs is an important mechanism linking genetic variation to disease susceptibility. However, as the cell-type-specific regulatory targets of RBPs have not been systematically characterized, our understanding of these critical mechanisms remains incomplete.

The advent of single-cell RNA sequencing (scRNA-seq) has enabled high-resolution, high-throughput characterization of cellular heterogeneity and dynamic cell states across complex tissues^20^. Cell-specific regulatory programs are shaped by coordinated transcriptional and post-transcriptional regulators, including transcription factors (TFs), RBPs and non-coding RNAs such as microRNAs^1,21,22^; for example, the LIN28–let-7 regulatory axis plays a key role in developmental timing and cell fate transitions during neurogenesis^23,24^. Accumulating evidence indicates that a subset of TFs, particularly zinc-finger proteins, can also bind RNA. These RNA-binding activities of TFs are largely distinct from their DNA-binding preferences and with emerging roles in post-transcriptional regulation^25,26^. RBP-target interactions are often reflected in coordinated expression patterns between RBPs and their target transcripts^27^, enabling transcriptome-based inference of regulatory activity^4,28^. Recent advances in long-read sequencing technologies now enable direct profiling of full-length transcript isoforms, capturing alternative transcription start sites, exon usage and polyadenylation. Integration with single-cell barcoding has further enabled isoform profiling with cell type, regional and developmental specificity^29–31^, thereby providing a critical foundation for dissecting cell-specific RBP-mediated regulation at isoform resolution.

Gene regulatory network (GRN) inference frameworks based on TFs have proven powerful for dissecting regulatory programs underlying cell fate specification and cellular identity^21,22,32–35^. Similar to TFs, RBP binding targets can be computationally defined based on motif specificity, which scores RNA sequences according to their likelihood of harboring RBP binding sites^8,36^. Incorporating motif-based binding models has substantially improved identification of RBP–target interactions from transcriptomic data^2,4,8^. However, available RBP motif resources remain fragmented across databases and experimental platforms, hindering the construction of unified transcriptome-wide regulatory maps. Moreover, computational frameworks for regulatory network inference remain TF-centric, and systematic identification of RBP regulatory targets at transcriptome scale—particularly at single-cell resolution—remains limited.

GWAS of complex diseases have identified thousands of risk-associated loci^37–39^, yet translating this polygenicity into downstream cellular and molecular mechanisms remains a major challenge^40^. Integrating scRNA-seq data with GWAS summary statistics has increasingly been used to resolve the core cell types and pathways underlying the polygenic architecture of complex disease at single-cell resolution^41–46^. Complementary efforts have constructed gene co-expression networks and TF-centered regulatory programs and integrated GWAS signals to identify hub genes and key transcriptional regulators underlying disease risk^29,32,35,47,48^. These advances highlight the power of integrating polygenic signals with single-cell transcriptomic profiles to capture trait-relevant cellular features. However, current approaches have largely focused on TF-to-gene transcriptional regulation and lack scalable, principled frameworks for modeling RBP-centered regulatory programs, limiting the ability to link genetic risk to disease-relevant regulatory circuitry in a cell type-specific manner.

To address these gaps, we developed scRBP (single-cell RNA-Binding Protein regulon inference), a computational method that reconstructs RBP-mediated regulatory networks at gene and isoform resolution from short- and long-read single-cell transcriptomes. scRBP implements common- and rare-variant models that separately integrate cell-specific regulon activity with common-variant GWAS signals or gene-level rare-variant association statistics to prioritize trait-relevant RBP regulons. Across simulations and 6.2 million single-cell transcriptomes, scRBP demonstrates robust and scalable performance in regulon inference and trait-relevant network prioritization. Applications across 32 tissues and 50 complex traits uncovered extensive lineage-specific RBP regulatory programs. In the developing human cortex, common- and rare-variant analyses independently converged on overlapping, developmentally dynamic neuronal RBP regulons, highlighting post-transcriptional regulation as a shared molecular layer of neuropsychiatric disease risk.

## RESULTS

### scRBP reconstructs cell-specific RBP regulons and links them to genetic risk

scRBP provides a systematic and scalable framework for reconstructing RBP-mediated gene and isoform regulatory networks (“regulons”) from single-cell transcriptomes and prioritizing regulons associated with complex disease. The framework comprises six main modules: (i) developing a comprehensive compendium of RBPs and their associated motif clusters from diverse public resources; (ii) motif-guided, transcriptome-wide ranking of candidate RBP targets at gene and isoform resolution; (iii) inference of RBP-gene and RBP-isoform co-expression networks from short- and long-read single-cell transcriptomes, respectively; (iv) motif-based refinement of RBP-target interactions to define high-confidence regulons and quantify cell type-specific regulon activity scores (RASs); (v) implementation of common- and rare-variant models that separately map common-variant GWAS signals or gene-level rare-variant association statistics to regulons to derive regulon-level genetic association scores (RGSs); and (vi) calculation of a trait-relevance score (TRS) for each regulon in each cellular context by integrating RAS with the corresponding common- or rare-variant RGS, with statistical significance assessed by Monte Carlo (MC) sampling against matched null regulons (**Fig. 1a-d**). The two genetic models are applied independently, enabling common- and rare-variant associations to be analysed separately and their convergence at the regulon level to be evaluated. Individual components are discussed further below with full details provided in **Methods**.

**Figure 1.**
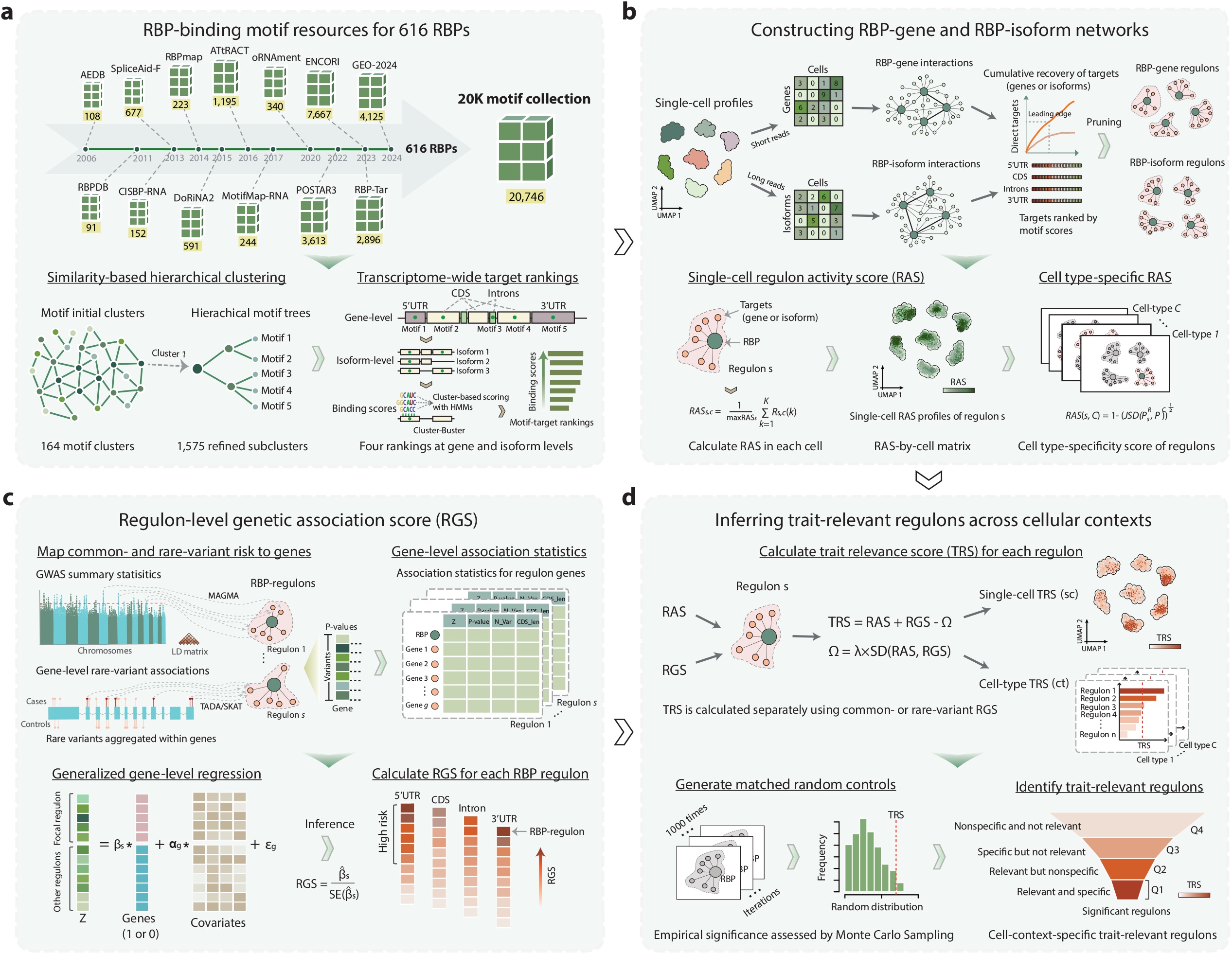
scRBP maps common- and rare-variant associations to cell-specific RBP regulons. (a) RBP-binding motif resources and transcriptome-wide target ranking. A unified compendium of 20,746 RBP-binding motifs representing 616 RBPs was assembled by integrating and harmonizing 13 distinct resources. Motifs were clustered by pairwise similarity into 164 initial clusters and further refined into 1,575 subclusters. Motif binding regions were scored using Cluster-Buster, yielding transcriptome-wide motif-target rankings across four transcript regions (5’UTR, CDS, intronic, and 3’UTR) at both gene and isoform resolution. (b) RBP regulon reconstruction and activity scoring. RBP–gene and RBP–isoform co-expression networks were inferred from short-read gene-expression profiles and long-read isoform-expression profiles, respectively. Co-expression–based RBP–target interactions were refined using transcriptome-wide motif rankings and pruned to define RBP–gene and RBP–isoform regulons. Regulon activity scores (RAS) were computed in individual cells from the expression ranks of regulon targets and subsequently summarized to quantify cell-type–specificity of each regulon. (c) Mapping common- and rare-variant associations to RBP regulons. For the common-variant model, SNP-level GWAS summary statistics were converted into linkage disequilibrium (LD)-adjusted gene-level association statistics (MAGMA). For the rare-variant model, scRBP uses precomputed gene-level association statistics from gene-based rare-variant sequencing analyses (e.g., TADA, SKAT and SKAT-O). The case–control schematic illustrates the conceptual derivation of these rare-variant statistics through the aggregation of variants within genes; variant-level association testing is not performed by scRBP. Common- and rare-variant association data were analysed using parallel, independently applied models. Generalized gene-level regression tests whether genes within each focal regulon carry stronger association signals than background genes while adjusting for model-specific gene-level covariates, including coding-sequence length in the rare-variant model. Each model independently generates a regulon-level genetic association score (RGS), defined from the standardized regulon effect, for regulons targeting the 5′UTR, CDS, intronic or 3′UTR regions. (d) Identification of trait-relevant regulons across cellular contexts. For each genetic model, the corresponding RGS is combined separately with the cell-specific RAS to calculate a trait-relevance score (TRS). A penalty term Ω downweights regulons with discordant activity and genetic association. TRSs can be calculated at single-cell or cell-type resolution. Empirical significance is assessed separately for the common- and rare-variant models by Monte Carlo sampling against 1,000 matched random regulons, generating empirical null distributions and P values. Significant regulons are classified according to their genetic relevance and cellular specificity, yielding RAS, RGS and TRS estimates for each regulon–cellular context–trait combination.

To systematically infer RBP-RNA binding interactions, we first assembled a curated compendium of 20,746 unique, high-confidence RBP-binding motifs—the “20K motif collection”—from 13 distinct resources, covering 616 RBPs with experimental evidence of RNA-binding (**Methods; Fig. 1a, Supplementary Fig. 1a-d** and **Supplementary Table 1**). Two orthogonal measures of motif quality showed strong concordance, supporting the robustness of the motif collection (r = 0.85, P < 2.2×10^−16^; **Supplementary Fig. 1e** and **Supplementary Note 1**). Because functionally related motifs often co-occur within localized genomic regions to collectively contribute to gene regulation^49^, we performed pairwise motif similarity and clustering analyses, yielding 1,575 high-resolution non-redundant motif clusters (mean 5.45 motifs per cluster; **Supplementary Fig. 1f,g**). scRBP then used cluster-based motif scanning^50^ to score and rank candidate targets across four transcript regions—the 5′ untranslated region (5′UTR), coding sequence (CDS), introns and 3′ untranslated region (3′UTR)—at both gene and isoform resolution (**Methods**). Benchmarking against ENCODE eCLIP data^8^ that scRBP consistently outperformed four alternative approaches^22^, achieving recall values of 0.848 and 0.834 at the gene and isoform levels, respectively, corresponding to improvements of 6.5% and 8.7% over the mean performance of the alternative methods (**Supplementary Fig. 2a-j**; **Supplementary Note 2**).

To reconstruct regulatory networks, scRBP used gradient boosting^51^ to infer RBP-gene or RBP-isoform interactions from short- or long-read single-cell transcriptomes, leveraging a seeded co-expression-based ‘guilt-by-association’ framework^52^ (**Fig. 1b**). Consistent with earlier studies^4,20^, benchmarking supported gradient boosting as the best-performing network-inference strategy among the methods evaluated (**Supplementary Fig. 3a-j**). The resulting co-expression edges were refined using the precomputed motif-target rankings, retaining motif-supported interactions while pruning indirect or weakly supported links to define RBP–gene and RBP–isoform regulons (**Methods**; **Fig. 1b**; **Supplementary Fig. 4a**,**b**). To enable analysis of increasingly large single-cell datasets, scRBP additionally incorporated geometric sketching subsampling^53^, which preserved transcriptional diversity and improved network-recovery accuracy relative to random downsampling with minimal additional computational cost (**Supplementary Fig. 4c,d**).

scRBP next quantified the activity of each regulon in individual cells by testing whether its targets were preferentially represented among the most highly expressed genes or isoforms (**Methods**). The resulting RAS, computed as the normalized area under the cumulative recovery curve within a predefined top-ranked window (5% by default), captures both the proportion of regulon members and their relative expression ranks within each cell (**Fig. 1b**). Because RAS is based on within-cell ranks rather than absolute expression values, it is robust to differences in expression scale and normalization. Single-cell RAS values were subsequently aggregated using an entropy-based framework to quantify the specificity of regulon activity across annotated cell types^21,22,54^. Regulon recovery remained highly consistent across varying cell counts, supporting the stability of the inferred regulatory programs (**Supplementary Fig. 5a–c**).

To quantify the genetic relevance of each regulon, scRBP implements both common- and rare-variant models that accommodate the distinct statistical properties of the two classes of genetic association data (**Fig. 1c**). In the common-variant model, SNP-level GWAS summary statistics are converted into LD-adjusted gene-level association statistics and tested for enrichment within each regulon using competitive gene-set regression, adjusting for gene size, SNP density, effective gene size and their log-transformed terms. In the rare-variant model, scRBP takes precomputed gene-level association statistics from sequencing studies and applies a separate gene-level regression framework to test whether rare-variant association signals are enriched among genes within each regulon, adjusting for coding-sequence length. Each model yields an RGS, which provides a standardized measure of genetic support for a focal regulon within the corresponding genetic analysis. For the common-variant model, sensitivity analyses across SNP-to-gene mapping windows showed stable performance, with an intermediate window of approximately 50 kb yielding the greatest separation between genetically enriched and background regulons (**Supplementary Fig. 6a**,**b**).

For each genetic model, scRBP combines cell-specific RAS with the corresponding common- or rare-variant RGS to calculate a trait-relevance score (TRS) for each regulon in a given cellular context (**Methods**; **Fig. 1d**). A regularization term penalizes discordance between regulon activity and genetic association, thereby prioritizing regulons supported by both signals over those driven predominantly by one component. The regularization parameter is optimized by cross-validation to maximize E-statistics, while avoiding over-fitting (**Supplementary Fig. 6c**,**d**). Statistical significance is assessed using MC sampling against 1,000 matched null regulons that preserve regulon size, genomic and expression properties. RAS, RGS and TRS values are standardized against their respective empirical null distributions, facilitating comparisons across traits, cell types, and genetic models.

In summary, scRBP reconstructs RBP-gene and RBP-isoform regulons, quantifies their activity across individual cells and cell types, and independently maps common- or rare-variant association signals to these networks. By integrating single-cell transcriptomes, curated RBP-binding motifs and complementary genetic association data, scRBP identified regulons that combine genetic relevance with cellular specificity and enables direct comparison of the post-transcriptional programs implicated by common and rare genetic variation.

### RBP regulons capture regulatory programs complementary to TF networks

We examined the regulatory landscape captured by scRBP and its relationship to TF-centered regulatory networks. As expected, the 616 RBPs represented in the motif compendium were strongly enriched for RNA splicing, RNA metabolism and RNA localization pathways (**Fig. 2a**; **Supplementary Table 4**). Comparison with 1,390 TFs curated in pySCENIC^21,55^ identified 66 shared proteins, representing a significant overlap between the two collections (OR = 1.59; P = 0.001, Fisher’s exact test; **Fig. 2b**). Of these shared factors, 83.3% belonged to the zinc-finger family, consistent with the capacity of many zinc-finger proteins to bind both RNA and DNA and thereby connect transcriptional and post-transcriptional regulatory layers^25,26^ (**Supplementary Table 5**).

**Figure 2.**
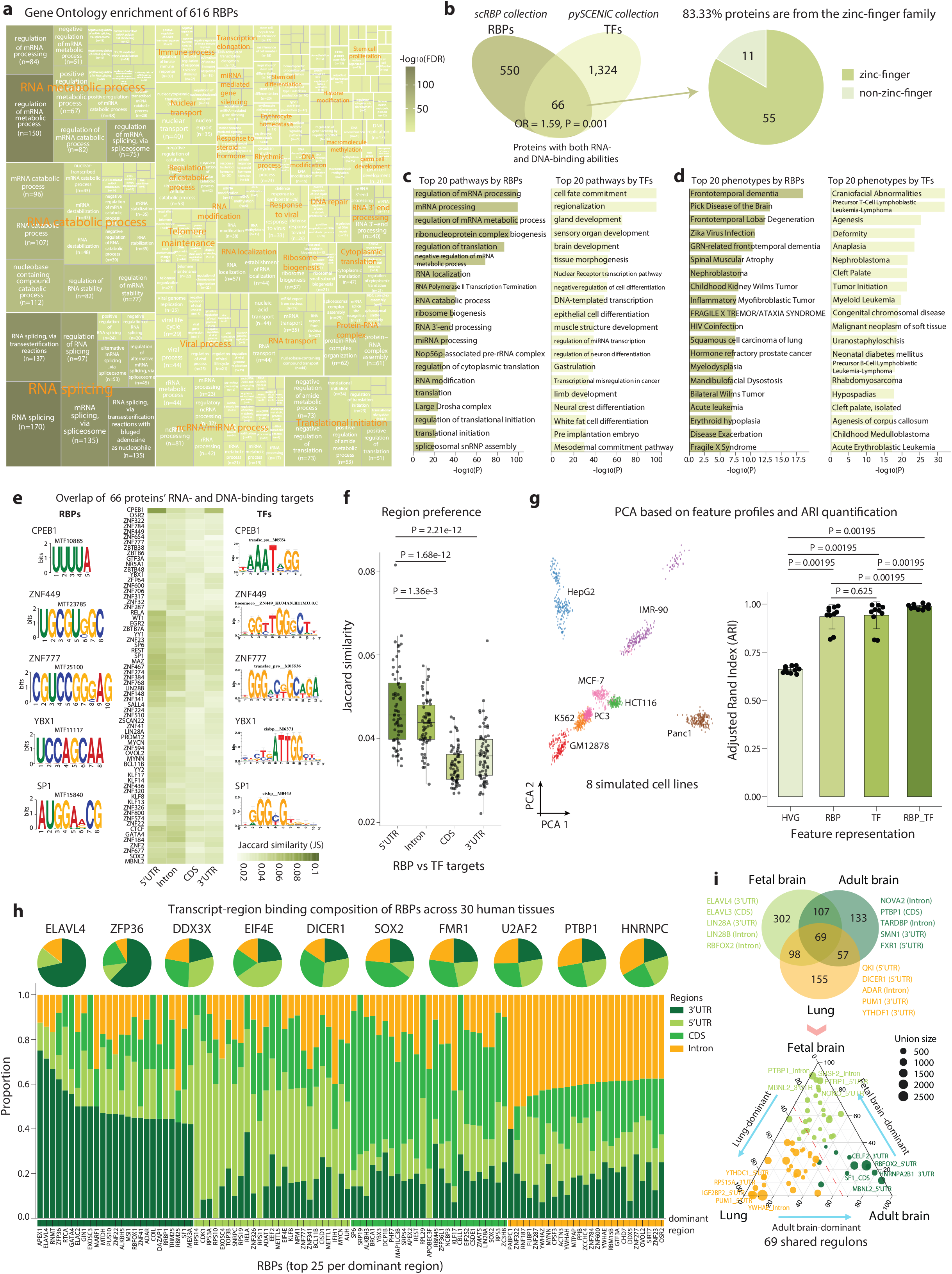
scRBP delineates RBP regulatory programs complementary to TF-centric networks. (a) Gene Ontology (GO) biological process enrichment of the 616 RBPs curated in the scRBP motif collection. Enrichment analysis was performed using *enrichGO*. (b) Overlap between curated RBPs and TFs. Left, comparison of 616 RBPs in the scRBP collection with 1,390 human TFs in the pySCENIC database identifies 66 proteins annotated with both RNA- and DNA-binding capabilities (odds ratio (OR) = 1.59, *P* = 0.001, two-sided Fisher’s exact test). Right, distribution of protein families among these 66 dual-binding proteins, showing enrichment of zinc-finger proteins. (c) Top 20 enriched pathways of 616 RBPs (left) and 1,390 TFs (right), illustrating functional divergence between post-transcriptional and transcriptional regulatory programs. Pathway enrichment analysis was performed using Metascape. (d) Top 20 enriched phenotypes of 616 RBPs (left) and 1,390 TFs (right), revealing distinct disease associations across regulatory modalities. Phenotype enrichment analysis was performed using DisGeNET. (e) Overlap between TF-derived and RBP-derived regulatory targets among the 66 dual-binding proteins. Heatmap displays Jaccard similarity between TF motif targets and RBP motif targets derived from 5′UTR, intronic, CDS, and 3′UTR motif rankings. For each protein, the top 1,500 targets were retained using minimum-rank aggregation across motifs. Representative motif logos illustrate distinct sequence features captured by TF and RBP motif models. (f) Region-specific similarity between TF and RBP targets across the 66 shared proteins. Boxplots show distributions of Jaccard similarity values per region. TF targets exhibit significantly greater similarity to 5′UTR-derived RBP targets compared to intronic, CDS, or 3′UTR-derived targets (paired Wilcoxon signed-rank test; P values indicated). (g) PCA-based visualization and ARI benchmarking of feature representations across eight deeply profiled ENCODE cell lines. Left, PCA projection of pseudocells (n = 1,600 cells) generated from simulated data (200 cells for each cell line). Right, clustering accuracy measured by the Adjusted Rand Index (ARI) for feature representations based on the top 2,000 highly variable genes (HVGs), RBP regulons, TF regulons, and combined RBP+TF regulons. Clustering was performed independently across 10 replicates for each feature group. Bars represent mean ARI values, error bars indicate standard deviation, and dots denote individual replicates (paired Wilcoxon signed-rank test; P values indicated). (h) Transcript-region binding landscape of RBPs across 30 human tissues. Stacked bar plot showing the transcript-region composition of the top 100 RBPs ranked by regulatory enrichment across four transcript regions (3′UTR, 5′UTR, coding sequence (CDS), and intronic), aggregated across tissues. For each RBP, the fraction of tissues showing significant enrichment in each transcript region was calculated based on scRBP pruning results and normalized so that regional proportions sum to one. RBPs are grouped according to their dominant transcript region. Only RBPs showing enrichment in at least five tissues were retained. Pie charts above highlight representative RBPs and illustrate their transcript-region binding compositions. (i) Tissue sharing of RBP-regulons for individual transcript regions illustrated as ternary plots. Each point represents one RBP-regulon recovered for that region; its barycentric position encodes the relative contribution of fetal brain, adult brain, and lung. Points near a vertex indicate tissue-biased RBP-regulons; points toward edges or center indicate RBPs shared by two or three tissues. Selected RBPs characteristic of each vertex are annotated. The Venn diagram above the ternary plot report overlap in RBP-regulons across the three tissues for the corresponding region; representative RBPs enriched in each tissue are listed around the diagram.

Despite this overlap, RBPs and TFs showed distinct functional and disease associations. RBPs were preferentially associated with mRNA processing and translational regulation, whereas TFs were enriched for programs related to cell fate commitment, regionalization and tissue morphogenesis (**Fig. 2c**; **Supplementary Fig. 7a**,**b**). Disease-phenotype enrichment also differed: RBPs were more strongly linked to neurodegenerative and neurodevelopmental disorders, including frontotemporal dementia and fragile X syndrome, whereas TFs exhibited greater enrichment for oncogenic and carcinogenesis-related phenotypes (**Fig. 2d**). Even among the 66 shared factors, RNA- and DNA-binding motifs and their predicted targets were largely distinct (**Fig. 2e**). Despite this overall divergence, RBP- and TF-derived target-gene sets showed modest but non-random overlap (median Jaccard similarity (JS) = 0.039; permutation P = 9.99 × 10^−4^; **Supplementary Fig. 7c**). Representative shared factors, including CPEB1 and SP1, exemplified this divergence, displaying markedly different RNA- and DNA-binding motifs (**Fig. 2e**). TF target sets showed the greatest similarity to RBP targets defined from 5′UTRs, compared with those derived from CDSs, introns or 3′UTRs (**Fig. 2f**; **Supplementary Fig. 7d**), consistent with preferential coupling between transcriptional regulation and 5′UTR-associated post-transcriptional control^25,56^.

To determine whether these regulatory representations capture cellular identity, we compared feature spaces derived from highly variable genes (HVGs), RBP regulons, TF regulons or their combination using simulated single cell profiles from eight ENCODE cell lines (**Fig. 2g**; **Supplementary Fig. 7e**). Across ten independent replicates, regulon-based representations outperformed HVG-based features in recovering cellular identity (P = 1.95 × 10^−3^). RBP regulons performed comparably to TF regulons (P = 0.625), whereas the combined RBP–TF representation showed the strongest overall performance across metrics (**Fig. 2g**). Thus, RBP regulons encode cell-state information that is complementary to, rather than redundant with, TF-centred networks.

We next characterized transcript-region preferences across 30 human tissues (**Methods**). Hierarchical clustering of region-enrichment profiles identified distinct classes of RBPs preferentially associated with 3′UTRs, 5′UTRs, CDSs or introns (**Fig. 2h**; **Supplementary Fig. 8a**; **Supplementary Table 6**). These patterns were broadly consistent with established RBP functions: ELAVL4 and ZFP36 belonged to 3′UTR-dominant class, in line with their established roles in mRNA stability and translational control, whereas HNRNPC and PTBP1 showed intron-dominant profiles consistent with their functions in RNA splicing (**Fig. 2h**; **Supplementary Fig. 8b**). Some RBPs displayed broader regional preferences. For example, although RBFOX1 is best known as a splicing regulator, it also showed substantial targeting outside introns (**Supplementary Fig. 8b**), consistent with cytoplasmic RBFOX1 binding to 3’UTRs^57,58^. Notably, 5′UTR-dominant RBPs showed the strongest overlap with OMIM-associated Mendelian disease genes, particularly genes implicated in neurodevelopmental and psychiatric disorders, consistent with recent results that constrained genes have longer 5’ UTRs and exhibit greater TSS diversity^59^ (**Supplementary Fig. 8c**).

Finally, we applied scRBP to developing human brain^60^, adult brain^61^, and lung^62^ single-cell datasets each subsampled to 100,000 cells for comparability. Hundreds of regulons were identified in each tissue, with both conserved and tissue-specific distributions across transcript regions (**Supplementary Fig. 9a, b**). Cross-region target overlap was generally limited, indicating that individual RBPs can define distinct regulatory programs according to transcript location (**Supplementary Fig. 9c**). Cross-tissue comparisons similarly revealed shared and tissue-dominant regulons across developing brain, adult brain and lung (**Figure 2i**; **Supplementary Table 7**). Collectively, these analyses establish RBP regulons as a distinct and complementary layer of cell- and tissue-resolved regulatory organization that is not captured by TF-centred networks alone.

### RBP regulons underlying hematological traits across cellular contexts

To assess the ability of scRBP to identify trait-relevant RBP regulons, we next integrated GWAS summary data for blood monocyte count (N = 563,946; **Supplementary Table 2**), a highly heritable and tractable hematopoietic trait^41,42^. We further integrated results with a ground-truth single-cell transcriptomic dataset derived from PBMCs, comprising 3,313 monocytes and 3,280 T cells (**Fig. 3a**). To ensure analytical robustness, we retained only those regulons containing ten or more genes (~60% of all inferred regulons; **Supplementary Fig. 10a**; **Supplementary Table 8**). In this context, scRBP identified ten RBP-mediated regulons—including CELF1 (3’UTR), QKI (5’UTR), and RBM47 (Intronic)—that were significantly and specifically associated with monocyte count (FDR_RAS_ < 0.05, FDR_RGS_ < 0.05 and FDR_TRS_ < 0.05; **Fig. 3b**; **Supplementary Table 9**). Notably, nine of these regulons exhibited monocyte-specific activation patterns, indicating that scRBP preferentially captures trait-relevant regulatory programs in biologically-relevant cell types (**Fig. 3b**). At single-cell resolution, these monocyte count-associated regulons displayed highly specific enrichment within monocytes (**Fig. 3c**; **Supplementary Fig. 10b**), supporting biologically coherent and cell type-resolved regulatory contributions to trait variation.

**Figure 3.**
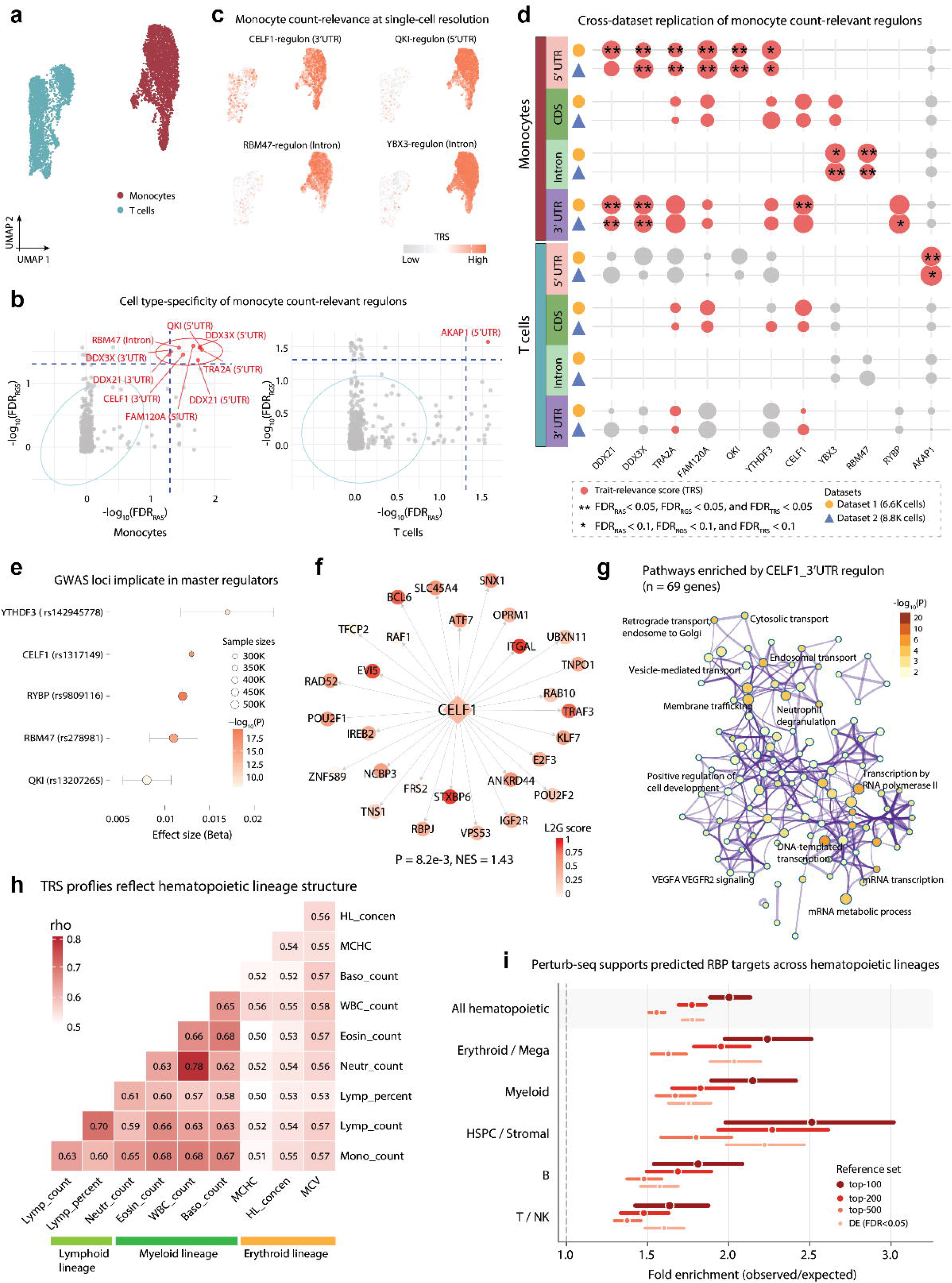
scRBP uncovers critical RBP-regulons relevant to hematological traits in a cell type-specific manner. (a) UMAP visualization of PBMC from single-cell Dataset 1 (n = 6,593 cells), highlighting the two cell types used for trait mapping: monocytes (red, n = 3,313 cells) and T cells (teal, n = 3,280 cells). (b) Scatter plots of scRBP-identified regulons associated with the monocyte count trait in monocytes (left) and T cells (right) in dataset 1. Each dot represents an RBP-regulon, with −log_10_(FDR_RGS_) on the y axis and −log_10_(FDR_RAS_) on the x axis. Gray dots denote nonsignificant regulons, and red dots indicate significant regulons. Statistical significance was assessed by an MC permutation test (one-sided, upper tail; 1,000 iterations; multiple-testing corrected). (c) Representative examples illustrating scRBP-identified monocyte count trait-relevant regulons at single-cell resolution. Four trait-relevant RBP regulons exhibit markedly stronger TRSs in monocytes compared with T cells. (d) Summary of scRBP results for the monocyte count trait across two PBMC datasets (Dataset 1: n = 6.6K cells; Dataset 2: n = 8.8K cells) and four transcript regions (5′UTR, CDS, intronic, and 3′UTR). Each dot represents an RBP regulon tested in the indicated cell type and transcript region. Dot color denotes the sign of the trait-relevance score (TRS; red, positive; gray, negative). ** indicates FDR_RAS_, FDR_RGS_, and FDR_TRS_ < 0.05; * indicates FDR < 0.1 for all three scores. Ten regulons were significantly associated with monocyte count trait. Two additional regulons, YTHDF3 (5’UTR) and YBX3 (Intronic), showed suggestive associations (FDR_RAS_, FDR_RGS_, and FDR_TRS_<0.1). (e) Forest plot of master regulators of identified RBP-regulons harboring monocyte count-associated variants within their gene loci. Points indicate GWAS effect size (beta) with 95% confidence intervals (CIs); point color corresponds to –log_10_(P-value), and point size reflects GWAS sample size. (f) Network representation of the monocyte-enriched CELF1_3′UTR regulon (n = 69 genes) genetically prioritized for monocyte count. The regulator is shown as a diamond and target genes as circles; node color encodes Open Targets L2G score. The GSEA significance for this regulon is indicated. (g) Term–term similarity network for pathways enriched among CELF1_3′UTR targets (Metascape). Nodes denote enriched terms and edges connect terms with high overlap; node color encodes −log_10_(P), and densely connected modules represent functional clusters. (h) Pairwise Spearman correlations of cell type-resolved regulon TRS profiles across ten blood-cell traits, revealing greater concordance among traits within the same hematopoietic lineage. (i) Enrichment of scRBP-inferred hematopoietic regulon targets among genes responsive to cognate RBP perturbation in a genome-scale K562 CRISPRi Perturb-seq screen. Results are shown for 1,702 regulons spanning 161 RBPs, pooled across all hematopoietic lineages or stratified by lineage, using four definitions of perturbation-responsive genes. Points and bars indicate pooled observed-to-expected enrichment and 95% bootstrap confidence intervals, respectively; the dashed line indicates no enrichment. Targets were enriched across all lineages and response thresholds (1.37–2.51-fold; permutation FDR < 2 × 10^−4^).

We next evaluated the reproducibility of these findings in an independent PBMC dataset (n = 3,546 monocytes and 5,231 T cells; **Fig. 3d**; **Supplementary Fig. 10c**). Eight of the nine monocyte-specific regulons remained significantly associated with monocyte count after FDR correction, whereas the T cell-specific regulon showed nominal significance (**Fig. 3d**; **Supplementary Fig. 10d-f**; **Supplementary Table 10**). Within each cell type, regulon-level TRSs were highly concordant between datasets (r = 0.83, P < 2.2×10^−16^; **Supplementary Fig. 11a**). Sensitivity analyses further localized these signals predominantly to CD14+ monocytes, with weaker enrichment in CD16+ monocytes and minimal activity in CD4+ or CD8+ T cells (**Supplementary Fig. 11b-d**). Cross-dataset TRS concordance reflected cell-subtype relatedness, remaining high between matched and closely related subtypes within the same lineage but decreasing markedly across lineages (**Supplementary Fig. 11e**).

To link these associations with genetics, we assessed whether master regulators of these significant regulons colocalize with monocyte count-associated GWAS loci. Notably, nearly half of these RBPs harbored genome-wide significant (GWS) associations with monocyte count (P < 5×10^−8^; **Fig. 3e**; **Supplementary Table 11**), including *CELF1, QKI, RBM47, YTHDF3*, and *RYBP*. Furthermore, we characterized regulon enrichment using Open Targets locus-to-gene (L2G) scores^63^, for monocyte count GWAS, scRBP-inferred regulons showed significant enrichment for L2G prioritized genes, with CELF1 (3’UTR), QKI (5’UTR), and RBM47 (Intronic) among the strongest signals (**Supplementary Fig. 12a-g**; **Supplementary Tables 12-14**). Concordantly, target genes within these regulons also exhibited significant L2G support (**Fig. 3f**; **Supplementary Fig. 12c**,**d**). Functionally, *CELF1* has been implicated in inflammatory responses in monocyte-derived macrophages^64^, and the CELF1 (3’UTR) regulon was enriched for cytosolic and endosomal trafficking as well as post-transcriptional gene silencing pathways (**Fig. 3g**; **Supplementary Fig. 12e**; **Supplementary Table S12**). *QKI*, an inflammation-sensitive alternative splicing regulator during monocyte-to-macrophage differentiation^65^, showed enrichment of 5’UTR-linked targets in cell-cell signaling and metabolic processes (**Supplementary Fig. 12f**; **Supplementary Table 13**).

To assess the broader generalizability of scRBP, we extended the analysis to GWAS of nine additional blood cell traits (**Supplementary Table 2**). scRBP identified cell-type-specific trait-relevant regulons (**Supplementary Fig. 13a**), with highly concordant trait-regulon association profiles among cell populations within the same hematopoietic lineage (**Fig. 3h**), consistent with established lineage-specific genetic architectures of blood traits^41,42^. Notably, 67.61% of trait-relevant regulons were associated with multiple hematological traits (**Supplementary Fig. 13b**), consistent with widespread genetic pleiotropy across blood cell phenotypes^66^, and these associations were supported by L2G enrichment (**Supplementary Fig. 13c**). Convergence was also evident at the regulator level: 68.97% of RBPs in trait-relevant regulons harbored GWS variants associated with at least two hematological traits (**Supplementary Fig. 14a-b**). For example, *CELF1* was associated not only with monocyte count but also with neutrophil and eosinophil counts.

To determine whether scRBP regulons capture functional RBP–target relationships beyond co-expression, we evaluated lineage-resolved hematopoietic regulons against a genome-scale CRISPRi Perturb-seq screen in K562 cells^67^. After matching perturbed RBPs to the hematopoietic regulon atlas, 1,702 lineage- and cell-type-resolved regulon instances derived from five lineage-specific scRBP analyses were retained for evaluation (**Supplementary Table 15**). Predicted targets were enriched among genes responsive to cognate RBP perturbation relative to all measured genes (1.4–2.5-fold; FDR < 2 × 10^−4^; **Fig. 3i**; **Supplementary Table 16**). This enrichment persisted when cognate perturbations were compared with non-cognate RBP perturbations (1.1–1.3-fold; **Supplementary Fig. 14c**), supporting RBP-specific effects beyond generic perturbation responses. At single-regulon resolution, cognate RBP perturbations ranked significantly higher than expected by chance (P = 9 × 10^−14^), with several established and disease-associated RBPs, including DDX3X and TARDBP, showing strong cognate specificity (**Supplementary Fig. 14d-f**). Together, these analyses reveal a reproducible and lineage-resolved RBP regulatory architecture associated with hematological traits, supported by convergent evidence from human genetics and functional perturbation.

### Dissecting cell type-specific RBP regulons underlying autoimmune diseases

We further integrated 172,601 cells spanning 13 immune cell types with GWAS data for eight autoimmune diseases (**Fig. 4a**; **Supplementary Table S2**). We identified 44 distinct RBP regulons showing cell-type-specific trait associations, with rheumatoid arthritis (RA) showing the largest number of associated regulons, followed by multiple sclerosis (MS) and type 1 diabetes (T1D; **Fig. 4b**; **Supplementary Table 17**). Associations were concentrated in myeloid populations, particularly CD14+ monocytes, CD16+ monocytes and myeloid dendritic cells (mDCs), and included MAP1LC3B 5’UTR- and CDS-associated regulons and DDX3X 3’UTR-, CDS- and intron-associated regulons (**Fig. 4c**; **Supplementary Table 18**). This myeloid predominance is consistent with the established contribution of innate immune programs to autoimmune genetic risk^68,69^. Several regulons were recurrently associated across lymphoid populations, including HNRNPUL1 (3’UTR), RBM17 (3’UTR), SSB (CDS), and SF3B1 (5’UTR), suggesting shared post-transcriptional programs across related lymphoid cell states (**Fig. 4b**). At single-cell resolution, representative trait-associated regulons, including MAP1LC3B (5′UTR), ZFP36L1 (3′UTR), RPS15A (5′UTR) and HNRNPUL1 (3′UTR), showed preferential activity in cells with high trait-relevance within their corresponding lineages (**Fig. 4d**; **Supplementary Fig. 15a-b**).

**Figure 4.**
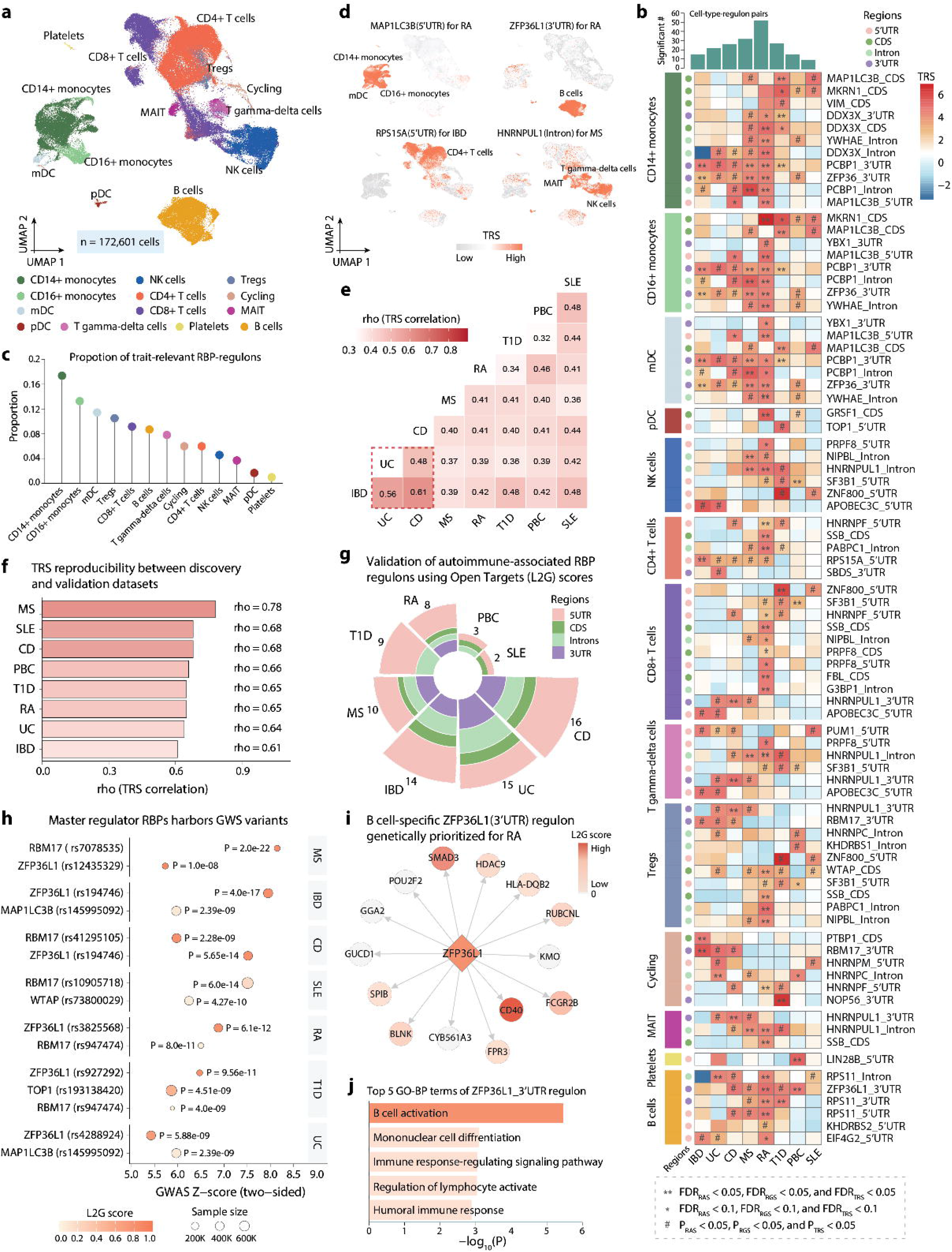
scRBP identifies immune cell type-specific regulons relevant to autoimmune diseases. (a) UMAP visualization of the PBMC single-cell transcriptome, colored by annotated immune cell types (n = 172,601 cells). This dataset includes CD14+ monocytes (n = 23,100), CD16+ monocytes (n = 5,537), myeloid dendritic cells (mDC; n = 1,417), plasmacytoid dendritic cells (pDC; n = 734), CD8+ T cells (n = 34,110), CD4+ T cells (n = 49,415), regulatory T cells (Tregs; n = 3,251), cycling (n = 1,012), T gamma-delta cells (n = 3,183), mucosal-associated invariant T cells (MAIT; n = 2,213), natural killer (NK) cells (n = 21,871), B cells (n = 26,132), and platelets (n = 626). (b) Heatmap summarizing scRBP-identified cell type-specific RBP regulons associated with eight autoimmune diseases across 13 immune cell types. ** indicates FDR_RAS_, FDR_RGS_, and FDR_TRS_ < 0.05; * indicates FDR < 0.1 for all three scores; ^#^ indicates nominal significance (P_RAS_, P_RGS_, and P_TRS_ < 0.05). Statistical significance was assessed by MC permutation testing (one-sided, upper tail; 1,000 iterations; multiple-testing corrected). Region labels denote transcript regions (5′UTR, CDS, intronic, and 3′UTR). (c) Proportion of autoimmune disease-associated RBP-regulons in each immune cell type. (d) Representative examples illustrating the cell type specificity of scRBP-identified regulons associated with autoimmune diseases. Four disease-associated RBP regulons show markedly stronger TRSs in their focal cell types compared with other immune populations. (e) Heatmap showing correlations of cell-type-resolved regulon TRSs across eight autoimmune diseases. The boxed region highlights strong concordance of trait relevance among IBD, UC, and CD. (f) Cell type-resolved regulon TRS correlations between discovery and validation datasets across eight autoimmune diseases. scRBP results derived from the discovery dataset were evaluated in an independent validation dataset (n = 10,393 cells). Pearson correlation coefficients are shown. (g) Radial stacked bar plot showing the number of autoimmune-associated RBP regulons across diseases, stratified by transcript regions (5’UTR, CDS, intronic, and 3’UTR). (h) Genome-wide significant (GWS) variants mapped to RBPs across autoimmune diseases. Each point represents a GWS variant-RBP association, plotted by the absolute GWAS Z score inferred from two-sided p values. Point color indicates the L2G score, and point size reflects GWAS sample size. P values for each association are shown; disease labels are indicated on the right. (i) Representative network of the B cell-specific ZFP36L1 (3’UTR) regulon genetically prioritized for RA. The diamond node denotes the regulator ZFP36L1, and circle nodes represent target genes within the regulon. Node color indicates L2G scores from Open Targets, reflecting the degree of genetic prioritization for RA. (j) Functional enrichment analysis of the ZFP36L1(3′UTR) regulon. Bar plot showing the top five enriched GO-BP terms, ranked by enrichment significance, as assessed by WebGestlat.

Trait-relevant regulons were extensively shared across autoimmune disorders, with 65.91% associated with at least two diseases (**Supplementary Fig. 15c**), consistent with pervasive genetic pleiotropy^35,70^. Several myeloid-enriched regulons, including MAP1LC3B (CDS), DDX3X (Intronic, 3’UTR and CDS), PCBP1 (3’UTR and Intronic), and ZFP36 (3’UTR), were associated with three or more autoimmune diseases, with comparable pleiotropy observed among lymphoid-enriched regulons such as RBM17 (3’UTR), RPS15A (5’UTR), HNRNPUL1 (3’UTR and Intronic), and WTAP (CDS) (**Supplementary Fig. 15d**). At the cell-type-level, regulon TRS profiles were highly concordant across related autoimmune diseases, with particularly strong similarity among inflammatory bowel disease (IBD), ulcerative colitis (UC), and Crohn’s disease (CD) (**Fig. 4e**).

To assess reproducibility, we evaluated scRBP in an independent PBMC single-cell dataset (**Supplementary Fig. 16a**). Trait-relevant regulons showed concordant associations in matched cell types (**Supplementary Fig. 16b**; **Supplementary Table 19**), and cell type-resolved TRS profiles were strongly correlated between discovery and validation datasets across all eight autoimmune disorders (ρ = 0.61-0.78; **Fig. 4f**; **Supplementary Fig. 16c**; **Supplementary Table 20**). Orthogonal analysis using L2G prioritization further supported these associations, with targets of scRBP-prioritized regulons significantly enriched among genetically prioritized genes across all eight diseases (**Fig. 4g**; **Supplementary Fig. 17a**). Regulons including ZFP36L1 (3’UTR), DDX3X (3’UTR), MAP1LC3B (CDS) and RPS15A (5’UTR) were recurrently implicated across multiple autoimmune diseases, reinforcing the presence of shared pleiotropic RBP programs (**Supplementary Fig. 17a**).

Five master-regulator RBPs—*ZFP36L1, RBM17, MAP1LC3B, WTAP* and *TOP1*—were located at GWS loci for one or more autoimmune diseases (**Fig. 4h**; **Supplementary Table 21**). *ZFP36L1* showed GWAS associations with five diseases and encodes a zinc-finger RNA-binding protein required for maintenance of the marginal-zone B cell compartment^71^. Consistent with previous associations with MS^72^ and RA^16,73^, scRBP identified a B-cell-specific ZFP36L1 (3’UTR) regulon associated with five autoimmune diseases, including MS and RA (**Fig. 4b**). Targets of this regulon showed strong L2G prioritization for RA and other autoimmune diseases (**Fig. 4i**; **Supplementary Fig. 17b**) and were enriched for pathways related to B cell activation and immune signaling (**Fig. 4j**). Similarly, *RBM17, MAP1LC3B, WTAP*, and *TOP1*, each previously implicated in autoimmunity^17,18,74,75^, defined genetically supported, cell-type-specific regulons enriched for inflammatory and immunity-related pathways (**Supplementary Fig. 17c-f**). Collectively, these findings link autoimmune genetic risk to lineage-specific post-transcriptional programs in immune cells.

### RBP regulons links genetic risk to metabolic and cardiovascular traits

We next integrated liver and vascular single-cell datasets with GWAS data for nine metabolic traits and six cardiovascular diseases, respectively (**Supplementary Fig. 18a, 19a**; **Supplementary Tables 2-3**). scRBP identified cell type-resolved trait-relevant regulons in both tissues (**Supplementary Fig. 18b-d, 19b-d**; **Supplementary Tables 22-23**). In the liver, hepatocyte-enriched regulons were prominently associated with lipid-related traits, including high density lipoprotein (HDL) and triglycerides (TG). In vascular tissue, regulons in endothelial cells, pericytes and macrophages were associated with hypertension (HT), coronary artery disease (CAD) and atrial fibrillation (AF). Many regulons were shared across related phenotypes while retaining marked cell-type specificity. At the regulator level, 31 RBPs mapped to GWS loci for metabolic traits, including *ZNF706* for HDL, *TOP1* for LDL and *FTO* for TG (**Supplementary Fig. 18e**; **Supplementary Table 24**), whereas 17 RBPs were located at cardiovascular disease loci, including *SP1* for HT, *CPEB4* for CAD and *XRN2* for AF (**Supplementary Fig. 19e**; **Supplementary Table 25**). Together, these findings show that trait-associated RBP regulatory architecture extends across tissues while retaining distinct cellular specificity for metabolic and cardiovascular phenotypes.

### Mapping cell-specific trajectories of RBP regulons across human brain development

Post-transcriptional regulation is central to human cortical development, enabling rapid, cell-type-specific control of RNA processing and protein output during progenitor expansion, neuronal differentiation and migration^6,7,29^. Yet unlike TF-mediated programs^34,35,60,76^, the cell-type-resolved architecture of RBP regulation across human neurodevelopment remains poorly defined. We therefore applied scRBP to a developing human cortical atlas^60^ comprising 599,221 cells from 96 individuals spanning 6 weeks post-conception to 8 months postnatal (**Fig. 5a**; **Supplementary Fig. 20a-e**; **Supplementary Table 26**). Across 14 major cell types, scRBP identified 576 RBP regulons containing at least ten target genes, revealing distinct lineage-structured programs across neuronal, progenitor, glial, and neurovascular populations (**Fig. 5b**; **Supplementary Fig. 20f**; **Supplementary Table 27**). Most regulons were preferentially active in neuronal lineages, whereas a distinct subset was enriched in progenitor and glial populations (**Supplementary Fig. 20g**). These patterns recapitulated established post-transcriptional biology: ELAVL- and RBFOX-family regulons showed strong neuron-specific activity, consistent with their roles in neuronal RNA processing and fate specification^6,7,77,78^, whereas a TARDBP-associated regulon was preferentially active in progenitor and glial populations, consistent with its predominant functions in glial biology^79,80^.

**Figure 5.**
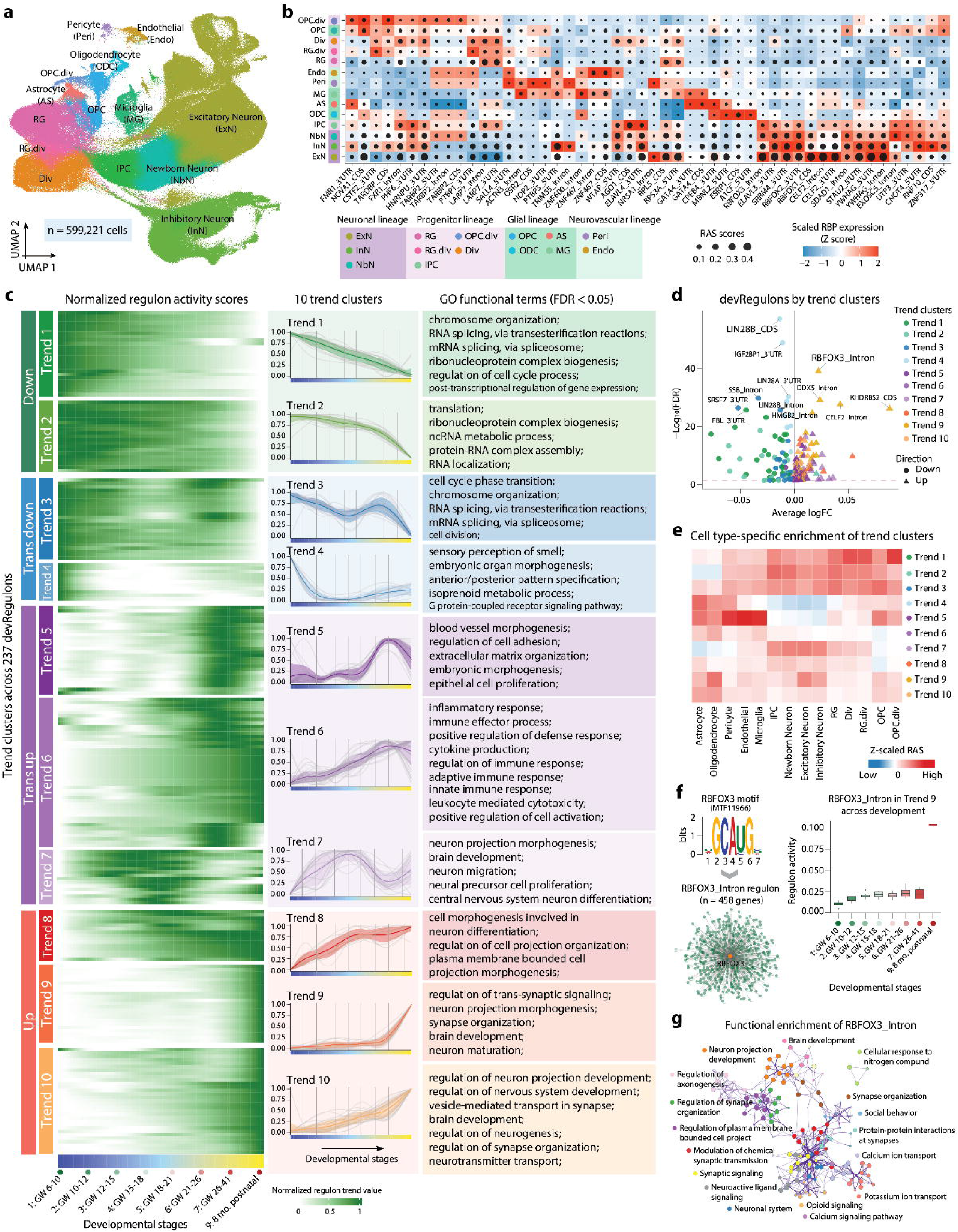
scRBP reveals cell type-specific and developmental dynamics of RBP regulons across human cortical development. (a) UMAP visualization of an integrated single-cell transcriptomic meta-atlas of the developing human cortex, colored by 14 annotated immune cell types (n = 599,221 cells). The integrated atlas captured 14 main cell populations, including excitatory neuron (ExN), inhibitory neuron (InN), newborn neuron (NbN), intermediate progenitor cell (IPC), radial glia (RG), oligodendrocyte lineage cells (OPC and ODC), astrocytes (AS), microglia (MG), dividing populations (Div, RG.div, OPC.div), and endothelial and perivascular cells. (b) Heatmap/dotplot showing RBP expression of the regulon on a color scale and cell type-specific RAS of the regulon on a size scale. Cell types are ordered based on their RBP expression similarity. (c) Developmental dynamics of RBP-regulons across eight developmental stages. Differential regulon activity analysis across developmental stages identified 237 devRegulons (FDR < 0.05), which were grouped into ten clusters (“regulon trends”). These trends form four archetypal developmental patterns: early downregulated (Trends 1–2), transiently downregulated (Trends 3–4), transiently upregulated (Trends 5–7), and progressively upregulated (Trends 8–10). Faint lines show the individual devRegulons, and bold line is the average across all devRegulons in the trend. Right panel: Functional enrichment analysis of each regulon trend cluster. Selected enriched GO terms are shown (FDR < 0.05). (d) Volcano plot showing differential activity of devRegulons across developmental stages. Each point represents a devRegulon, colored by its assigned temporal trend cluster (Trends 1–10). The x-axis indicates average log fold-change (logFC) across developmental stages, and the y-axis shows statistical significance (−log_10_FDR). Upregulated regulons are indicated by triangles, whereas downregulated regulons are shown as circles. The dashed horizontal line denotes the FDR significance threshold. Selected representative devRegulons with strong statistical support are highlighted. (e) Cell type–resolved enrichment of developmental regulon trend clusters. Enrichment was quantified by averaging RAS within each trend cluster for each cell type and normalizing values as z-scores across clusters within each cell type. (f) RBFOX3 (Intronic) regulon motif, target network, and developmental activity. Sequence logo of the canonical UGCAUG-centered RBFOX3 motif and network representation of its intronic regulon (458 predicted targets). Boxplots show regulon activity across developmental stages for the RBFOX3 (Intronic) regulon (Trend 9). (g) Functional enrichment analysis of the RBFOX3 (Intronic) regulon. The pathway enrichment analysis was performed by using Metascape.

Beyond cell type specificity, regulatory networks are dynamically remodelled across development. We therefore quantified RBP regulon activity across developmental stages using donor-level pseudobulk profiles and *limma* models with empirical Bayes moderation^78^ (**Methods**). We identified 237 developmentally dynamic regulons (“devRegulons”) that differed significantly between at least two stages (FDR < 0.05; **Supplementary Fig. 21a**; **Supplementary Table 28**). Hierarchical clustering resolved these devRegulons into ten temporal trajectories encompassing four archetypal patterns: early downregulation (Trends 1–2), transient downregulation (Trends 3–4), transient upregulation (Trends 5–7), and progressive upregulation (Trends 8–10) (**Fig. 5c; Supplementary Fig. 21b**; **Supplementary Table 29**). Early downregulated trajectories were most active during early gestation and were enriched for mRNA splicing, ribonucleoprotein complex biogenesis and cell cycle regulation (**Fig. 5c**; **Supplementary Table 30**), consistent with proliferative progenitor programs^76,81^. Trend 4 additionally captured anterior–posterior patterning and early morphogenesis. By contrast, transiently upregulated trajectories reflected more lineage-restricted programs: Trend 5 was enriched for vascular morphogenesis, endothelial remodeling and tissue-organization; Trend 6 for immune and inflammatory responses, including cytokine signalling, potentially reflecting microglial processes; and Trend 7 for neuronal migration, neural precursor proliferation and projection morphogenesis, consistent with active cortical lamination^76,78^. Progressively upregulated trajectories were enriched for neurite outgrowth, synapse organization and neurotransmission, consistent with synaptic refinement and circuit maturation^76,81^. Several trajectories, particularly Trends 3, 4 and 9 included RBPs with established roles in cortical development (**Fig. 5d**; **Supplementary Fig. 21c**).

To delineate the cellular origins of developmental regulon dynamics, we projected these global temporal trajectories onto each of the 14 major developing brain cell types (**Methods**). Across lineages, regulon activity was relatively stable during early gestation but increased coordinately toward late gestation and the perinatal period, with lineage-specific variation in the timing and magnitude of this rise (**Supplementary Fig. 21d**), consistent with heightened regulatory dynamics around birth^78^. Cell-type-resolved decomposition revealed both broadly shared and lineage-biased temporal programs (**Fig. 5e**; **Supplementary Fig. 21e**). Early declining trajectories (Trends 1–3) were broadly distributed across cell types, whereas transient programs showed greater lineage specificity: Trend 7 peaked in neuronal populations during mid-gestational neurogenesis and migration, while Trend 5 was enriched in endothelial cells and pericytes. Late development was dominated by progressively upregulated trajectories (Trends 8–10), particularly Trend 9, which showed the strongest late-stage increase and neuronal enrichment (**Fig. 5c,e**). Cell subtype-level analysis further revealed both shared and divergent temporal dynamics among closely related neuronal and glial populations (**Supplementary Fig. 22a–c**).

These analyses defined ten temporal regulon trajectories across cortical development and prioritized candidate post-transcriptional regulators of corticogenesis, including RBFOX3, LIN28B and HMGB2 (**Fig. 5f**; **Supplementary Fig. 23a-d**). The ExN-specific RBFOX3 (Intronic) regulon emerged as a prominent late-activated program within Trend 9, consistent with RBFOX3 being widely used as the NeuN marker of postmitotic neurons^82^ (**Fig. 5f**). Its predicted targets were enriched for the consensus UGCAUG binding motif and for pathways related to neuronal projection development, synapse organization and synaptic signalling (**Fig. 5g**), consistent with established roles of RBFOX proteins in neuronal splicing and circuit formation^6,7,57,58^. By contrast, the Trend 4 LIN28B (CDS) regulon was most active in radial glia and enriched for axon guidance, forebrain development and neuronal differentiation (**Fig. 5d**; **Supplementary Fig. 24a**). Its targets were also enriched for predicted let-7 microRNA targets (OR = 2.4, P < 0.017; **Methods**; **Supplementary Fig. 24b-d**; **Supplementary Table 31**), recapitulating the established LIN28–let-7 axis in neurodevelopment^23,24^. Overall, these findings reveal temporally structured RBP regulatory programs aligned with major milestones of human cortical development.

### Identifying cell type-resolved RBP regulons relevant to neuropsychiatric disorders

Genetic risk for neuropsychiatric disorders is developmentally patterned, often implicating discrete windows of early brain development^76,78^. Yet the convergent cell types and molecular programs through which polygenic risk for these disorders acts during early brain development remain poorly defined. We therefore used scRBP to integrate the developing human brain single-cell atlas with GWAS summary statistics for eight neuropsychiatric disorders (NPDs), including schizophrenia (SCZ), bipolar disorder (BIP), attention deficit hyperactivity disorder (ADHD), autism (ASD), major depressive disorder (MDD), obsessive compulsive disorder (OCD), Tourette syndrome (TS), and anorexia nervosa (AN) (**Supplementary Table S2**).

We identified 97 regulons associated with at least one neuropsychiatric disorder (**Fig. 6a**; **Supplementary Table 32**). Associations were concentrated in neuronal and progenitor populations, particularly ExN, NbN, IPC and InN (**Fig. 6b**; **Supplementary Table 33**). SCZ yielded the largest number of cell-type-specific regulons, followed by BIP, ASD, and ADHD (**Fig. 6c**). UTR-targeting regulons accounted for 55 of the 97 associations (56.7%; **Fig. 6c**; **Supplementary Table 34**), underscoring the contribution of post-transcriptional regulation to neuropsychiatric disease biology^10^. Regulon associations were frequently shared across related cell types within the same lineage: RBFOX2 5′UTR and intronic regulons were associated with SCZ in both ExN and InN, SRRM4 3′UTR with BIP in ExN and InN, and DHX9 CDS with ASD across dividing progenitor populations, including Div, RG.div and OPC.div (**Supplementary Fig. 25a-h**). At single-cell resolution, these regulons were preferentially active in cells with elevated TRSs within the corresponding populations (**Supplementary Fig. 26a**). As a negative control, we analysed Alzheimer’s disease (AD) and Parkinson’s disease (PD), two predominantly late-onset neurodegenerative disorders whose major pathogenic processes are thought to manifest in ageing brain contexts. In contrast to the robust associations observed for neuropsychiatric disorders, we detected no regulons that remained significant for AD or PD after multiple-testing correction (FDR_RAS_, FDR_RGS_, and FDR_TRS_ < 0.05 or 0.1; **Supplementary Fig. 26b**), supporting the developmental specificity of psychiatric disorder-relevant post-transcriptional programs.

**Figure 6.**
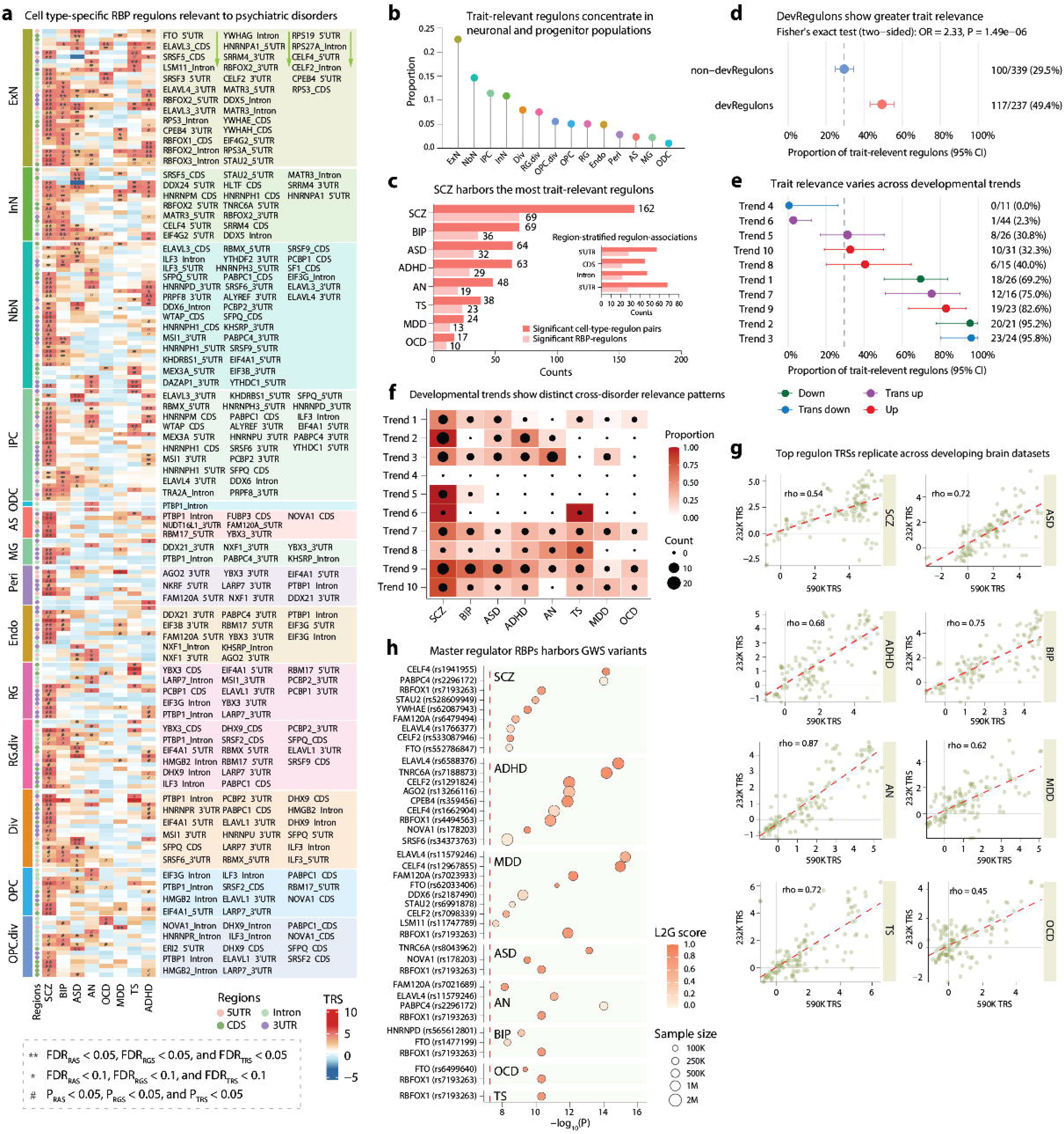
scRBP distinguishes brain cell type-resolved RBP regulons relevant to neuropsychiatric disorders. (a) Heatmap summarizing scRBP-identified cell type-specific RBP regulons associated with eight neuropsychiatric disorders across 14 brain cell types. ** indicates FDR_RAS_, FDR_RGS_, and FDR_TRS_ < 0.05; * indicates FDR < 0.1 for all three scores; ^#^ indicates nominal significance (P_RAS_, P_RGS_, and P_TRS_ < 0.05). Statistical significance was assessed by MC permutation testing (one-sided, upper tail; 1,000 iterations; multiple-testing corrected). Region labels denote associated transcript regions (5′UTR, CDS, intronic, and 3′UTR). Labels shown to the right of the heatmap denote representative regulons for the corresponding heatmap rows and should be read column-wise, from top to bottom within each column, proceeding from the first column to the second and then the third. (b) Proportion of neuropsychiatric disorder-associated RBP-regulons in each brain cell type. (c) Trait-relevant regulons across eight neuropsychiatric disorders (NPDs). Bar plots indicate the number of significant RBP regulons identified for each neuropsychiatric disorder, with the inset showing their distribution by RBP binding region (5′UTR, CDS, intronic, and 3′UTR). (d) Comparison of developmental regulons (devRegulons) versus non-developmental regulons (non-devRegulons) in neuropsychiatric trait associations. Points indicate odds ratios from Fisher’s exact tests with 95% confidence intervals. (e) Proportion of trait-relevant regulons stratified by developmental trend clusters. Points indicate estimated proportions with 95% confidence intervals for each temporal trajectory cluster. (f) Heatmap depicts the relative enrichment of trait-relevant regulons across developmental trajectory clusters for each neuropsychiatric disorder. Color denotes the proportion of associated regulon–cell-type pairs, and dot size indicates their counts. Proportions were calculated as the fraction of regulon–cell-type pairs within each trend significantly associated with the corresponding disorder. (g) Cell type–resolved regulon TRS correlations between discovery and validation developmental brain datasets across eight neuropsychiatric disorders. TRS values inferred from the discovery dataset (590K cells) were compared with those from an independent validation dataset (232K cells). Pearson correlation coefficients are shown. (h) Master regulators of scRBP-identified regulons harbor genome-wide significant (GWS) loci. Points represent RBPs harboring GWS variants across eight neuropsychiatric disorders. The x-axis shows −log_10_(P) values for variant associations; color indicates L2G scores and point size reflects GWAS sample size.

Neuropsychiatric disorder-relevant regulons showed pervasive pleiotropy, with 79.4% associated with at least two disorders (**Supplementary Fig. 27a**). The ELAVL4 (3’UTR), SRRM4 (3’UTR) and CELF2 (3’UTR) regulons were each associated with five disorders (**Supplementary Fig. 27b**). Mirroring LDSC-estimated genetic correlation^39^ (**Supplementary Fig. 27c**), cell type-resolved regulon TRS profiles were highly concordant across disorders (**Supplementary Fig. 27d**). TRS-based clustering prominently grouped SCZ with BIP, recapitulating their well-established genetic relationship (**Supplementary Fig. 27e**). Their TRS profiles were particularly concordant across neuronal and progenitor populations, with the strongest correlation observed in ExN (**Supplementary Fig. 27f**). These results indicate shared post-transcriptional regulatory programs as a pervasive component of polygenic risk across neuropsychiatric disorders.

We next examined whether developmentally dynamic RBP programs were preferentially implicated in neuropsychiatric risk. DevRegulons were more frequently trait-relevant than non-devRegulons across the eight disorders (49.4% versus 29.5%; Fisher’s exact test, OR = 2.33, P = 1.49 × 10^−6^; **Fig. 6d**; **Supplementary Table 35**). This enrichment varied substantially across temporal trajectories, peaking in early declining programs (Trends 1–3), the mid-gestational Trend 7 and the late-upregulated Trend 9 (**Fig. 6e**). DevRegulons also comprised a larger fraction of trait-relevant programs for most individual disorders, particularly SCZ, ADHD and ASD (**Supplementary Fig. 27g**). Trend-stratified analyses further revealed distinct patterns of disease coupling: mid-to-late developmental trajectories (Trends 7–10) showed broad enrichment across disorders, whereas early transient programs (Trends 1–3) were more selectively associated with individual conditions (**Fig. 6f**; **Supplementary Fig. 27h**).

To assess reproducibility, we evaluated scRBP in an independent developing human neocortex dataset comprising 232,328 cells^76^ (**Supplementary Fig. 28a**; **Supplementary Table 36**). Most disorder-associated regulons identified in the discovery dataset showed concordant support in matched cell types at significant or suggestive thresholds (**Supplementary Fig. 28b**; **Supplementary Table 37**). TRS estimates were moderately to strongly correlated between discovery and validation datasets across all eight disorders, both among top-ranked regulons (ρ = 0.45–0.87; **Fig. 6g**) and across the full regulon set (ρ = 0.51–0.63; **Supplementary Fig. 28c**). Orthogonal gene-set enrichment analysis using disorder-specific L2G gene rankings further supported these associations, with particularly strong evidence for ASD, ADHD, MDD and SCZ (**Supplementary Fig. 29a**,**b**). Several leading regulons, including RBFOX1 (CDS), RBFOX2 (5′UTR), CPEB4 (5′UTR), CELF2 (3′UTR) and RBFOX3 (intronic), were enriched across seven or more disorders (**Supplementary Fig. 29c**), reinforcing the convergence of neuropsychiatric genetic risk on shared post-transcriptional programs.

To further prioritize trait-associated regulons, we examined whether their regulator RBPs were located at GWAS loci (P < 5 × 10^−8^). 17 RBPs from significant regulons map to loci associated with at least one neuropsychiatric disorder, providing additional regulator-level genetic support (**Fig. 6h**). Examples included *RBFOX1, CELF4* and *CELF2* for SCZ; *ELAVL4, TNRC6A* and *CPEB4* for ADHD; and *TNRC6A, NOVA1* and *RBFOX1* for ASD (**Supplementary Table 38**). 12 of these RBPs, including *RBFOX1, CELF4* and *CELF2*, were also catalogued in SFARI Gene based on human genetic evidence for ASD^83^ (**Supplementary Table 39**). Collectively, these findings identify cell type- and developmentally resolved RBP regulons as a genetically supported layer of post-transcriptional regulation underlying neuropsychiatric risk.

### Rare-variant risk in ASD and SCZ converges on excitatory-neuron RBP regulons

Because common and rare variation may implicate overlapping but distinct genes and cellular contexts, we extended scRBP to rare-variant association data for ASD and SCZ, two neuropsychiatric disorders with large-scale exome-sequencing resources^84,85^ (**Supplementary Table S40**). These statistics were integrated with cell type-resolved RBP-regulons in the developing human cortex atlas used for the common-variant analysis (**Methods**).

In ASD, scRBP identified 71 rare-variant-associated regulons involving 54 RBPs, with signals concentrated in neuronal and progenitor populations, particularly NbN, IPC and ExN (**Fig. 7a,b**; **Supplementary Table 41**). 14 of these RBPs had previously been implicated in ASD or other neurodevelopmental disorders through rare-variant studies (**Supplementary Table 42**). UTR-targeting regulons accounted for 66% of associations (3′UTR, 35%; 5′UTR, 31%), compared with 16% targeting CDS and 18% targeting introns (**Fig. 7b**), mirroring the common-variant results (**Fig. 6c**). Nearly half of ASD-associated regulons overlapped devRegulon set (45.1%), predominantly Trends 7-10, which were linked to neurogenesis, neural precursor proliferation and synapse organization (**Fig. 5c**; **Supplementary Fig. 30a**). Common- and rare-variant-based TRS profiles were modestly but significantly correlated across regulon-cell-type pairs (ρ = 0.19, P = 8×10^−68^; **Fig. 7c)**, indicating partial convergence between the two genetic architectures. Core rare-variant-associated regulators included *RBFOX1–3, CPEB4, MATR3, CELF2, CELF4, SRRM4* and *ELAVL3*, primarily in ExN and NbN, several of which were also supported by the common-variant analysis (**Fig. 7c**; **Supplementary Table 43**).

**Figure 7.**
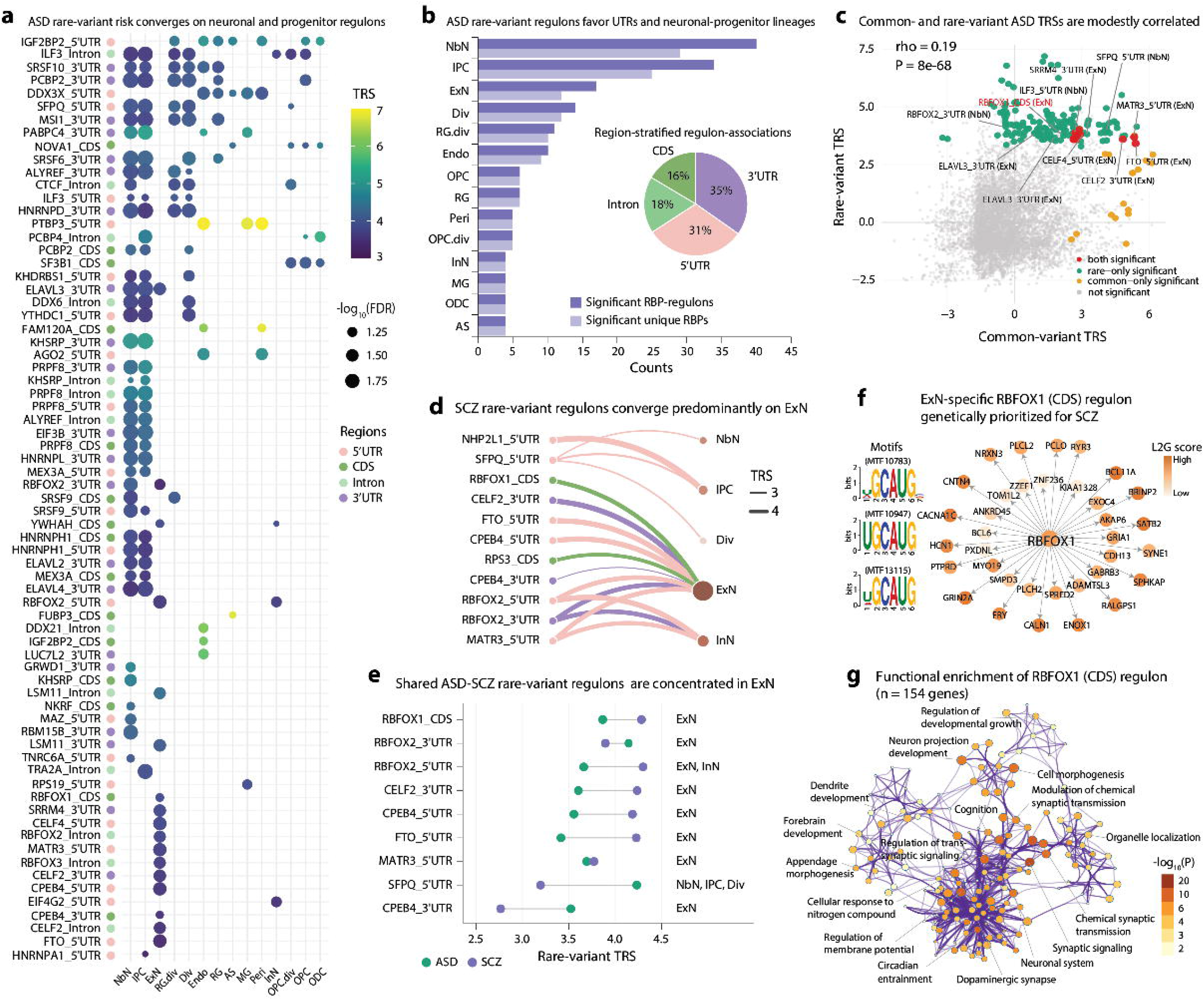
Rare-variant risk in ASD and SCZ converges on shared excitatory-neuron RBP regulons. (a) Cell type-resolved RBP regulons associated with ASD rare-variant burden. The 164 significant regulon–cell type associations comprised 71 regulons from 54 RBPs across 14 cell types. Rows denote regulons defined by an RBP and its targeted transcript region, and columns denote cell types. Dots indicate associations significant across all three scRBP components (FDR_RAS_, FDR_RGS_ and FDR_TRS_ < 0.1). Dot colour denotes the trait-relevance score (TRS), and dot size denotes −log_10_ of the maximum FDR across the three tests. Row annotations indicate the targeted transcript region. (b) Numbers of significant regulons and unique RBPs in each cell type, ordered by regulon count. Inset, distribution of the 164 significant associations across transcript regions: 3′UTR, 35%; 5′UTR, 31%; intron, 18%; and CDS, 16%. (c) Relationship between common- and rare-variant TRSs across all ASD regulon–cell type pairs tested (n=8,064). Points are classified as significant in both models, in the rare-variant model only, in the common-variant model only or in neither model. Rare-variant significance required FDR_RAS_, FDR_RGS_ and FDR_TRS_ < 0.1; common-variant significance was defined as in Fig. 6. Spearman’s correlation is shown (ρ=0.19, two-sided P=8×10^−68^). Selected regulons are labelled, and the RBFOX1 (CDS) regulon in ExN is highlighted in red. (d) Bipartite representation of the 16 regulon–cell type associations significant for SCZ rare-variant burden, comprising 11 regulons across 5 cell types. Arcs denote significant associations, with colour indicating transcript region and width proportional to TRS. Cell-type node size reflects the number of associated regulons. (e) Nine regulons that were significant in both ASD and SCZ in at least one shared cell type. Points show rare-variant TRSs in ASD and SCZ, and connecting lines indicate differences between disorders. Cell types in which each regulon was significant in both disorders are shown at right. Regulons are ordered by their mean TRS across ASD and SCZ. (f) ExN-specific RBFOX1 (CDS) regulon genetically prioritized for SCZ. The network shows RBFOX1 and its predicted targets, with node colour indicating the corresponding L2G score. Representative RBFOX1-binding motifs are shown at left. (g) Functional enrichment network for the RBFOX1 (CDS) regulon (n = 154 target genes). Term–term similarity network generated by using Metascape. Nodes denote enriched terms, edges denote term overlap and node color indicates −log_10_(P).

Extending the analysis to SCZ identified 16 cell-type-resolved regulon-trait associations involving nine RBPs, most of which localized to ExN (**Fig. 7d**; **Supplementary Fig. 30b**; **Supplementary Table 44**). TRSs derived from common- and rare-variant analyses were again modestly but highly significantly correlated across SCZ-associated regulon–cell-type contexts (ρ = 0.24, P = 2 × 10^−108^; **Supplementary Fig. 30c**). Regulons prioritized by both models included RBFOX1 (CDS), RBFOX2 (3’UTR and 5’UTR), CELF2 (3’UTR), CPEB4 (5’UTR), and MATR3 (5’UTR). Cross-disorder analysis identified nine rare-variant regulons that were significant in both ASD and SCZ in at least one matched cell type, seven of which were shared in ExN (**Fig. 7e**; **Supplementary Table 45**). By contrast, applying the same framework to rare-variant associations for the late-onset neurodegenerative disorders AD and PD, as well as the non-neuronal negative-control trait LDL cholesterol, identified no regulons meeting joint significance across RAS, RGS, and TRS at either FDR < 0.05 or FDR < 0.1 (**Supplementary Fig. 30d**), supporting the specificity of these associations to neurodevelopmental psychiatric disease contexts.

The ExN-specific RBFOX1 (CDS) regulon represented a prominent point of convergence between common- and rare-variant risk. *RBFOX1* is a highly conserved neuronal splicing factor that recognizes (U)GCAUG motifs and regulates programs implicated in ASD and SCZ^57,86,87^. scRBP reconstructed an ExN-specific RBFOX1 (CDS) regulon defined by (U)GCAUG motifs and associated with multiple psychiatric disorders (**Fig. 7f**; **Supplementary Fig. 31a**). Its target genes were enriched for synaptic signaling, dendrite development and forebrain development, and were linked to neurodevelopmental and cognitive phenotypes (**Fig. 7g**; **Supplementary Fig. 31b**,**c**). Moreover, 32.5% of targets had annotated interactions with FDA-approved drugs, including *SCN2A* with *lamotrigine* and *carbamazepine* and *CACNA1C* with *haloperidol decanoate* and *valproic acid* (**Supplementary Fig. 31d**; **Supplementary Table 46**). Together, these findings reveal that common- and rare-variant associations in ASD and SCZ partially converge on developmentally patterned excitatory-neuron RBP programs.

## DISCUSSION

Here we present scRBP, a framework that reconstructs cell-specific RBP regulons at gene and isoform resolution and links them to common- and rare-variant genetic associations. Across 6.2 million cells spanning 32 tissues and 50 complex traits, scRBP revealed that RBP-mediated regulation is strongly organized by cellular lineage and developmental contexts, while related traits frequently share post-transcriptional regulatory programs. In the developing human brain, common- and rare-variant associations further showed partial convergence on neuronal RBP regulons, highlighting post-transcriptional regulation as a cell-specific layer of complex-trait genetic architecture.

RBP regulons captured a regulatory layer that was complementary to TF-centred networks^21,22,32–34,88^. Although a subset of factors, particularly zinc-finger proteins, exhibited both RNA- and DNA-binding activities, their predicted targets and binding preferences were largely distinct. RBP regulons recovered cellular identity as effectively as TF regulons, whereas combining the two regulatory representations provided the strongest performance, indicating that transcriptional and post-transcriptional networks encode complementary features of cellular state. Regulon-level modelling may also improve robustness to the sparsity and technical variation inherent to single-cell transcriptomes by aggregating signals across coordinated target sets rather than relying on individual genes^21,41,89^. Importantly, targets inferred across hematopoietic lineages were enriched among genes responsive to perturbation of the cognate RBP in a genome-scale CRISPRi Perturb-seq screen in K562 cells. Although these experiments do not validate every cell-type-specific regulatory edge, they provide orthogonal functional support for the inferred RBP–target relationships.

Across diverse traits, scRBP recapitulated established cellular lineage relationships. Their regulon-trait associations were reproducible across independent datasets and frequently shared among related phenotypes while remaining strong cell-context specificity. These findings extend previous efforts^41,43,44,46^ to map genetic associations to tissues and cell types by resolving an RBP-centered regulatory layer. Rather than implicating individual genes in isolation, scRBP identifies coordinated post-transcriptional programs through which dispersed genetic signals may converge, consistent with network-based models of complex-trait architecture^35,41,43,47,90^. The observed lineage structure further suggests that related cell populations share components of RBP-mediated regulation, whereas these programs diverge across more distantly related lineages^22,91^.

The developing human cortex provided a prominent example of this organization. RBP regulons followed ten temporally ordered trajectories spanning progenitor proliferation, neurogenesis and neuronal maturation. Neuropsychiatric common-variant associations were concentrated in neuronal and progenitor populations and frequently involved developmentally dynamic regulons. Transient mid-gestational programs were linked to neurogenesis and neuronal migration, whereas later programs were associated with synaptic organization and circuit maturation, consistent with established developmental trajectories of human corticogenesis^76,81^. Members of the RBFOX, CELF and ELAVL families were recurrently prioritized, consistent with their established functions in neuronal RNA processing^3,6,7,57,58,78,92^. Importantly, the implicated regulons extended beyond splicing-related programs to multiple layers of post-transcriptional regulation, including RNA stability, localization and translational control, suggesting a broader contribution of RBP-mediated RNA regulation to neuropsychiatric disease risk. Rare-variant analyses in autism and schizophrenia independently implicated many of the same neuronal regulatory systems, with the strongest convergence in excitatory-neuron regulons. The modest correlations between common- and rare-variant-based TRS scores indicate partial rather than complete overlap, suggesting that the two genetic architectures perturb related post-transcriptional programs through partly distinct sets of genes. Together, these results identify RBP regulation as a convergent molecular layer connecting developmental cell states with both common and rare neuropsychiatric risk.

Several limitations should be considered. First, experimental maps of RBP–RNA interactions remain concentrated in a small number of cell lines and incompletely capture cell-type- and isoform-specific binding. The K562 CRISPRi Perturb-seq analysis provides orthogonal support for the inferred RBP–target relationships, but its single-cell-line context and potential indirect transcriptional responses preclude direct validation across all tissues and lineages examined here. Second, motif availability and quality vary substantially among RBPs, and the current collection is focused primarily on human data. Incorporating additional CLIP datasets and orthologous RBP motifs from model organisms should expand motif coverage and improve cross-species transferability^5,36^. Third, regulon–trait associations do not establish causality and may reflect direct regulatory relationships, indirect co-expression effects or shared cellular programs^41,43,44,93,94^. Fourth, the common-variant model relies primarily on proximity-based SNP-to-gene mapping and therefore incompletely captures distal regulatory interactions, whereas rare-variant analyses depend on the calibration and comparability of externally derived gene-level association statistics. Finally, empirical significance depends on the construction and depth of matched null regulons, although requiring concordant significance across RAS, RGS and TRS provides a conservative prioritization strategy. Integration of cell-context-specific CLIP data, isoform-resolved perturbational profiles and long-range variant-to-gene maps should further improve regulatory resolution and mechanistic interpretation.

In summary, our analyses reveal RBP-mediated post-transcriptional regulation as a cell-type- and lineage-specific layer of complex-trait genetic architecture. Across tissues and developmental states, genetically implicated RBP regulons were frequently shared among related traits yet remained strongly constrained by cellular context, with common- and rare-variant associations further converging on overlapping neuronal programs in the developing human brain. Together, these findings position post-transcriptional regulation as a cell-specific regulatory layer through which human genetic variation is organized across development and disease.

## METHODS

### scRBP methodology

In brief, scRBP is a unified and scalable computational framework for reconstructing cell type-specific RBP-gene and RBP-isoform regulons from short- and long-read single-cell transcriptomes, respectively, and for prioritizing regulons associated with complex traits and diseases. The framework integrates motif-guided network inference, single-cell regulon activity scoring and parallel common- and rare-variant association models to quantify the cellular and genetic relevance of each regulon. The overall workflow comprises six analytical modules, summarized in **Fig. 1**. Detailed methods are described in the following steps.

### Systematic collection and curation of RBP motifs

In the initial freeze (version 1, 10/10/2024) of scRBP, we systematically assembled a comprehensive catalogue of 616 RBPs and their associated motifs from 13 publicly available resources, including 12 curated databases and 26 CLIP-seq datasets deposited in GEO repository. The aggregation incorporated motifs from established databases (e.g., ATtRACT^5^ and CISBP-RNA^95^) as well as motifs newly inferred from CLIP-seq data (e.g., iCLIP, eCLIP, PAR-CLIP, and HITS-CLIP) using the HOMER pipeline^96^ (**Supplementary Note 1**). Sequence-specific binding preferences were encoded as position probability matrices (PPMs), which quantify the probability of each nucleotide (A, C, G, U) at each motif position. In total, this effort yielded 30,357 experimentally supported motifs corresponding to 616 RBPs.

To obtain a high-fidelity, non-redundant motif collection, we implemented a multi-step filtering and consolidation procedure. First, motifs with identical PPMs from different sources were merged, resulting in 29,180 unique motifs. Motif-to-motif similarity was then quantified using *Tomtom* (MEME Suite v5.5.7)^97^, and motifs of the same length with highly significant similarity (q-values < 1×10^−40^) were merged to reduce redundancy while minimizing inappropriate consolidation of weakly related motifs. Motifs with low information content (IC < 5) were removed to exclude low-complexity or poorly specified binding patterns^22,98^. To further refine motif boundaries, flanking positions with IC ≤ 0.1 were trimmed, and base probabilities ≤ 0.025 were rounded using the *universalmotif* R/Bioconductor package^99^, thereby reducing noise from poorly supported positions. Together, these filtering steps yielded 20,746 high-confidence motifs. As a complementary quality assessment, we additionally calculated HOMER-derived log-odds scores^96^, which quantify enrichment of observed versus expected nucleotide frequencies.

For motif clustering, we focused on the 8,041 motifs showing significant similarity to at least one other motif (*Tomtom*-based q-value < 1×10^−3^), while 12,345 motifs without significant matches were retained as singletons, consistent with previous studies^21,22^. *Tomtom*-based q-values were transformed into a motif similarity score (MSS) matrix using:

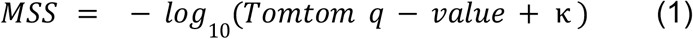

where κ is a small constant added to avoid undefined values when q-values approach zero. The *MSS* matrix was normalized, scaled, and subjected to principal component analysis (PCA) using Seurat (v5.4.0)^100^. The top 100 principal components (PCs) were used to construct a shared nearest-neighbor graph via *Seurat::FindNeighbors()* function, followed by Leiden clustering. Based on empirical testing of clustering resolutions, a resolution of 15 was selected to achieve an appropriate level of clustering granularity, yielding 164 initial motif clusters. For visualization, dimensionality reduction was performed using t-distributed Stochastic Neighbor Embedding (t-SNE) on the same 100 PCs.

To derive high-resolution and non-redundant motif groups, we further refined these clusters using RSAT matrix-clustering^101^, which builds a hierarchical motif tree based on pairwise PPM similarity and partitions motifs into subclusters. This procedure produced 1,575 motif subclusters (mean = 5.45 motifs per subcluster), each represented by a consensus PPM and motif logo. RBP annotations for each subcluster were aggregated from all contributing motifs, acknowledging that a single motif may map to multiple RBPs and that individual RBPs may have multiple distinct motifs.

### Transcriptome-wide ranking of motif-target genes

RBPs interact with *cis*-acting regulatory elements distributed across entire transcripts—including the 5’UTR, CDS, intronic, and 3’UTR—to mediate diverse co-/post-transcriptional regulatory processes^8,102^. To quantify RBP binding potential in a domain-resolved manner, we generated eight distinct transcriptome-wide motif-target ranking databases, corresponding to motif occurrences within each of the four transcript domains at both the gene and isoform resolution.

Domain-specific motif scanning was performed using Cluster-buster (v1.0)^50^, a Hidden Markov model-based algorithm that detects clusters of motif occurrences rather than isolated matches. Cluster-buster evaluates the enrichment of motif clusters using a log-likelihood ratio score as previously applied in regulatory network analyses^21,22^. In scRBP, motifs belonging to the same motif cluster were scanned jointly to identify composite binding regions enriched for any motif within the cluster. Transcript sequences were extracted using GENCODE v44 annotations^103^, incorporating all annotated isoforms for isoform-level analyses. For gene-level analyses, we preserved transcript-region specificity by merging isoform-derived regions within each transcript domain, resulting in four gene-level scanning regions per gene.

To provide orthogonal evidence and strengthen motif-target predictions, we implemented four complementary motif-scanning strategies: (i) Individual motif (uncluster) scanning, where each motif was scanned independently using Cluster-Buster^50^. (ii) Consensus motif (archetype) scanning, using one representative motif from each of the 1,575 subclusters derived from RSAT *matrix-clustering*^*101*^. (iii) FIMO-based motif scanning (MEME Suite v5.5.7)^104^, which scores motif matches using PPM-derived log-likelihood statistics (threshold: P < 10^−3^). (iv) HOMER2-based motif scanning^96^, which quantifies motif occurrence and positional enrichment relative to background expectations (**Supplementary Note 2**).

Each strategy generated a quantitative motif-binding score (*BS*) for each motif-region pair, reflecting predicted binding strength and regulatory potential. Binding regions were mapped to genes or isoforms using BEDTools^105^. For each motif, genes or isoforms were ranked in descending order of *BS* to generate transcript region-resolved motif-target rankings. Targets without detectable motif overlap (e.g., with zero or missing *BS*) were assigned random ranks at the bottom of the ranking list. To benchmark the specificity and accuracy of the motif-target rankings, we implemented an AUC-based recovery framework using 223 ENCODE eCLIP-seq datasets^8^ covering 103 RBPs in HepG2 and 120 RBPs in K562. Performance was assessed using RBP-centric recovery, target-centric recovery, and cross-cell-line reproducibility. Detailed benchmarking procedures are provided in the **Supplementary Note 3**.

### Constructing RBP-target regulatory networks from single-cell data

To infer preliminary RBP-target regulatory networks, scRBP implements a gradient boosting regression algorithm (GRNBoost2)^51^. Given a predefined set of RBPs (n = 616) and a single-cell gene-by-cell or isoform-by-cell expression matrix, the *getModule()* function in scRBP fits an individual gradient-boosted regression model for each expressed feature (gene or isoform), predicting its expression across cells from the expression levels of all candidate RBPs. The resulting importance measures (IMs) are interpreted as directed edge weights, yielding a ranked adjacency matrix of putative RBP-target interactions. To enhance robustness in sparse single-cell data and reduce overfitting^106,107^, scRBP employs stochastic gradient boosting with out-of-bag early-stopping regularization, and repeats model training across 30 random initializations by default (extended to 50-100 iterations for selected case studies). Final edge weights are obtained by averaging IMs across runs. Within this framework, higher IMs represent stronger putative regulatory influence, enabling the prioritization of high-confidence RBP-gene or RBP-isoform interactions, consistent with established network-inference strategies^21,22^.

To benchmark the accuracy of RBP-target interactions inferred from single-cell data, we additionally evaluated four well-established GRN inference methods—DeepSEM (GPU-based)^106^, PIDC^108^, GENIE3^109^, and ARACNe-AP^107^ (**Supplementary Note 4**). Following an earlier study^4^, ENCODE eCLIP datasets from two cell lines (i.e., HepG2 and K562) were used as experimentally validated ground truth for RBP-RNA interactions. For HepG2, we analyzed two scRNA-seq datasets obtained through Smart-seq2 (n = 68 cells)^110^ and DNBelab data (n = 1,628 cells)^111^; for K562, we evaluated CEL-seq (n = 239 cells)^112^ and STORM-seq (n = 70 cells)^113^ datasets. Consistent with our benchmarking results and previous studies^4,20^, the gradient boosting machine method GRNBoost2^51^ showed superior performance and was therefore selected as the default inference engine in scRBP. To enhance computational scalability, scRBP incorporates automated partitioning of large expression matrices together with parallelized execution. As an alternative option, scRBP also provides support for GENIE3^109^, a random forest-based GRN inference approach.

Moreover, we systematically evaluated 15 module-construction strategies for defining RBP regulatory modules. These included three classes of selection rules: top *n* targets per RBP (*n* = 50, 100, 200, 300, 500 or 1,000), top *n* RBPs per target (*n* = 3, 5, 10, 25 or 50), and percentile-based IM thresholds per RBP (50th, 75th, 90th or 95th percentile). Interactions with IM < 0.005 were excluded in all cases. To compare strategies, we calculated pairwise Jaccard similarity indices (JSI) based on overlap between inferred target sets for each RBP, and averaged these across RBPs to obtain a global strategy-to-strategy similarity matrix. Hierarchical clustering of the corresponding JSI distance matrix identified five coherent groups representing distinct module-construction principles. For visualization, we performed PCA on a strategy-by-feature matrix summarizing module size and overlap structure. We further assessed each strategy by bootstrap-based enrichment analysis, testing whether recovered modules were enriched for significant RBP modules from the full dataset, defined as modules with greater connectivity than those obtained from label-shuffled null distributions (100 permutations). Enrichment *Z*-scores were calculated relative to overlaps from 100 randomly generated control strategies. Based on these analyses and prior studies^21,22^, scRBP adopts six representative default strategies: top 50 targets per RBP, top 5, 10 or 50 RBPs per target, and the 75th and 90th IM percentile thresholds.

To classify the resulting RBP-regulatory modules into activating or repressing categories, scRBP calculates Spearman correlation coefficients between each RBP and its putative targets, leveraging the full expression matrix to alleviate sparsity-related bias. Following previous approaches^21,22^, RBP-target pairs with positive correlation coefficients ρ ≥ 0.03 are classed as activating links, while those with negative correlation coefficients ρ ≤ −0.03 are classed as repressing links. Using a permissive correlation threshold allows for the inclusion of both linear and nonlinear regulatory effects, reflecting the complex and context-dependent nature of co-/post-transcriptional regulation mediated by RBPs^2^. Because very small modules are unstable and sensitive to dropout^20,41,89^, scRBP by default retains only modules containing more than 20 genes (or isoforms), unless specified otherwise. The final output of the *getModule()* function in scRBP package is a ranked list of RBP-gene or RBP-isoform interactions, each annotated with an IM score and a correlation coefficient, representing inferred regulatory strength and activation mode, respectively. In order to support scalability in large-scale single-cell datasets, scRBP provides a built-in function *getSketch()* that applies geometric sketching algorithm^53^ to select a transcriptomically representative subset of cells.

### Identification of direct RBP-target interactions

To delineate direct RBP-target interactions, scRBP applies a ranking-based motif enrichment framework that maps each co-expression-derived regulatory module to high-fidelity target genes or isoforms. Preliminary RBP-target relationships inferred from single-cell expression data may include both direct and indirect effects, as they arise solely from co-expression patterns without explicitly incorporating physical RBP-RNA binding information. To refine these networks, each predicted RBP-regulatory module is treated as a foreground gene or (isoform) set, and enrichment is evaluated using precomputed transcriptome-wide motif-target rankings (“20K motif collection”) derived from 616 RBPs across four transcript domains (5’UTR, CDS, intronic, and 3’UTR) at both gene and isoform levels.

Enrichment is quantified using the area under the cumulative recovery curve (AUC), which measures the extent to which foreground genes or isoforms are preferentially ranked toward the top of each motif’s background ranking. To focus on high-confidence signals, AUC is computed only over the upper portion of each background ranking—by default, the top 5% for gene-level rankings and the top 1% for isoform-level rankings. The distribution of AUC values across all motifs forms an empirical reference. For each motif *b* (*b ∈* {1, 2, …, *B*}), the normalized enrichment score (*NES*) is defined as:

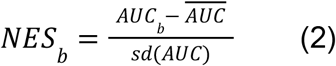

where 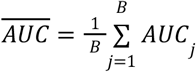 denotes the mean AUC across all motifs (with *j* indexing motifs), 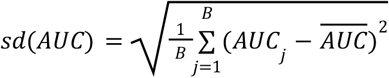 is the corresponding standard deviation, and *B* is the total number of motifs. NES thus captures the standardized deviation of each motif’s AUC from the empirical background distribution, providing a quantitative measure of motif enrichment. By default, motifs with NES ≥ 3.0 are considered notably enriched; greater specificity can be achieved by increasing the threshold to 4.0, whereas a threshold of 2.5 increases sensitivity for detecting additional enriched motifs.

For each enriched motif, scRBP further defines a leading-edge index on the recovery curve to identify a subset of high-confidence direct targets, consistent with prior approaches^21,114–116^. The leading edge (*LE*) is defined as the rank position at which the deviation between the motif-specific recovery curve and the background reference curve—defined as the pointwise mean recovery plus two standard deviations—is maximized. Formally, for motif *b*, the *LE* is calculated as:

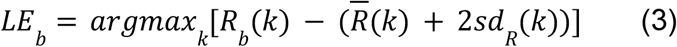

Where 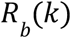 is the cumulative recovery curve for motif *b* at rank position *k*, 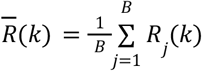 denotes the pointwise mean recovery curve across all motifs, and 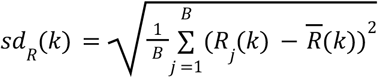 indicates the pointwise standard deviation. This automated thresholding procedure identifies the most discriminative point at which the recovery curve of an enriched motif exceeds the background expectation by more than two standard deviations. Only genes or isoforms within the foreground module that are ranked above *LE*_*b*_ are retained as direct targets in the resulting regulon. If no enriched motif can be assigned to an RBP, the corresponding module is discarded. The sets of predicted direct target genes (or isoforms) from all remaining modules related to the same RBP are then consolidated into a single unified regulon. The *getRegulon()* function in scRBP returns these high-fidelity regulons across all motifs in the “20K motif collection”.

### Common-variant SNP-to-gene mapping and gene-level association scoring

To link common-variant associations to RBP regulons, we systematically benchmarked six SNP-to-gene mapping windows (0, 10, 20, 50, 100 and 200 kb) using two independent single-cell datasets from PBMC and midbrain, together with GWAS summary statistics for ten blood-cell traits and nine aging-related traits. On the basis of these analyses, we selected a 50-kb window centered on the transcription start site (TSS) as the default mapping setting in scRBP. This choice is consistent with previous observations that a substantial fraction of expression quantitative trait loci reside near the TSS across diverse cell types and tissues^93,117^.

For each SNP *u ∈ U*, MAGMA^118^ converted the marginal GWAS P value into an equivalent two-sided χ^2^ statistics. SNP-level association statistics assigned to each gene are then aggregated into a gene-level test statistic while accounting for local linkage disequilibrium (LD) using ancestry-matched reference genotypes. Unless otherwise specified, scRBP uses the MAGMA SNP-wise mean χ^2^ model for the set of SNPs *U*_*g*_ = {*u*_1,*g*_, *u*_2,*g*_, …, *u*_|*U*|,*g*_} assigned to gene *g*. The resulting gene-level P values are transformed into standard normal scores and used as input for downstream regulon-level regression.

### Computing common-variant regulon-level genetic association score

After SNP-level associations were aggregated into gene-level statistics, scRBP evaluated the common-variant association of each RBP regulon using a competitive gene-set regression framework^118^. Competitive tests assess whether genes within a focal regulon show stronger association with the trait than genes outside that regulon, while controlling for gene-level genomic properties^119^.

Let *S* denote the set of regulons, and let 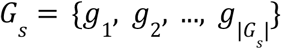 denote the genes assigned to regulon *s ∈ S*. For each regulon, the gene-level association score is modelled as:

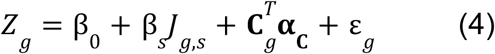

where *Z*_*g*_ is the LD-adjusted gene-level association score for gene *g, J*_*g,s*_ = 1 if gene *g ∈ G*_*s*_ and *J*_*g,s*_ = 0 otherwise, and β_*s*_ quantifies the excess common-variant association attributable to membership in regulon *s*. The vector **C**_*g*_ contains gene-level covariates, including gene size, SNP density, and effective gene size, together with their log-transformed terms. Effective gene size represents the effective number of SNP parameters after accounting for local LD. The vector **α**_**C**_ contains the corresponding regression coefficients, and ε_*g*_ denotes the residual error.

For each regulon *s*, the common-variant regulon-level genetic association score (RGS) is defined by default as the standardized regulon-membership effect: 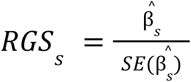 Under this formulation, a larger *RGS* indicates stronger common-variant association among genes within the focal regulon than among other genes in the analysed gene universe, after adjustment for genomic covariates. Alternatively, scRBP supports a non-directional RGS formulation, *RGS*_*s*_ = − *log*_10_(*P*_*s*_), where *P*_*s*_ is the regulon-level association P value.

### Computing rare-variant regulon-level genetic association scores

For the rare-variant model, scRBP uses precomputed gene-level association statistics from rare-variant analyses of whole-genome or whole-exome-sequencing studies. Depending on the statistical framework of the source study, gene-level evidence is represented by association P values, log Bayes factors or directly supplied quantitative scores. These statistics are harmonised to the genes represented in the regulon universe and converted into standardised gene-level rare-variant association scores. Because rare-variant evidence is already aggregated at the gene level, no SNP-to-gene mapping or LD reference panel is required.

For each regulon *s*, scRBP evaluates rare-variant association using a competitive gene-level regression model:

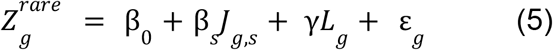

Where 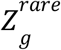 is the standardised gene-level rare-variant association score for gene *g, J*_*g,s*_ indicates whether gene *g* belongs to regulon *s*, and *L*_*g*_ denotes standardized log-transformed union coding-sequence (CDS) length. CDS length accounts for differences in coding mutational target size among genes. The rare-variant RGS was defined analogously to the common-variant RGS, using either the standardized regulon-membership effect from the CDS-length-adjusted regression or the corresponding non-directional − *log*_10_ (*P*_*s*_) score, as detailed in the **Supplementary Note 5**.

### Quantifying regulon activity score at single-cell level

To quantify the regulon activity score (RAS) of each RBP-regulon at single-cell resolution, scRBP adopts the widely used AUCell algorithm^21^, which evaluates whether the genes within a regulon are preferentially enriched among the highest-expressed genes in a given cell. In this context, the RAS reflects both the proportion of regulon genes that are expressed and their relative expression ranking compared with all other genes in the cell, rendering the method independent of expression units and normalization procedures.

For each cell *c*, all detected genes 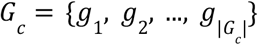 are ranked in descending expression, generating a cell-specific ranked expression profile (ties such as zero values are randomly ordered). Let *h*(*i*) = 1(*g*(*i*) *∈ G*_*s*_) denote the indicator function specifying whether the gene at rank *i* belongs to regulon *G*_*s*_. The cumulative recovery curve was calculated using: 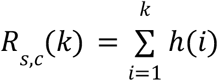, repres enting the number of regulon genes recovered within the top *k* ranked positions. scRBP computes the raw activity score of regulon *s* in cell *c* as the area under this cumulative recovery curve within a predefined top-rank window *K* (by default, the top 5% of expressed genes): 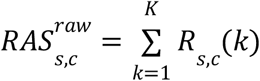. To ensure comparability across regulons of different sizes, the raw activity score is normalized by the theoretical maximum area when all regulon genes appear within the top *K* rank positions. The final normalized RAS for regulon *s* in cell *c* is defined as:

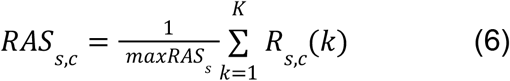

where *R*_*s,c*_ (*k*) denotes the cumulative recovery of regulon genes in cell *c*, and *maxRAS*_*s*_ represents the maximum possible area under the recovery curve given |*G*_*s*_| and *K*. Thus, *RAS*_*s,c*_ *∈* [0, 1] quantifies the enrichment of regulon genes among the highest-ranked genes in each cell, with higher values indicating greater post-transcriptional activity of the corresponding RBP.

### Inference of cell-type-resolved regulon activity

To obtain robust cell type-level estimates of regulon activity, scRBP aggregates single-cell RAS profiles using an entropy-based Jensen-Shannon divergence (JSD) algorithm^54,120^. For regulon *s*, let **r**_*s*_ = (*r*_*s*,1_, *r*_*s*,2_, …, *r*_*s,N*_) denote its *RAS* values across *N* cells. These values are normalized to define a probability distribution: 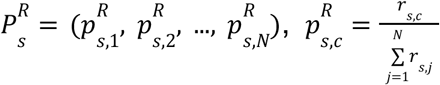. Let *M* denote the number of predefined cell types, and let *C*_*m*_ (*m ∈* {1, 2,…, *M*}) denote the *m*-th cell type. For each cell type *C*_*m*_, scRBP defines a corresponding cell-type membership distribution 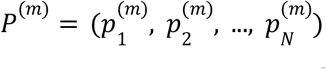 where 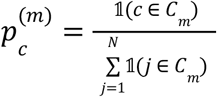. The divergence between the regulon-derived distribution 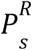 and the cell-type membership distribution *P*^(*m*)^ is quantified using the *JSD*:

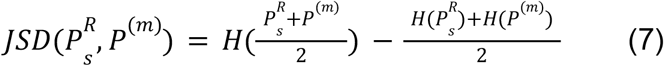

where 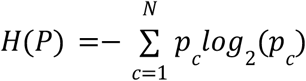 denotes the Shannon entropy. Under this definition, *JSD* values range from 0 for identical distributions to 1 for maximally distinct distributions.

To convert this divergence into an interpretable measure of cell-type-specific regulon activity, scRBP defines the cell-type-level regulon activity score as:

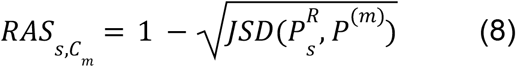

Higher values indicate that regulon *s* is more strongly and specifically active within cell type *C*_*m*_. For clarity, we describe the gene-level formulation here; the isoform-resolved regulon activity score is defined analogously using isoform-ranked expression profiles from single-cell long-read sequencing. This entropy-based aggregation framework mitigates sparsity-related noise in single-cell data while preserving biologically meaningful, cell-type-specific regulatory signals^21,22,54^.

### Calculation of trait-relevance scores

To prioritize RBP-regulons that are both active in a given cellular context and genetically associated with a trait, scRBP combines the RAS with the corresponding RGS to calculate a trait-relevance score (TRS). Common- and rare-variant models are applied independently, such that the common-variant or rare-variant RGS is combined separately with RAS to generate the corresponding TRS. TRSs can be calculated at both single-cell and cell-type resolution.

To place RAS and RGS on comparable scales while limiting the influence of extreme outliers, each score distribution was capped at its 99th percentile and subsequently transformed by min-max scaling to the interval [0,1]. For the common- and rare-variant models, RGS normalization was performed separately within each analysis. For a given RBP regulon *s ∈* {1, 2, …, |*S*|} in a cell *c ∈* {1, 2, …, *N*}, let 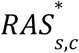 denote the scaled regulon activity of *s* in cell *c* and let 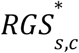 denote the corresponding scaled common- or rare-variant genetic association score. Following previous work^35^, TRS for regulon *s* in cell *c* is defined as:

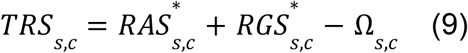

where 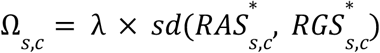 is a regularization term that penalizes discordance between regulon activity and genetic association, and λ is a tunable parameter (λ = 1 by default). Treating 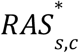 and 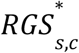 as two observations, their sample standard deviation is 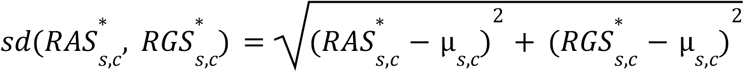, where 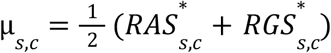 is their mean, and then the standard deviation is calculated as:

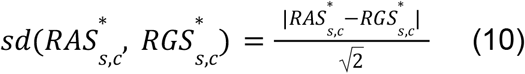

The penalty therefore increases monotonically with the discrepancy between cellular activity and genetic association, and equals zero when the two scores are identical. Under this formulation, regulons that are concordantly high RAS and RGS consequently receive high TRS values, whereas regulons supported predominantly by only one modality are downweighted (**Supplementary Note 6**). All scaling, TRS calculation and empirical significance testing were performed independently for the common- and rare-variant models.

scRBP supports two complementary levels of cellular resolution. In single-cell mode, TRS is calculated for each regulon in each cell, enabling fine-scale characterization of heterogeneity in trait-associated regulon activity. In cell-type mode, the same formulation is applied to cell-type-level RAS and the corresponding RGS to identify regulatory programs preferentially associated with a trait in specific cellular populations.

### Empirical significance testing of RBP regulons

To assess whether the observed activity, genetic association and trait relevance of an RBP regulon exceeded those expected for regulons with comparable properties, scRBP implements a Monte Carlo matched-null sampling framework. Common- and rare-variant models are evaluated independently, using model-specific covariates to construct empirical null distributions. For each observed regulon *s* comprising gene set 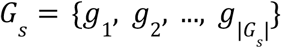, *D* matched null regulons 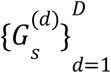 were generated while preserving regulon size and the distribution of relevant gene-level properties (*D* = 1,000 by default).

For the common-variant model, background genes are stratified according to four covariates: the number of SNPs assigned to each gene (*N*_*SNP*_), the effective number of independent SNP predictors after LD adjustment (*N*_*PARAM*_), mean expression across cells or cell types (*mean_expr*) and the fraction of cells with detectable expression (*pct*_*detected*_). Each covariate was discretized into quantile bins, and null regulons were sampled to reproduce the joint covariate-bin composition of the observed regulon. This procedure generated null sets matched for regulon size, SNP burden, LD-adjusted model complexity and expression properties. For the rare-variant model, matched null regulons were constructed using coding-sequence opportunity, represented by the standardized log-transformed union CDS length, together with mean expression and the fraction of cells with detectable expression. CDS length was included as a covariate in the rare-variant regression, whereas the expression-derived variables were used only for matched-null construction. The same competitive rare-variant regression was applied to the observed and null regulons. When the full three-dimensional matching stratum contained insufficient genes, sampling was progressively relaxed to strata defined by CDS length and mean expression, CDS length alone and, finally, the full matched gene universe, while preserving regulon size wherever possible.

For each observed regulon and cellular context, RAS, RGS and TRS were precomputed for all matched null regulons using the same analytical procedure as for the observed regulon. For the three scores, the one-side empirical P values are calculated as:

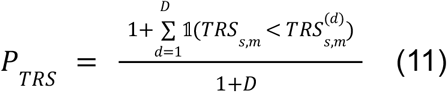

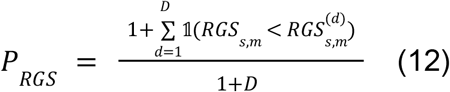

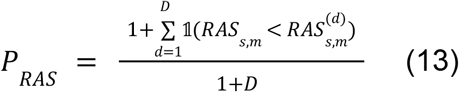

Where *TRS*_*s,m*_, *RAS*_*s,m*_ and *RGS*_*s,m*_ denote the observed scores for regulon *s* in a given cell type *C*_*m*_ (*m ∈* {1, 2, …, *M*}), and 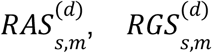 and 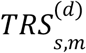 denote the corresponding score for matched null regulon *d*. The addition of one to the numerator and denominator prevents zero empirical P values. Null distributions and empirical P values were generated separately for the common- and rare-variant models.

Empirical P values for RAS, RGS and TRS were adjusted separately for multiple testing using the Benjamini–Hochberg procedure. Unless otherwise specified, a regulon was considered significant only when all three scores satisfied their respective false-discovery rate thresholds: *FDR*_*TRS*_ < 0.05, *FDR*_*RAS*_ < 0.05 and *FDR*_*RGS*_ < 0.05. This three-score criterion provides a conservative strategy for prioritizing regulons with both genetic support and cellular specificity. Analysis-specific thresholds could be applied where indicated.

To facilitate comparison across traits, cellular contexts and genetic models, observed scores were additionally standardized relative to their matched null distributions. Specifically, 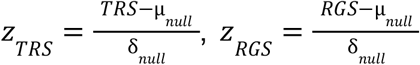, and 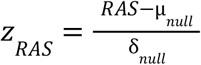, where µ_*null*_ and δ_*null*_ denote the mean and standard deviation of the corresponding matched-null score distribution, respectively. Regulons were subsequently classified into four categories according to trait relevance and cellular specificity: trait relevant and specific (Q1), trait relevant but nonspecific (Q2), specific but not trait relevant (Q3), and neither trait relevant nor specific (Q4).

### Systematic assessment of scRBP performance

To quantify the cell-type specificity of scRBP-inferred TRS profiles, we used the energy distance statistic^35,121^, a multivariate measure rooted in the physical analogy of gravitational energy. The *E-statistic* quantifies the degree of separation between two high-dimensional distributions by comparing their inter-group divergence to their respective intra-group variability. Intuitively, two distributions are considered distinguishable when the dissimilarity between them substantially exceeds the internal spread within each group.

Let *t*_1_, *t*_2_, …, *t*_|*S*|_ *∈ R*^|*S*|^ denote the TRS scores of |*S*| regulons in a target cell type, and 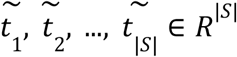 is the corresponding TRS values sampled from all non-target cell types. We compute the mean pairwise distance between the groups:

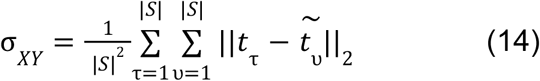

and the within-group dispersions for the target and non-target cells,

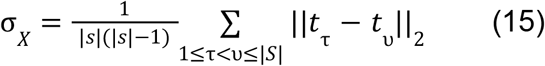

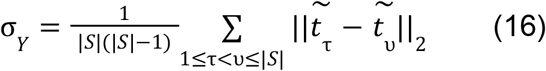

The E-statistic is then defined as:

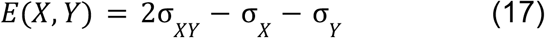

where σ_*X*_ and σ_*Y*_ characterize the internal compactness of each distribution, and σ_*XY*_ measures their mutual separation. *E*(*X, Y*) is positive when the two distributions are more separated from each other than from themselves, and increases monotonically with the degree of distributional divergence. Accordingly, a higher E-statistic indicates that the inferred regulons confer strong and cell-type-restricted TRS profiles, whereas a lower value suggests that regulatory activity is broadly shared across cell types with weak specificity.

### Motif targets overlap between dual-binding proteins

To systematically evaluate the complementarity between post-transcriptional and transcriptional gene regulatory networks, we compared the scRBP regulon framework with pySCENIC (v0.12.1)^122^, a widely used TF-centric regulon inference tool. The pySCENIC TF reference resource^122^ (hg38 genome assembly, motif collection v9) was intersected with the curated human TF catalogue reported by Lambert et al.^123^ to define an analysis-ready set of 1,390 human TFs with available cisTarget motif annotations^21^. The overlap between scRBP RBPs and pySCENIC TFs was quantified using a two-sided Fisher’s exact test on a 2 × 2 contingency table, with all human protein-coding genes as the reference background.

For the 66 proteins shared between the two collections, we next quantified concordance between TF-associated and RBP-associated target programs using Jaccard similarity. TF motif target rankings were retrieved from the pySCENIC cisTarget database, and RBP motif target rankings were obtained from the scRBP motif ranking databases across four transcript regions (5’UTR, CDS, intronic, and 3’UTR). For proteins associated with multiple motifs, motif-specific rankings were merged by minimum-rank aggregation. For each protein, the top 1,500 ranked targets were retained, and pairwise Jaccard similarity was computed between the TF target set and each region-specific RBP target set. Statistical significance of the observed target overlap was assessed using a permutation-based null distribution generated from 1,000 permutations in which protein labels were randomized while preserving the number of targets per protein. Differences in Jaccard similarity between TF motif targets and region-specific RBP motif targets were evaluated using a paired Wilcoxon signed-rank test.

### Cell-identity benchmarking using ENCODE cell lines

To benchmark the capacity of RBP regulons, TF regulons, and conventional highly variable gene (HVG) features to recover cellular identity, we performed 10 independent simulation replicates using bulk RNA-seq profiles of eight deeply profiled ENCODE cell lines (GM12878, K562, HCT116, MCF-7, HepG2, IMR-90, PC-3, and Panc1). In each replicate, TPM-normalized expression profiles derived from RSEM gene quantification were used to generate 200 pseudo-cells per cell line by negative binomial sampling (target library size, 5,000 counts per cell; dispersion parameter, 0.3), yielding 1,600 pseudo-cells per replicate.

Simulated count matrices were processed using Scanpy (v1.9)^124^. Genes detected in fewer than 10 cells and cells expressing fewer than 200 genes were excluded. Library sizes were normalized to 10,000 counts per cell and log-transformed using log1p. Four feature representations were evaluated: (i) the top 2,000 HVGs selected using the Seurat (v5.4.0)^100^, followed by scaling and PCA (50 components); (ii) RBP regulon activity scores inferred by scRBP; (iii) TF regulon activity scores inferred using the standard pySCENIC workflow; and (iv) a combined RBP + TF regulon representation. For regulon-based representations, AUCell-derived activity scores were used as the cell-by-feature matrix. For each feature representation, Leiden clustering was performed independently across 10 replicate simulations generated with distinct random seeds. Clustering performance was quantified by comparing inferred cluster assignments against ground-truth cell-line labels using the adjusted Rand index (ARI) and normalized mutual information (NMI). Differences in clustering performance across feature representations were evaluated using a paired Wilcoxon signed-rank test.

### Transcript-region binding landscape of RBPs across 30 human tissues

To characterize the transcript-region binding preferences of RBPs, scRBP was applied to single-cell transcriptomic datasets from 30 human tissues, encompassing fetal brain^60^, adult brain^61^, and 28 somatic tissues from the *Tabula Sapiens* atlas^125^, each processed through the full scRBP inference and regulatory network pruning pipeline. For each tissue, region-specific regulon gene sets were defined across four transcript regions: 5’UTR, CDS, intronic, and 3’UTR. A regulon was retained for downstream analyses if it satisfied all of the following criteria: normalized enrichment score (NES) ≥ 3.0, minimum regulon size ≥ 10 genes, and motif rank threshold ≤ 1,500. Enrichment events passing these thresholds were aggregated across all 30 tissues, and for each RBP, the proportion of enrichment occurrences attributable to each transcript region was computed and normalized to sum to unity, yielding a tissue-aggregated regional binding profile per RBP. RBPs with enrichment detected in at least five tissues were retained for comparative analysis and grouped by their dominant transcript region. Hierarchical clustering was applied within each regional group to reveal shared binding landscape patterns among RBPs with similar regional preferences. Disease relevance of region-dominant RBP classes was assessed by cross-referencing identified RBPs with Mendelian disease gene annotations from the Online Mendelian Inheritance in Man (OMIM) database^126^, with further stratification by curated gene lists for neurodevelopmental and psychiatric disorders.

### Perturb-seq evaluation of inferred RBP-target relationships

Five hematopoietic lineages from an integrated Human Cell Atlas reference were analysed separately with scRBP to construct lineage-specific RBP regulons. Within each lineage, regulon targets were further intersected with cell type-specific differentially expressed genes to generate cell type-resolved regulon instances. These regulons were evaluated against an independent genome-scale CRISPRi Perturb-seq screen in K562 cells^67^ targeting 9,866 genes. Gene-level differential-expression log-fold changes and P values for all measured response genes from each perturbation were obtained from the reanalysis by Ota et al.^47^. The analysis was restricted to 8,071 expressed non-ribosomal genes. After excluding the perturbed RBP itself, perturbation-responsive genes were defined either as the top 100, 200 or 500 genes ranked by ascending P value, or as all genes with Benjamini–Hochberg FDR < 0.05.

After harmonization of RBP identities and target genes between the scRBP and Perturb-seq datasets, 1,702 lineage- and cell type-resolved regulon instances were retained for evaluation. For each definition of perturbation-responsive genes, enrichment of predicted regulon targets among responsive genes was quantified relative to random expectation and summarized across each hematopoietic lineage. To account for differences in baseline expression and nonspecific perturbation responsiveness, we additionally calculated a cognate-to-noncognate specificity ratio, defined as the enrichment observed following perturbation of the cognate RBP divided by the mean enrichment observed across non-cognate RBP perturbations.

At single-regulon resolution, each regulon was evaluated across 496 RBP perturbations by ranking perturbations according to enrichment of its predicted targets among the top 1,000 perturbation-responsive genes. Performance was summarized using a truncated AUROC, and the percentile rank of the cognate RBP perturbation among all RBP perturbations was defined as the self-rank. Statistical significance was assessed using RBP-identity permutation tests, one-sided Wilcoxon signed-rank tests and one-sided binomial tests, with Benjamini–Hochberg correction for multiple testing. 95% confidence intervals were estimated by bootstrap resampling of regulons.

### Differential analysis of regulon activity across developmental stages

Differential regulon activity across developmental stages was assessed using the limma framework with empirical Bayes moderation^127^, following established pseudo-bulk analysis workflows^78,128^. Single-cell regulon activity scores were aggregated by donor and developmental stage to generate one pseudo-bulk profile for each donor-stage combination. An intercept-free design matrix was constructed with developmental stage as the primary variable of interest, and dataset was included as a covariate when multiple datasets were analysed jointly. Linear models were fitted to the pseudo-bulk regulon activity matrix using *limma::lmFit*. Pairwise contrasts were specified to compare each developmental stage with the earliest developmental stage as the reference and were fitted using *limma::contrasts*.*fit*. Empirical Bayes moderation was subsequently applied using *limma::eBayes* to shrink regulon-specific residual variances toward a common prior estimate, thereby stabilizing standard errors and improving statistical power.

For each contrast, effect estimates, moderated test statistics and *P* values were extracted. To summarize evidence for developmental regulation across stage contrasts, contrast-specific *P* values were combined for each regulon using Fisher’s method, yielding one omnibus *P* value per regulon. Omnibus *P* values were adjusted for multiple testing using the Benjamini–Hochberg procedure, and development-associated regulons (“devRegulons”) were defined at FDR < 0.05. Regulon-level developmental effects were summarized using the mean signed difference and mean absolute difference in activity across contrasts. Potential confounding variables were evaluated at the pseudo-bulk level, and covariates showing systematic associations with regulon activity were retained in the final model.

### Identification of developmental regulon trend clusters

Stage-specific mean regulon activity was calculated for each regulon by averaging pseudo-bulk activity across donors at each developmental stage. Developmental trajectories were then modelled as a function of developmental stage using generalized additive models (GAMs) with thin-plate regression splines, as implemented in the *mgcv* R package. Fitted response curves were predicted at 80 evenly spaced positions along the developmental axis to enable comparison across regulons. Fitting smooth curves to the pseudo-bulk trajectories, rather than clustering the observed activity values directly, facilitated the identification of coherent temporal patterns and distinct regulatory dynamics across development. Each fitted trajectory was subsequently rescaled to the [0,1] range to emphasize relative temporal dynamics independently of absolute activity magnitude.

The normalized trajectory profiles were clustered by hierarchical clustering using Euclidean distance and Ward’s minimum variance linkage, as implemented in the *SciPy* Python package (v1.7.1)^129^. The number of clusters was selected empirically to balance temporal distinctiveness and redundancy among trajectory patterns, yielding ten regulon trend clusters representing distinct developmental activation programs, including early, late, transient, and sustained patterns. For visualization, the 80-point interpolation grid was partitioned into eight bins corresponding to the predefined developmental-stage intervals, with stage boundaries indicated by vertical lines. Within each cluster, individual regulon trajectories were overlaid to illustrate intra-cluster heterogeneity. A cluster-level mean trajectory was calculated by averaging the normalized activity of member regulons at each position along the interpolation grid, with the surrounding shaded band representing variability across regulons. Cluster-level trajectories were displayed in a vertically faceted layout to facilitate comparison of temporal regulatory programs across development.

### Enrichment of let-7 targets in the LIN28B regulon

Predicted let-7 targets were obtained from TargetScan v8.0^130^ using cumulative weighted context++ scores. Enrichment of let-7 targets within the LIN28B (CDS) regulon inferred from the developing human brain atlas was assessed using a one-sided hypergeometric test, with 20,000 protein-coding genes as the background universe. Fold enrichment was calculated as the ratio of the observed to expected overlap. Robustness to background-universe definition was evaluated by repeating the analysis using background sizes ranging from 15,000 to 25,000 genes in increments of 1,000. The LIN28B–let-7 network included predicted let-7 targets overlapping the LIN28B (CDS) regulon and related components of the LIN28B–let-7 regulatory axis. TargetScan-predicted let-7–target interactions were weighted by cumulative context++ scores, with more negative values indicating stronger predicted repression. PubMed identifiers were annotated where literature evidence supporting the corresponding interactions was available.

### Statistics and reproducibility

Gene ontology biological-process enrichment of the 616 RBPs was performed using *enrichGO* in clusterProfiler^131^. Comparative pathway and disease-association analyses of RBPs and 1,390 TFs were conducted using Metascape^132^ and DisGeNET^133^. Functional enrichment of trait-relevant RBP regulons was assessed using Metascape^132^ and WebGestalt^134^. Trait-specific locus-to-gene (L2G) scores from the Open Targets platform^63^ were additionally analyzed by gene-set enrichment analysis. Single-cell transcriptomic data were analyzed using Scanpy (v1.9)^124^ and Seurat (v5.4.0)^100^. Pearson correlation was used to assess associations between motif-quality metrics and concordance of single-cell TRS profiles. Differences in RBP–target importance scores, computational time, transcript-region preferences and cell-identity benchmarking metrics were evaluated using Wilcoxon rank-sum tests. Fisher’s exact tests were used to assess overlap between the RBP and TF collections and differences in the proportion of neuropsychiatric disorder-associated regulons between developmental and non-developmental regulons.

This study comprised secondary analyses of publicly available GWAS summary statistics, gene-level rare-variant association data, and single-cell transcriptomic datasets; data sources and sample sizes are provided in Supplementary Note 7. No statistical methods were used to predetermine sample sizes. Cohort sizes for GWAS and rare-variant association analyses were determined by the originating studies, and all available single-cell samples passing the prespecified quality-control criteria were included. LD calculations for common-variant analyses used the 1,000 Genome Project Phase 3 reference panel^135^. Common-variant analyses were restricted to autosomal variants with minor allele frequency > 0.01, and the major histocompatibility complex region (chr6:25-35Mbp) was excluded because of its extensive LD.

## Funding

This study was supported by R01-MH137578, R01-MH123922, and R01-MH121521 (M.J.G.). The funders had no role in study design, data collection and analysis, decision to publish or preparation of the manuscript.

## Author Contributions

Y.M. and M.G. conceived and designed the study. Y.M. developed the methodology and implemented the software. Y.M., M.T., V.S., J.L., C.J., J.E., R.S., D.V., Y.L., and M.M. curated and collected data. Y.M., M.T., V.S., and J.L. performed the bioinformatics analyses and interpreted the data. Y.M., B.P., O.S., M.Z., D.V., C.V., L.T.U, and M.G. wrote the manuscript. All authors reviewed and approved the final version of the manuscript.

## Competing Interests

The authors declare no competing interests.

## Data availability

The GWAS summary statistics used in this study were obtained from publicly available resources, including the Psychiatric Genomic Consortium database (https://pgc.unc.edu/), the MRC IEU OpenGWAS database (https://opengwas.io/), FigShare (https://figshare.com/), and Zenodo (https://zenodo.org/). Human scRNA-seq datasets were also collected from public resources. These included 6.6K and 8.8K PBMC datasets from 10x Genomics (https://www.10xgenomics.com/), 17K PBMC from the COVID-19 cell atlas (https://www.covid19cellatlas.org/), human hematopoietic cell atlas (https://explore.data.humancellatlas.org/projects), a 59K PD midbrain single-cell dataset from the Gene Expression Omnibus (GEO; GSE193688), a 599K developing brain dataset from the UCSC Cell Browser (https://cells.ucsc.edu/?ds=dev-ctx-meta-atlas), a 232K developing brain dataset from the human developing brain scMultiome atlas (https://cell.ucsf.edu/snMultiome/), a 210K adult brain dataset from the European Genome-phenome Archive (EGA; EGAS00001005691), and 28 somatic tissue datasets from the Tabula Sapiens atlas (https://figshare.com/articles/dataset/Tabula_Sapiens_v2/27921984). A single-cell long-read sequencing (ScISOr-Seq) dataset for the developing human brain was obtained from the Gandal Lab GitHub repository (https://github.com/gandallab/Dev_Brain_IsoSeq/). For HepG2, we analyzed two scRNA-seq datasets generated using Smart-seq2 (GSE150993) and DNBelab data (GSM5677000). For K562, we analyzed two scRNA datasets generated using CEL-se (GSE1599500) and STORM-seq (GSE181544). Processed K562 perturb-seq data was downloaded from https://zenodo.org/records/17174974. Summary of GWAS summary statistics, gene-level rare-variant association data, and single-cell transcriptomic datasets used in this study is provided in Supplementary Tables 2, 3 and 40.

## Code availability

The scRBP Python package, an open-source software, has been developed and is freely available in the Github repository (https://github.com/mayunlong89/scRBP). The code to reproduce the results is accessible on the Github repository (https://github.com/mayunlong89/scRBP_reproduce).

## Notes

### Competing Interest Statement

The authors have declared no competing interest.

