## Supplementary Figures for "scRBP infers cell-specific RNA-binding protein regulons from single-cell transcriptomes at gene and isoform resolution"

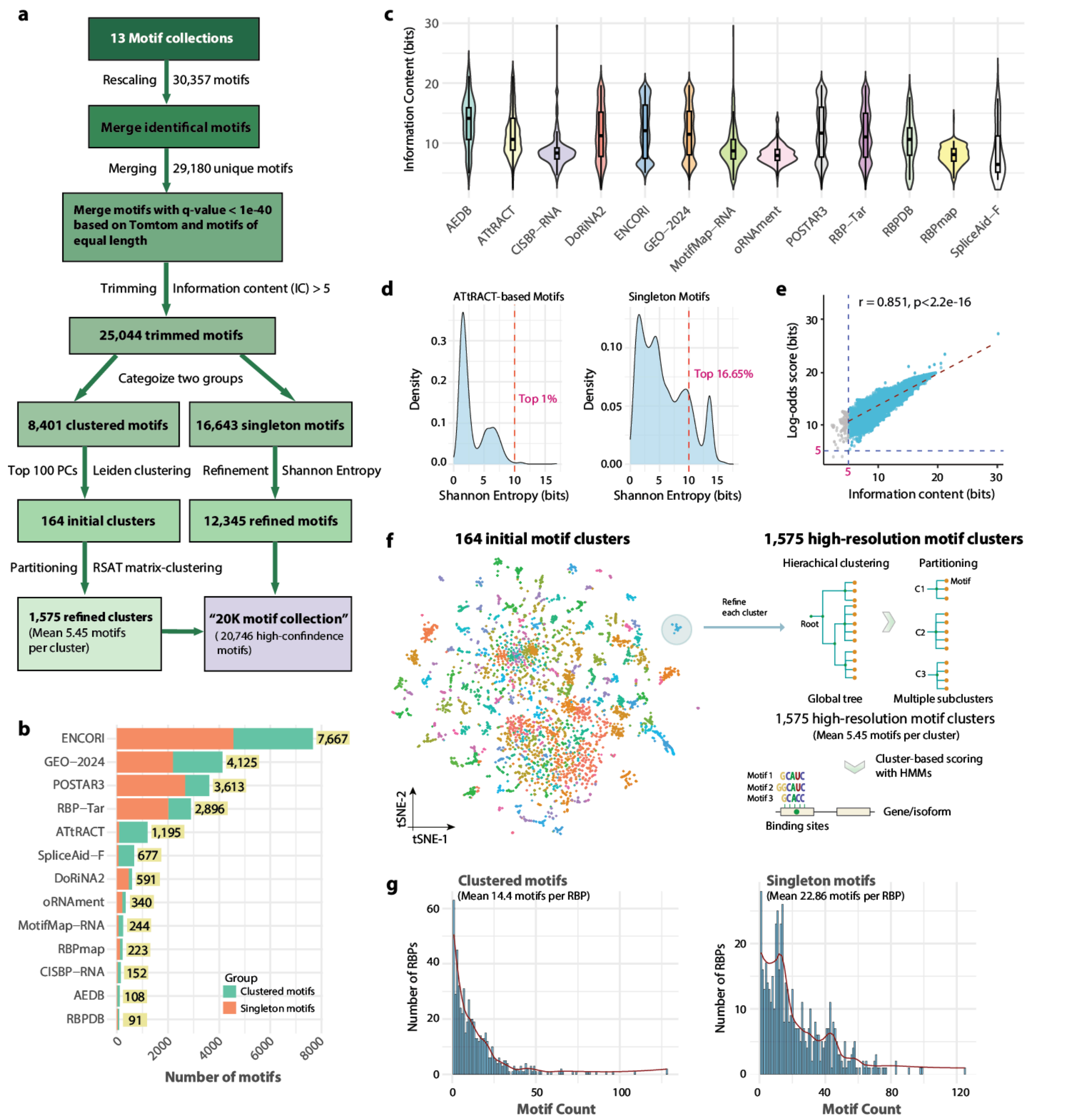


**Supplementary Fig. 1. Systematic collection and refinement of RNA-binding protein (RBP) motifs from public resources.**

(a) Overview of the motif curation and refinement workflow. In total, 30,357 motifs were collected from 13 public resources and rescaled using a unified normalization scheme. Motifs were iteratively merged and filtered to yield 8,401 clustered motifs and 12,345 singleton motifs, forming the final "20K motif collection" comprising 20,746 high-confidence motifs.
(b) Number of motifs contributed by each source, stratified into clustered and singleton motifs (colors). "GEO-2024" denotes motifs curated from 26 recently published CLIP-seq datasets retrieved from the GEO repository.
(c) Distributions of motif information content (IC) across the 13 source databases.
(d) Distribution of Shannon entropy for motifs from ATtRACT, CISBP-RNA, and RBPDB (left) and singleton motifs (right). High-entropy motifs (H > 10) are indicated by red dash lines.
(e) Pearson correlation between information content and HOMER-derived log-odds scores across all motifs (Pearson's r = 0.851, p-value < $2.2 \times 10^{-16}$).
(f) Two-step motif clustering strategy. Left, a t-distributed Stochastic Neighbor Embedding (t-SNE) visualization of 164 initial motif clusters derived from the 8,401 clustered motifs, based on motif-to-motif similarity. Right, hierarchical clustering applied within each initial cluster to resolve fine-grained structure, yielding 1,575 high-resolution motif subclusters.
(g) Distribution of motif counts per RBP. Number of motifs associated with each RBP in the clustered (left) and singleton (right) motif sets. RBPs in the clustered group harbor an average of 14.4 motifs per RBP; singleton motifs show a higher average of 22.86 motifs per RBP.

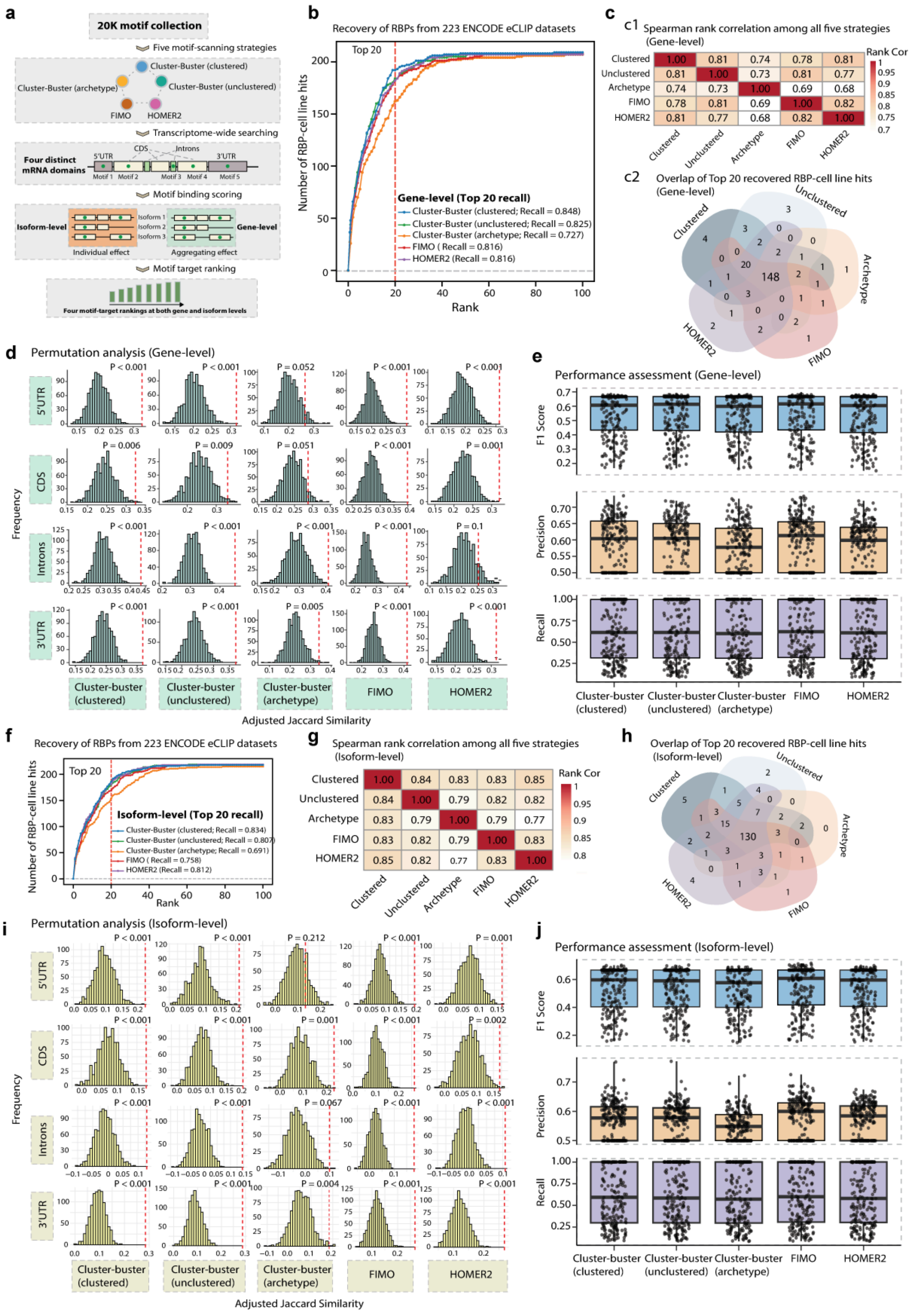


**Supplementary Fig. 2. Benchmarking motif-scanning strategies for transcriptome-wide RBP target**

**ranking**.

(a) Schematic overview of the motif-scanning framework. Motifs from the “20K motif collection” were scanned transcriptome-wide using five strategies: Cluster-Buster (clustered), Cluster-Buster (unclustered), Cluster-Buster (archetype), FIMO and HOMER2. Scans were performed across four transcript regions (5′UTR, CDS, intronic, and 3′UTR) at both gene and isoform resolution. Motif-derived binding scores were converted into motif-target rankings, yielding region-specific ranking resources at the gene and isoform levels.

(b) Gene-level RBP recovery using ENCODE eCLIP benchmarks (223 RBP-cell line datasets). Recall of matched RBP-cell line pairs among top-ranked predicted target genes is shown. Cluster-Buster (clustered) achieved the highest top-20 recall (0.848), followed by Cluster-Buster (unclustered; 0.825) and FIMO/HOMER2 (both 0.816).

(c) Concordance across scanning strategies at the gene level. (c1) Pairwise Spearman correlations between gene-level target rankings. (c2) Overlap of the top 20 recovered RBP–cell line hits across the five methods.

(d) Cross-cell-line reproducibility at the gene level assessed by adjusted Jaccard similarity. For 73 RBPs profiled in both HepG2 and K562, adjusted Jaccard similarity indices (JSIs) were computed for predicted target sets across four transcript regions. Null distributions were generated by 1,000 permutations; dashed red lines indicate observed means, and empirical P values are shown.

(e) Gene-level performance across methods using ENCODE gold-standard RBP–target gene sets. Metrics shown are F1 score (top), precision (middle) and recall (bottom).

(f) Isoform-level RBP recovery using ENCODE eCLIP benchmarks (223 RBP–cell line datasets), evaluated among top-ranked predicted target isoforms. Cluster-Buster (clustered) achieved the highest recall (0.834), followed by Cluster-Buster (unclustered; 0.807), HOMER2 (0.812) and FIMO (0.758).

(g) Pairwise Spearman correlations between isoform-level target rankings across the five scanning strategies.

(h) Overlap of the top 20 recovered RBP–cell line hits across the five methods at the isoform level.

(i) Cross-cell-line reproducibility at the isoform level assessed by adjusted Jaccard similarity. Observed JSIs between HepG2 and K562 are compared with null distributions from 1,000 permutations across four transcript regions (dashed red lines indicate observed means).

(j) Isoform-level performance across methods using ENCODE benchmark RBP–target isoform sets. Metrics shown are F1 score (top), precision (middle) and recall (bottom).

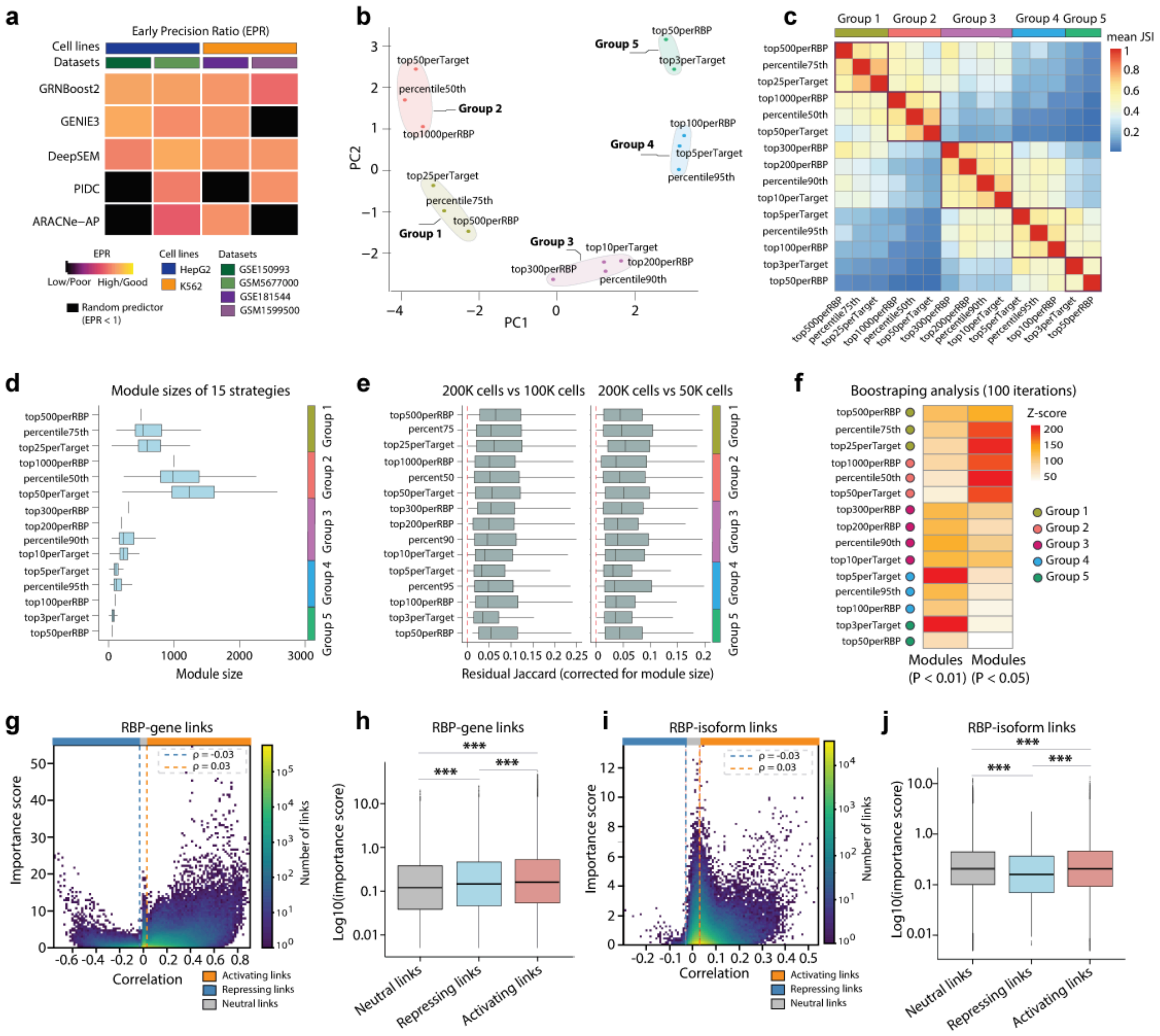


**Supplementary Fig. 3. Evaluation of RBP module construction strategies.**

(a) Heatmap of early precision ratio (EPR) for five GRN inference methods (GRNBoost2, DeepSEM, GENIE3, PIDC, and ARACNe-AP) across four scRNA-seq datasets and two cell lines (HepG2 and K562), evaluated against two independent ENCODE eCLIP-derived reference sets. Rows indicate methods and columns indicate dataset–cell line combinations. Values are EPRs (EPR = 1 denotes random expectation); black tiles indicate EPR < 1. GRNBoost2, GENIE3, and DeepSEM achieved superior performance for constructing RBP-mediated networks. Accordingly, scRBP uses the gradient boosting-based GRNBoost2 by default and supports GENIE3 as an alternative inference option.

(b) Principal component analysis (PCA) of 15 module construction strategies, showing distinct group separation along the first two principal components (i.e., PC1 and PC2).

(c) Unsupervised clustering of 15 RBP-module construction strategies using their mean Jaccard similarity indices (JSIs) across RBPs. Strategies partition into five groups (Groups 1–5) reflecting distinct design choices (e.g., top-*N* ranking versus percentile thresholding; gene-centric versus RBP-centric selection).

(d) Module size distributions for each strategy. Box plots summarize module sizes across RBPs (center line, median; box, IQR; whiskers, 1.5×IQR). Minima and maxima are represented by the whiskers.

(e) Cross-subsampling module preservation assessed by residual Jaccard similarity (adjusted for module size), comparing modules inferred from 200K cells to those inferred from 100K cells (left) or 50K cells (right). Positive residual Jaccard indicates greater-than-expected preservation given module size.

(f) Bootstrap-based assessment of strategy robustness. For each strategy, we quantified the enrichment of recovered modules among “significant” RBP modules from real data (defined by higher connectivity than label-shuffled null; 100 shuffles; P < 0.05 or P <

0.01). Z-scores were computed relative to overlaps obtained from 100 randomly generated strategies; points are colored by strategy group.

(g) Assigning regulatory signs to RBP–gene links. Scatter plot shows GRNBoost2 edge importance versus RBP–target Spearman correlation across cells. Vertical dashed lines denote correlation thresholds used to classify links as activating (r ≥ 0.03), repressing (r ≤ −0.03), or neutral; the horizontal dashed line indicates the minimum retained importance.

(h) Boxplot of RBP-gene importance score ($\log_{10}$ scale) by sign class. Both activating and repressing links show higher importance than neutral links (***P < 0.001, Wilcoxon signed-rank test).

(i) Assigning regulatory signs to all inferred RBP–isoform links. Density-colored scatter plot shows GRNBoost2 edge importance versus RBP–isoform Spearman correlation across all links. Vertical dashed lines denote the correlation thresholds used to classify links as activating (r ≥ 0.03), repressing (r ≤ −0.03), or neutral.

(j) Boxplot of RBP–isoform importance scores ($\log_{10}$ scale) across activating, neutral, and repressing link classes defined in panel (i). Activating and repressing links show significantly higher importance than neutral links (***P < 0.001, Wilcoxon signed-rank test).

Note: Data for panels (b–h) were derived from an adult brain single-cell RNA-seq dataset (Dataset ID: 210K_adult_brain; *n* = 209,801 cells), with specific analyses performed on relevant subsets where indicated. Data for panels (i–j) were derived from a developing brain single-cell long-read sequencing dataset (Dataset ID: IsoT7K_developing_brain; *n* = 7,189 cells), with specific analyses performed on relevant subsets where indicated. See Supplementary Table 3 for dataset details. For all boxplots, the center line indicates the median, boxes indicate the interquartile range (IQR), and whiskers extend to 1.5 × IQR.

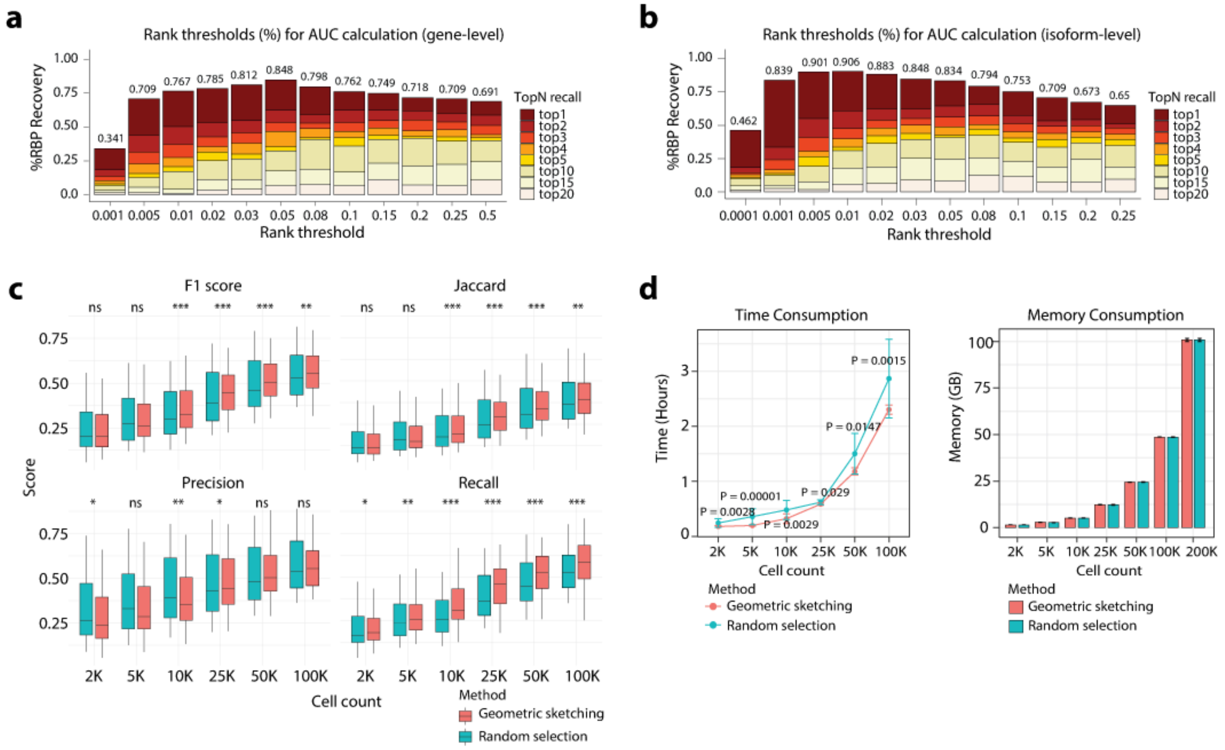


### Supplementary Fig. 4. Assessment of motif-enriched RBP-target interactions across different thresholds

(a) Rank-threshold sensitivity for gene-level AUC calculation. Stacked bars show the fraction of RBPs recovered across rank thresholds; colors indicate contributions from top-*N* recall (top1–top20). Numbers above bars denote total recovery.

(b) Rank-threshold sensitivity for isoform-level AUC calculation, shown as in (a).

(c) Benchmarking cell subsampling for RBP-module construction. Geometric sketching and random subsampling were evaluated at 2K, 5K, 10K, 25K, 50K and 100K cells using the 200K-cell analysis as reference. For each subset, we report distributions across RBPs for F1 score, Jaccard index, precision and recall (box, IQR; center line, median; whiskers, 1.5×IQR). Minima and maxima are represented by the whiskers. Significance was assessed by paired Wilcoxon tests (ns, not significant; * P < 0.05; ** P < 0.01; *** P < 0.001).

(d) Computational cost of geometric sketching versus random subsampling. Runtime (left) and peak memory (right) are shown across increasing cell counts.

Data for panels (d–f) were derived from an adult brain single-cell dataset (Dataset ID: 210K_adult_brain; n = 209,801 cells) reported by Palmer et al. (Supplementary Table 3), with specific analyses performed on relevant subsets where indicated.

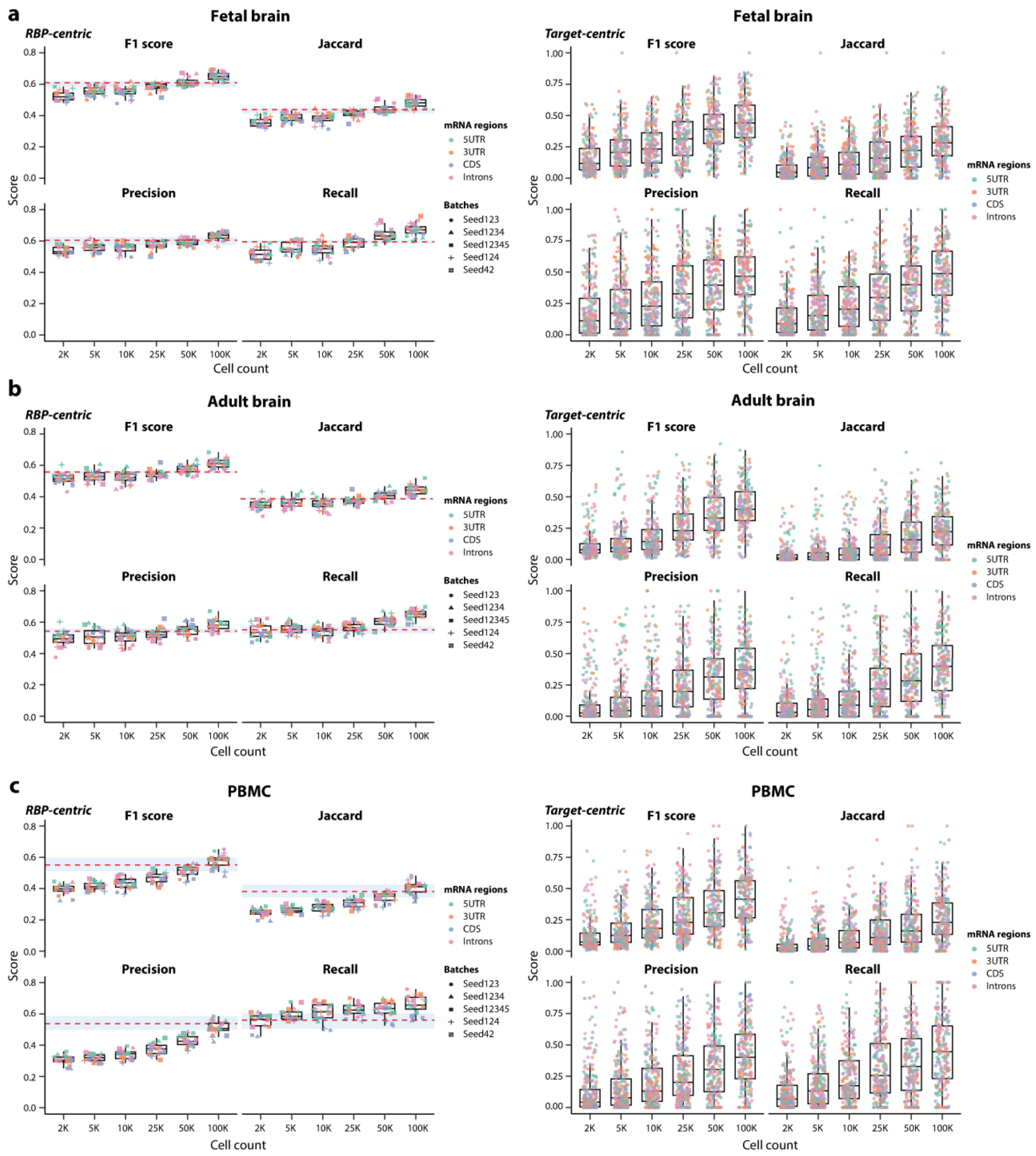


**Supplementary Fig. 5. The effects of cell number on RBP-regulon recovery across tissues and transcript regions.**

(a-c) Fetal brain (a), adult brain (b), PBMC (c). Subsampled datasets (2K, 5K, 10K, 25K, 50K and 100K cells) were benchmarked against a fixed reference (200K cells for fetal/adult brain; 170K cells for PBMC). For each cell-count setting, including the reference, datasets were generated in five independent replicates (random seeds: 123, 124, 1234, 12345, and 42). Reference RBP sets were defined as RBPs recovered in at least three of five reference replicates (≥3/5). Across tissues and transcript regions, performance metrics plateaued at approximately 50,000 cells, indicating stable regulon inference accuracy under moderate downsampling.

(1) RBP-centric evaluation (left panels). Overlap between RBPs recovered in each subsample and the reference RBP set was assessed across transcript regions (5′UTR, CDS, intronic, and 3′UTR) using F1 score, Jaccard index, precision and recall. Points are colored by region and shaped by random seed; boxplots summarize replicate distributions. The dashed line denotes the median baseline derived from seed-to-seed comparisons within the reference dataset (10 pairwise comparisons), and the shaded band indicates the range of region-specific medians. Boxes show the interquartile range with median center lines; whiskers extend to 1.5×IQR. Minima and maxima are represented by the whiskers.

(2) Target-centric evaluation (right panels). For RBPs shared between each subsample and the reference, agreement of target genes was evaluated within each RBP×region combination, treating reference regulon genes as positives. Metrics were computed per replicate and averaged across random seeds; each point represents one seed-averaged RBP×region pair. Boxplots summarize the distributions for the top 100 RBPs ranked by each metric (boxes, IQR; center line, median; whiskers, 1.5×IQR). Minima and maxima are represented by the whiskers.

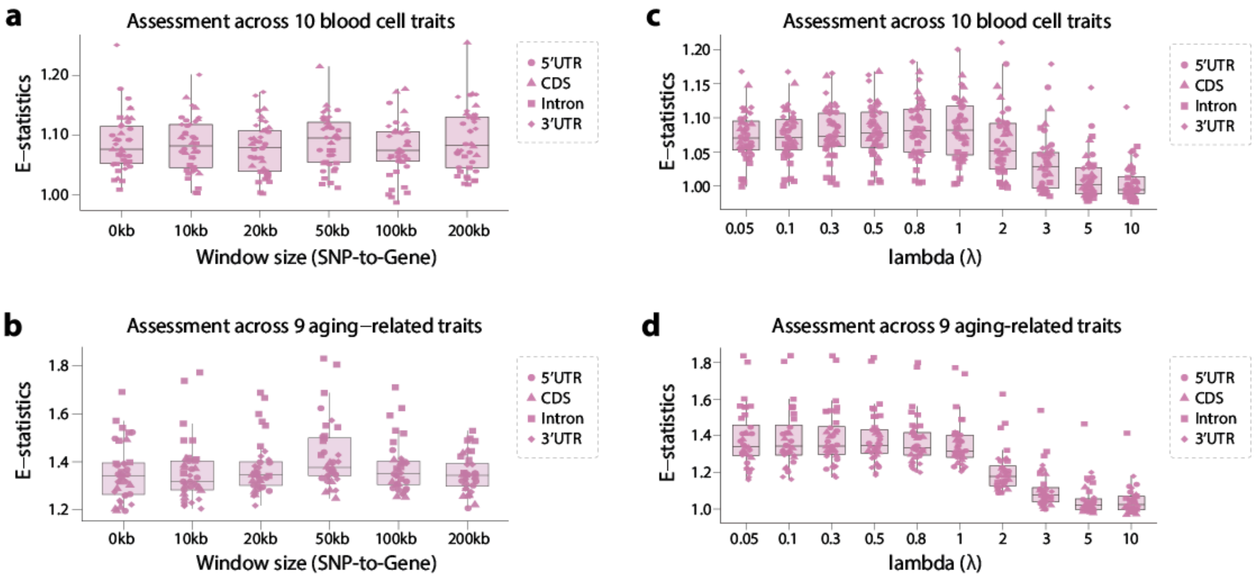


**Supplementary Fig. 6. Evaluation of scRBP performance under varying SNP-to-gene mapping windows and $\lambda$ parameters.**

(a) Distribution of E-statistics across ten hematological traits under different SNP-to-gene mapping windows (0 kb, 10 kb, 20 kb, 50 kb, 100 kb, and 200 kb). Results are shown for motif-target interactions located in four transcript regions: 5′UTR, coding sequence (CDS), intronic, and 3′UTR. Boxplots summarize the variation in enrichment statistics across ten hematological traits (Supplementary Table 2), with individual points representing trait-specific values. Analyses were performed using a peripheral blood mononuclear cell (PBMC) single-cell dataset (n = 6,593 cells; 3,313 monocytes and 3,280 T cells; Supplementary Table 3).

(b) Distribution of E-statistics across nine aging-related traits under varying SNP-to-gene mapping windows (0–200kb). Boxplots exhibit the variation in enrichment statistics across nine aging-related traits (Supplementary Table 2), with individual points representing trait-specific values across our transcript regions: 5′UTR, coding sequence (CDS), intronic, and 3′UTR. Analyses were conducted using a human midbrain single-cell dataset (n = 58,949 cells; Supplementary Table 3). E-statistics were calculated by comparing regulon activity enrichment between neurons and oligodendrocytes.

(c) Sensitivity of scRBP performance to the λ parameter controlling the integration of cell-type–specific regulon activity with genetic association signals. E-statistics were evaluated across ten hematological traits for four transcript regions (5′UTR, CDS, intronic, and 3′UTR) using the PBMC dataset.

(d) Sensitivity of scRBP performance to the λ parameter for nine aging-related traits based on the midbrain single-cell dataset. E-statistics were calculated analogously to panel (b), comparing enrichment between neurons and oligodendrocytes across transcript regions. For all panels, boxplots show the median (center line), interquartile range (box), and whiskers extending to 1.5× the interquartile range. Individual points denote trait-specific results.

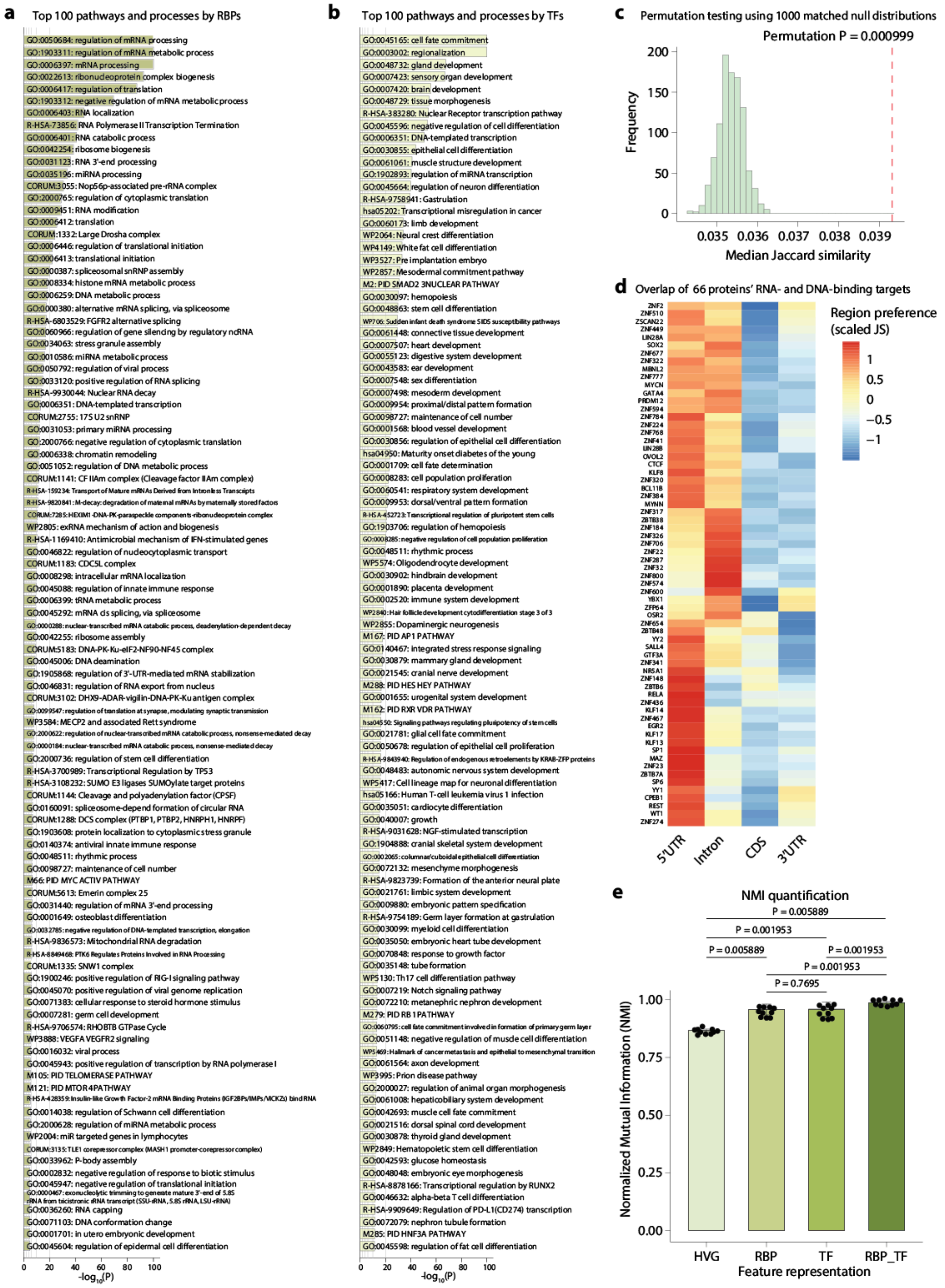


**Supplementary Fig. 7. Expanded functional enrichment landscape of RBPs, TFs, and dual-binding proteins.**

(a) Top 100 enriched pathways and biological processes for the 616 RBPs included in the scRBP motif collection. Functional enrichment analysis was performed using Metascape, and terms were ranked by -log$_{10}$(P value).

(b) Top 100 enriched pathways and biological processes for 1,390 TFs curated from the pySCENIC motif collection. Enriched terms

were ranked by -$\log_{10}$(P value) and visualized across functional categories.
(c) Permutation-based assessment of the global similarity between TF- and RBP-derived regulatory targets across dual RNA- and DNA-binding proteins. The null distribution of median Jaccard similarity was generated using 1,000 matched permutations, in which protein labels were randomized while preserving the number of targets per protein. For each permutation, motif-specific rankings were aggregated using the minimum-rank strategy and the top 1,500 targets per protein were retained to compute pairwise Jaccard similarity. The red dashed line indicates the observed median Jaccard similarity across the 66 dual-binding proteins.
(d) Overlap between TF-derived and RBP-derived regulatory targets among 66 dual RNA- and DNA-binding proteins. The heatmap displays row-scaled Jaccard similarity between TF motif targets and RBP motif targets derived from 5′UTR, intronic, CDS and 3′UTR motif rankings. TF motif target rankings were obtained from the *hg38__refseq-r80__10kb_up_and_down_tss.mc9nr.genes_vs_motifs.rankings.feather*. Because each TF or RBP can be associated with multiple motifs, motif-specific rankings were first integrated using minimum-rank aggregation, and the top 1,500 targets per protein were retained for similarity analysis.
(e) NMI benchmarking of feature representations across eight ENCODE cell lines. Simulated pseudocells (n = 1,600; 200 cells per cell line) were generated from bulk RNA-seq profiles of eight deeply profiled ENCODE cell lines. Clustering accuracy was evaluated using Normalized Mutual Information (NMI) for feature representations derived from the top 2,000 highly variable genes (HVGs), RBP regulons, TF regulons, and the combined RBP–TF regulons. Clustering was performed across 10 independent replicates for each feature representation. Bars represent mean NMI values, error bars denote standard deviation, and dots indicate individual replicates (paired Wilcoxon signed-rank test; P values indicated).
.

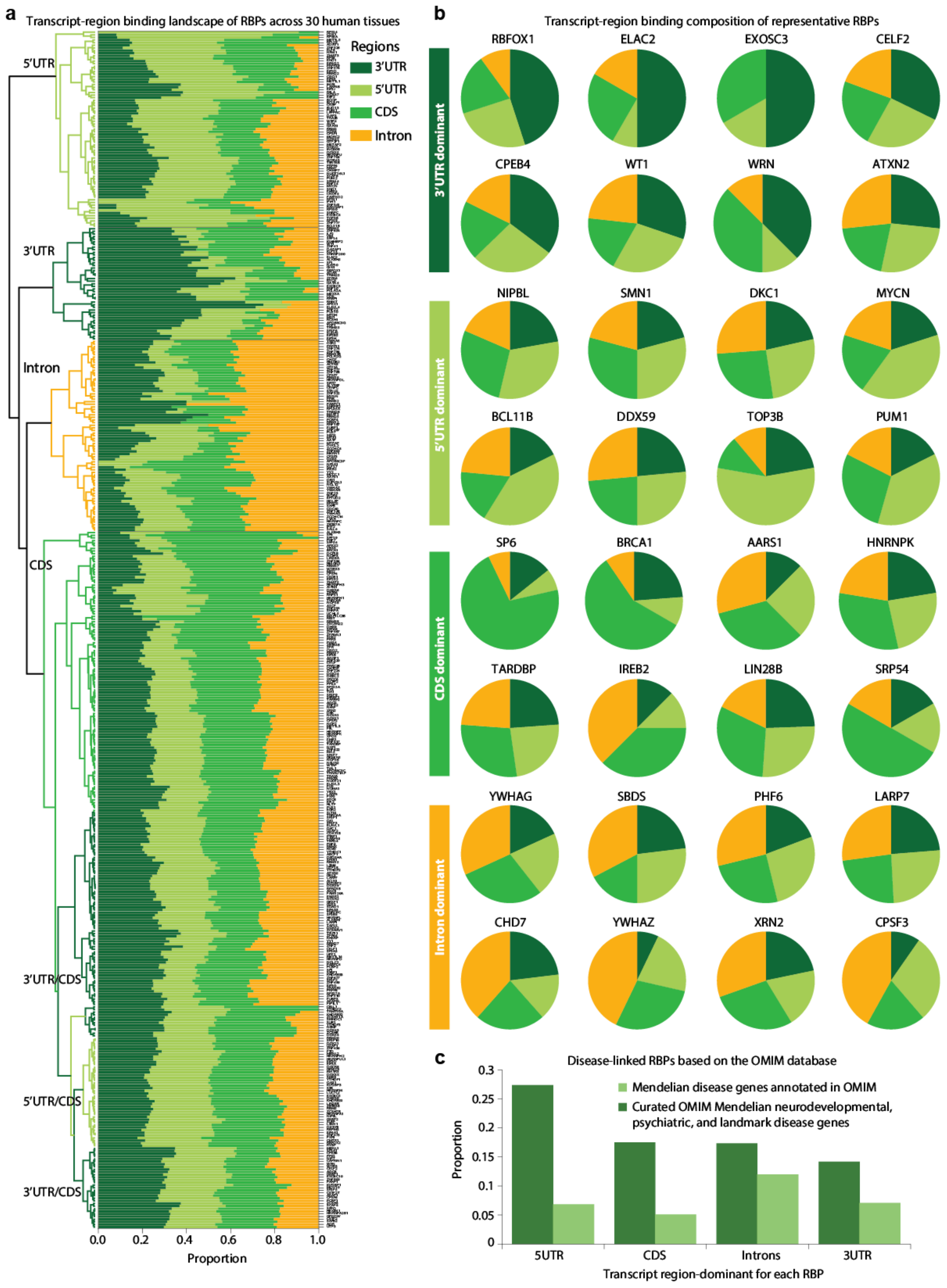


**Supplementary Fig. 8. Global landscape of RBP transcript-region binding preferences across 30 human tissues.**

(a) Stacked bar plot summarizing the transcript-region binding composition of RBPs across 30 human tissues. For each tissue, scRBP-derived region-specific regulon gene sets were defined for four transcript regions (3′UTR, 5′UTR, CDS, and intronic). For each RBP within each tissue, enrichment in a given transcript region was recorded as present or absent based on the scRBP pruning results. These enrichment events were aggregated across tissues to quantify the number of tissues in which each RBP exhibited significant enrichment in each transcript region. The stacked bars represent the fraction of region-enriched occurrences for each RBP across tissues, with proportions normalized so that the four transcript regions sum to one. RBPs are grouped according to their dominant transcript region (defined as the region with the highest composition fraction). Within each group, RBPs are hierarchically clustered based on their transcript-region composition profiles across tissues and ordered according to the resulting dendrogram structure. Only RBPs showing enrichment in at least five tissues were included. The 30 tissues analyzed include fetal brain, adult brain, liver, vasculature, fat, testis, large intestine, small intestine, bladder, lymph node, blood, bone marrow, ear, eye, kidney, heart, salivary gland, mammary, muscle, ovary, pancreas, prostate, skin, spleen, thymus, stomach, tongue, uterus, trachea, and lung (Supplementary Table 3).

(b) Pie charts illustrating transcript-region binding compositions for 32 representative RBPs. For each RBP, the proportions were calculated using the same cross-tissue aggregation strategy described in panel (a). Representative RBPs were selected to highlight distinct region-dominant classes as well as RBPs exhibiting mixed or multi-region binding preferences across tissues.

(c) Distribution of human RBPs associated with Mendelian diseases based on annotations from the Online Mendelian Inheritance in Man (OMIM) database. Genes were classified according to their known disease relevance: 1) light green: Mendelian disease genes annotated in OMIM; 2) dark green: Curated OMIM Mendelian disease genes relevant to neurodevelopment, psychiatry, and key human genetic disorders. Bar plots summarize the number of RBPs within each region-dominant class (3′UTR-, 5′UTR-, CDS-, and intronic-dominant) stratified by OMIM disease rank, providing an overview of the relationship between transcript-region binding preferences and genetic disease relevance among human RBPs.

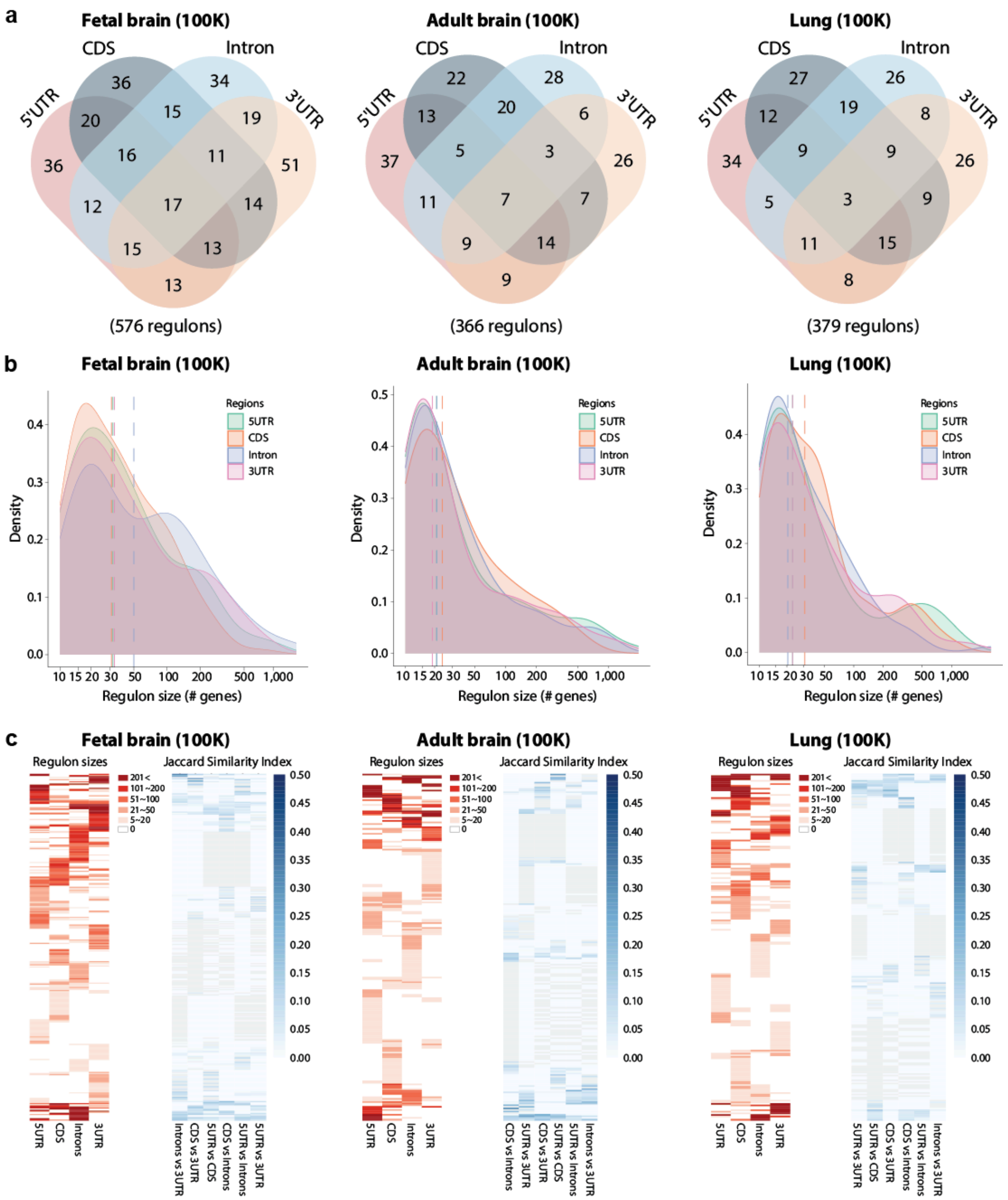


### Supplementary Fig. 9. Tissue- and transcript-region-resolved properties of scRBP regulons.

(a) Four-way Venn diagrams showing overlap of scRBP regulons inferred from four transcript regions (5′UTR, CDS, intronic, and 3′UTR) in fetal brain, adult brain and lung (each downsampled to $n$ = 100,000 cells). Numbers denote counts of RBPs with an inferred regulon in the indicated region combination; totals below each diagram indicate the number of region-specific regulons recovered in that tissue.

(b) Distributions of regulon sizes (number of target genes) stratified by transcript region in fetal brain, adult brain and lung. Curves show kernel density estimates for each region; dashed vertical lines indicate region-specific medians.

(c) Transcript-region “fingerprints” of inferred regulons in fetal brain, adult brain and PBMC. Left, heatmaps show regulon size for each RBP across the four regions. Right, heatmaps show pairwise Jaccard similarity among the corresponding region-specific target sets for the same RBPs, quantifying within-RBP consistency across transcript regions (darker indicates higher similarity).

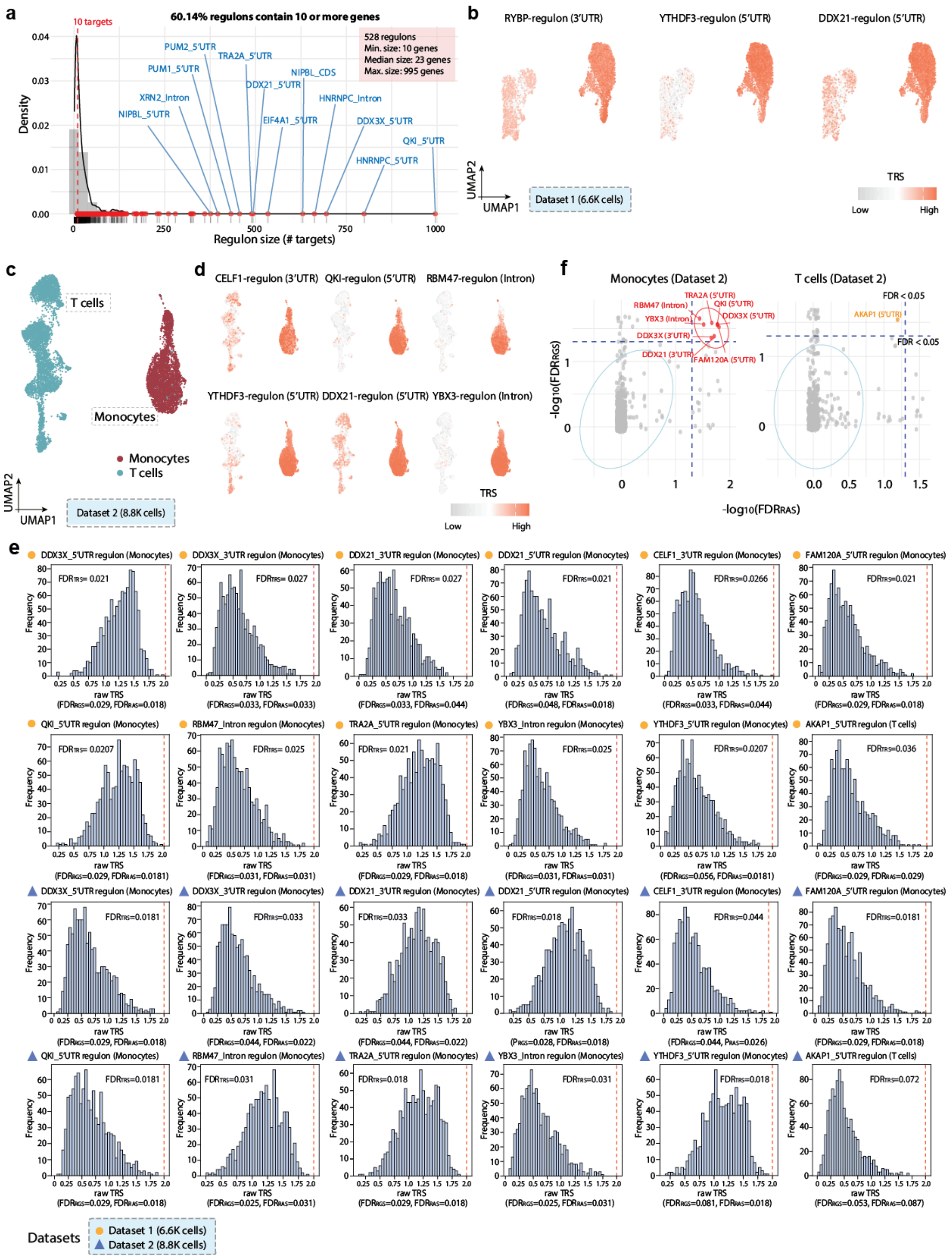


**Supplementary Fig.10. Robustness of scRBP in mapping monocyte count-associated regulons across PBMC datasets and transcript regions.**

(a) Distribution of regulon sizes (number of target genes) in PBMC Dataset 1 (n = 6.6K cells). Regulons with 10 or more targets (60% of all regulons) were retained for downstream analysis.

(b) Representative examples illustrating cell type specificity in discovery Dataset 1. Three monocyte count-associated RBP regulons (RYBP_3'UTR, YTHDF3_5'UTR, and DDX21_5'UTR) exhibit higher trait-relevance scores (TRSs) in monocytes than T cells.

(c) UMAP of PBMCs from validation dataset (Dataset 2: n = 8.8K cells), highlighting monocytes (red; n = 3,546 cells) and T cells (teal; n = 5,231 cells).

(d) Additional representative RBP regulons in Dataset 2. Six monocyte count-relevant RBP regulons (CELF1_3'UTR, QKI_5'UTR, RBM47_intronic, YTHDF3_5'UTR, DDX21_5'UTR, and YBX3_intronic) exhibit stronger TRSs in monocytes than in T cells.

(e) Empirical null distributions of TRSs for 12 selected regulons (DDX3X_5’UTR, DDX3X_3’UTR, DDX21_3’UTR, DDX21_5’UTR, CELF1_3’UTR, FAM120A_5’UTR, QKI_5’UTR, RBM47_intronic, TRA2A_5’UTR, YBX3_intronic, YTHDF3_5’UTR, and AKAP1_5’UTR), with observed TRSs from Dataset 1 (green) and Dataset 2 (orange) indicated.
(f) Scatter plots of scRBP-identified regulons associated with the monocyte count trait in monocytes (left) and T cells (right) in Dataset 2 (n = 8,777 cells). Each dot represents an RBP-regulon, with $-\log_{10}(FDR_{RGS})$ on the y axis and $-\log_{10}(FDR_{RAS})$ on the x axis. Gray dots denote nonsignificant regulons, and red dots indicate significant regulons. Statistical significance was assessed by an MC permutation test (one-sided, upper tail; 1,000 iterations; multiple-testing corrected).

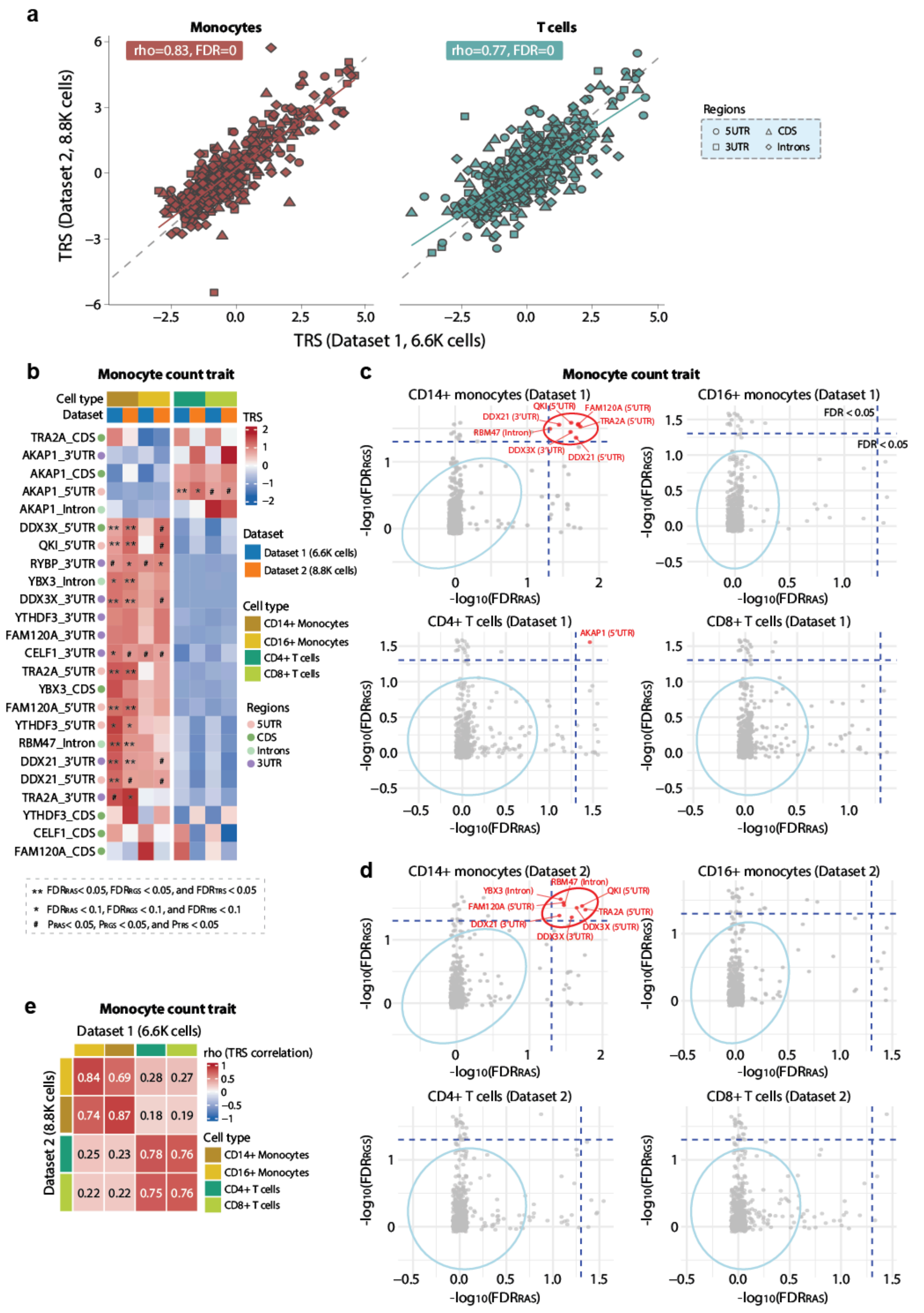


**Supplementary Fig. 11. Stratified scRBP analysis identifies subtype-specific regulons associated with monocyte count.**

(a) Cross-dataset concordance of regulon-level TRSs for the monocyte count trait. Scatter plots compare TRSs estimated in Dataset

1 (x-axis) and Dataset 2 (y-axis), shown separately for monocytes and T cells; Pearson correlation coefficients are indicated.
(b) Heatmap summarizing monocyte count-associated RBP-regulons across four immune cell subpopulations (CD14+ monocytes, CD16+ monocytes, CD4+ T cells, and CD8+ T cells) in the discovery dataset (Dataset 1; n = 6.6K cells) and an independent validation dataset (Dataset 2; n = 8.8K cells). ** indicates $FDR_{RAS} < 0.05$, $FDR_{RGS} < 0.05$, $FDR_{TRS} < 0.05$; * indicates $FDR < 0.1$ for all three scores; # indicates nominal significance ($P_{RAS}$, $P_{RGS}$, and $P_{TRS} < 0.05$).
(c,d) Scatter plots of scRBP trait-association results in each subpopulation for Dataset 1 (c) and Dataset 2 (d). Each dot denotes an RBP-regulon, with $-\log_{10}(FDR_{RGS})$ on the y axis and $-\log_{10}(FDR_{RAS})$ on the x axis. Dashed lines indicate FDR thresholds; highlighted points mark representative significant regulons. Significance was assessed by Monte Carlo permutation testing (one-sided upper tail; 1,000 permutations) with multiple-testing correction.
(e) Heatmap of Pearson correlations of regulon-level TRSs across immune subpopulations between the two PBMC datasets, demonstrating higher within-lineage concordance for myeloid and T-cell populations compared with cross-lineage comparisons.

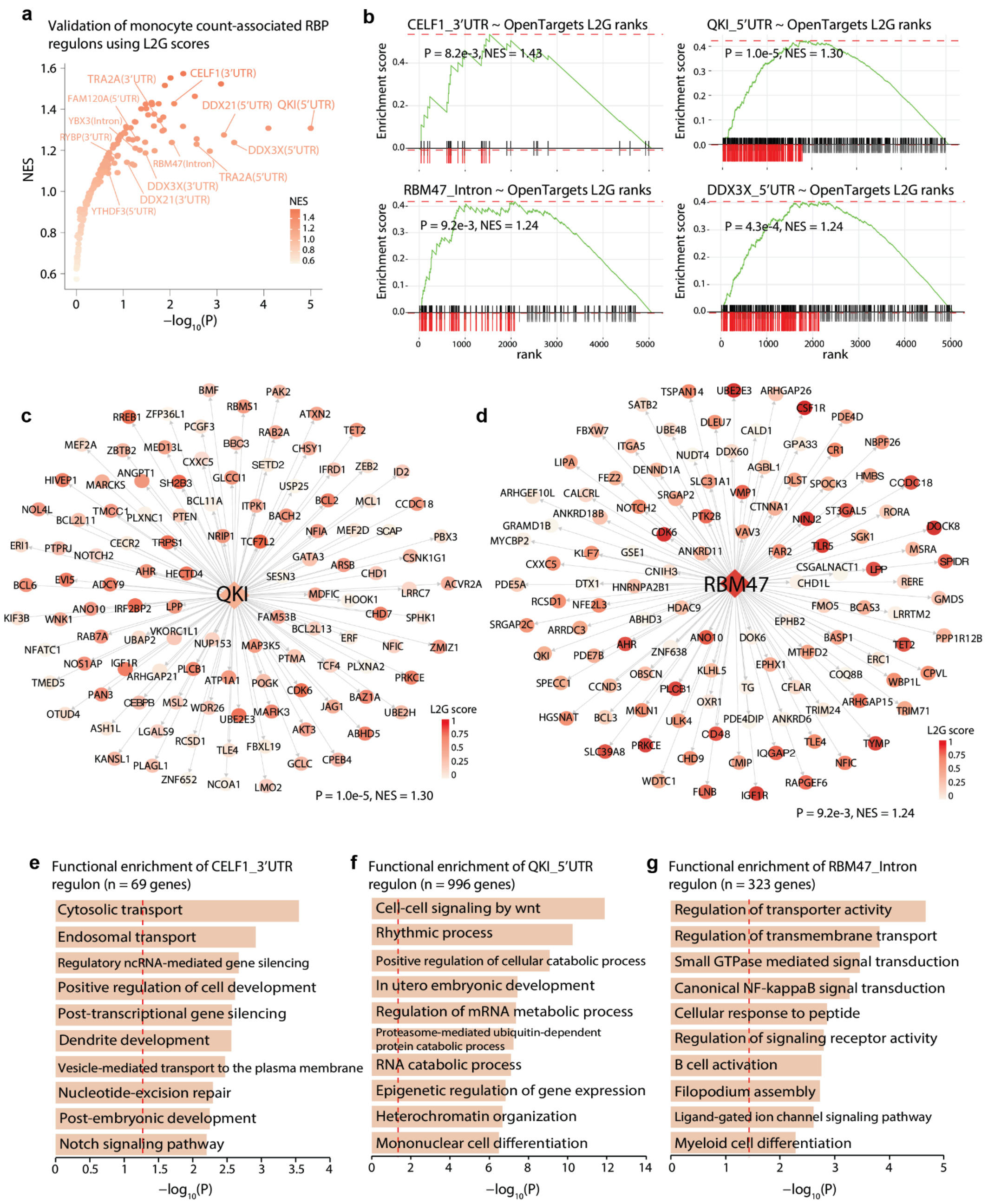


**Supplementary Fig. 12. Functional characterization and orthogonal validation of monocyte count-associated RBP-regulons.**

(a) Orthogonal validation of scRBP-prioritized regulons using Open Targets. Genes were ranked by locus-to-gene (L2G) scores for the monocyte count trait, and enrichment of each RBP regulon was evaluated by GSEA. The x-axis shows enrichment significance (-$\log_{10}$P), and the y-axis shows the normalized enrichment score (NES). scRBP-prioritized trait-relevant regulons (e.g., CELF1_3'UTR and QKI_5'UTR) are highlighted.

(b) GSEA of selected monocyte count-associated regulons using Open Targets locus-to-gene (L2G) scores. For each highlighted regulon (CELF1_3′UTR, QKI_5′UTR, DDX21_5′UTR, RBM47_intronic, TRA2A_5′UTR, DDX3X_5′UTR), genes were ranked by Open Targets monocyte count L2G score and tested for enrichment. Curves show the running enrichment score across the ranked list; tick marks denote regulon genes. Nominal *P* values and normalized enrichment scores (NES) are shown.

(c) Network representation of the monocyte-enriched QKI_5′UTR regulon (n = 996 genes). The regulator is shown as a diamond and target genes as circles; node color encodes Open Targets L2G score. The GSEA significance for this regulon is indicated.

(d) Network representation of the monocyte-enriched RBM47_intronic regulon (n = 323 genes), displayed as in (b).

(e)GO-BP enrichment for CELF1_3′UTR targets (top 10 terms shown), ranked by -$\log_{10}$(P) (WebGestalt).

(f) GO-BP enrichment for QKI_5′UTR targets (top 10 terms shown), displayed as in (d) (WebGestalt).
(g) GO-BP enrichment for RBM47_intronic targets (top 10 terms shown), displayed as in (d) (WebGestalt).

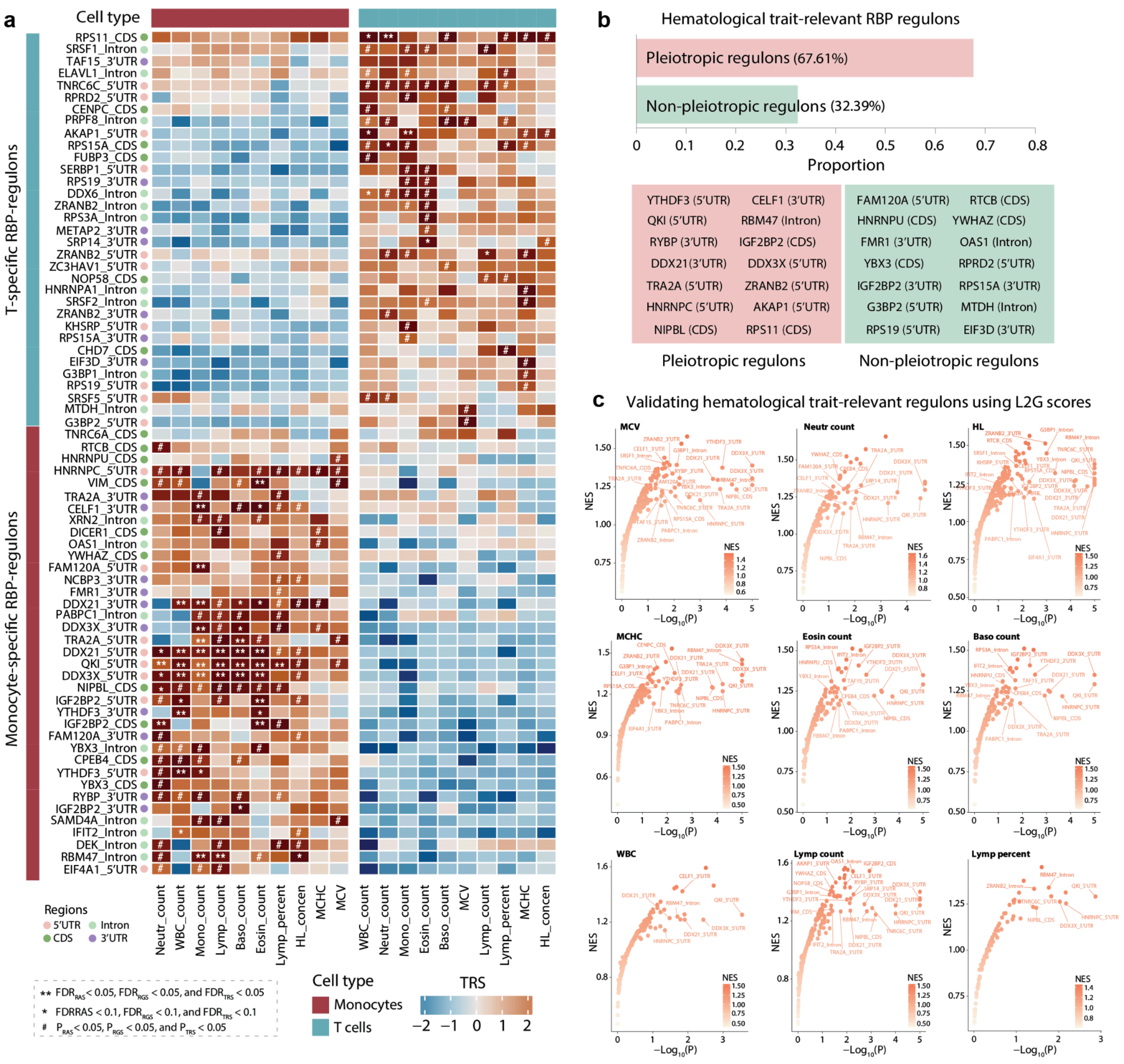


**Supplementary Fig. 13. scRBP identifies cell-type-specific RBP regulons linked to hematological traits and provides orthogonal genetic support.**

(a) Heatmap summarizing cell-type-specific, trait-associated RBP regulons across ten hematological traits in Dataset 1 (n = 6.6K cells). Tiles show regulon–trait association signals stratified by cell type. Significance annotations: ** indicates $FDR_{RAS} < 0.05$, $FDR_{RGS} < 0.05$, $FDR_{TRS} < 0.05$; * indicates FDR < 0.1 for all three scores; # indicates nominal significance ($P_{RAS}$, $P_{RGS}$, and $P_{TRS} < 0.05$).

(b) Proportion of pleiotropic regulons across ten hematological traits. Representative pleiotropic and non-pleiotropic regulons are shown in the bottom panel.

(c) Orthogonal validation using Open Targets of nine additional hematological traits. For each trait, genes were ranked by locus-to-gene (L2G) scores and tested for enrichment of each scRBP regulon using GSEA. Points summarize enrichment for regulons prioritized by scRBP (x axis, $-\log_{10}P$; y axis, normalized enrichment score (NES)).

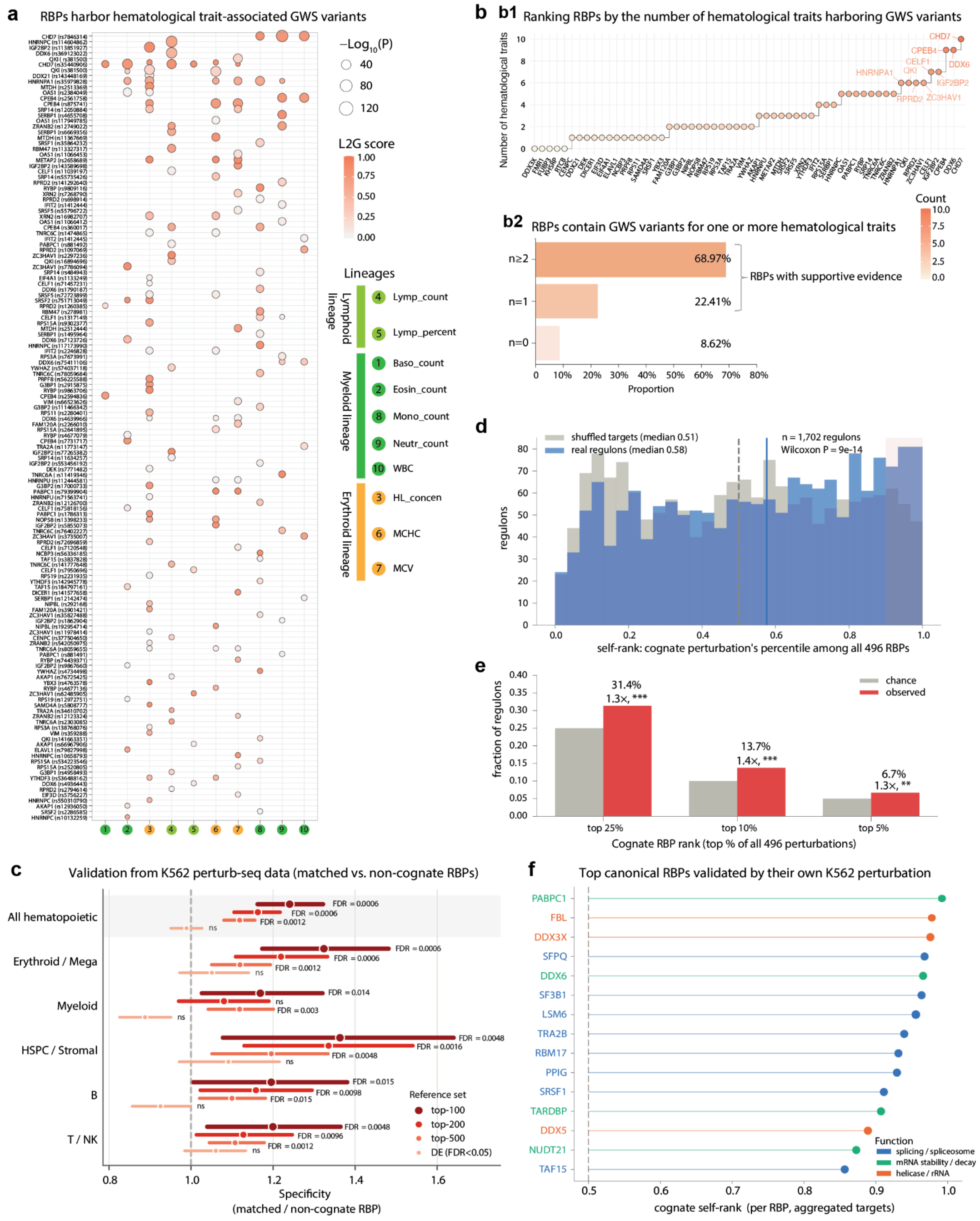


**Supplementary Fig. 14. Hematological trait-relevant scRBP regulons are supported by human genetics and cognate RBP perturbation.**

(a) Trait-by-trait overview of genome-wide significant (GWS) variants mapped to RBPs across the ten hematological traits. Each point denotes a variant–RBP assignment for a given trait; color encodes the L2G score and point size reflects GWAS significance.

(b) Burden of hematological trait associations across RBPs. (b1) RBPs ranked by the number of distinct traits with mapped GWS variants; labeled points highlight the top-ranked RBPs. (b2) Fractions of RBPs with GWS variants mapped to zero, one, or at least two traits (n = 0, n = 1, n ≥ 2); RBPs with n ≥ 1 are considered to have supportive genetic evidence.

(c) RBP-regulon targets respond preferentially to cognate RBP perturbation. scRBP regulons were inferred from an integrated Human Cell Atlas hematopoietic reference comprising 1,453,784 cells and evaluated against a genome-scale K562 CRISPRi Perturb-seq screen comprising 1,989,578 cells (Replogle et al., 2022). Specificity was defined as the pooled enrichment of regulon targets among genes responsive to perturbation of the cognate RBP divided by the corresponding enrichment across non-cognate RBP perturbations. Regulons, hematopoietic lineages and response-gene definitions are as in Fig. 3i. Points and error bars indicate pooled specificity ratios and 95% bootstrap confidence intervals, respectively; the dashed line indicates no cognate specificity (ratio = 1). Significance was assessed using 5,000 RBP-identity permutations followed by Benjamini–Hochberg correction; ns, FDR > 0.05.

(d) Distribution of cognate RBP rank percentiles ("self-ranks") across 1,702 regulons, obtained by ranking each regulon's target response across 496 RBP perturbations. Real regulons were shifted toward higher self-ranks relative to size-matched shuffled target sets (median, 0.58 versus 0.51; one-sided paired Wilcoxon signed-rank test, $P = 9 \times 10^{-14}$). Dashed line, chance expectation (0.5); solid blue line, median for real regulons; shading, top decile.

(e) Fraction of regulons whose cognate RBP ranked within the top 25%, 10% or 5% of all perturbations, compared with chance expectations. Cognate RBPs were enriched at all three thresholds (31.4%, 13.7% and 6.7%; 1.3–1.4-fold over chance; one-sided binomial test, $P = 2 \times 10^{-9}$, $5 \times 10^{-7}$ and $1 \times 10^{-3}$, respectively). ** $P < 0.01$; *** $P < 0.001$.

(f) Canonical RBPs are preferentially recovered by cognate perturbation. Cognate self-ranks for the 15 canonical RBPs with the highest values. For each RBP, targets were aggregated across its cell type- and transcript region-specific regulons and scored against 496 RBP perturbations in the K562 CRISPRi Perturb-seq dataset. Self-rank denotes the percentile rank of the cognate perturbation among all perturbations; the dashed line indicates chance expectation (0.5). All selected RBPs had self-ranks above 0.85. Points are coloured by the RNA-processing function.

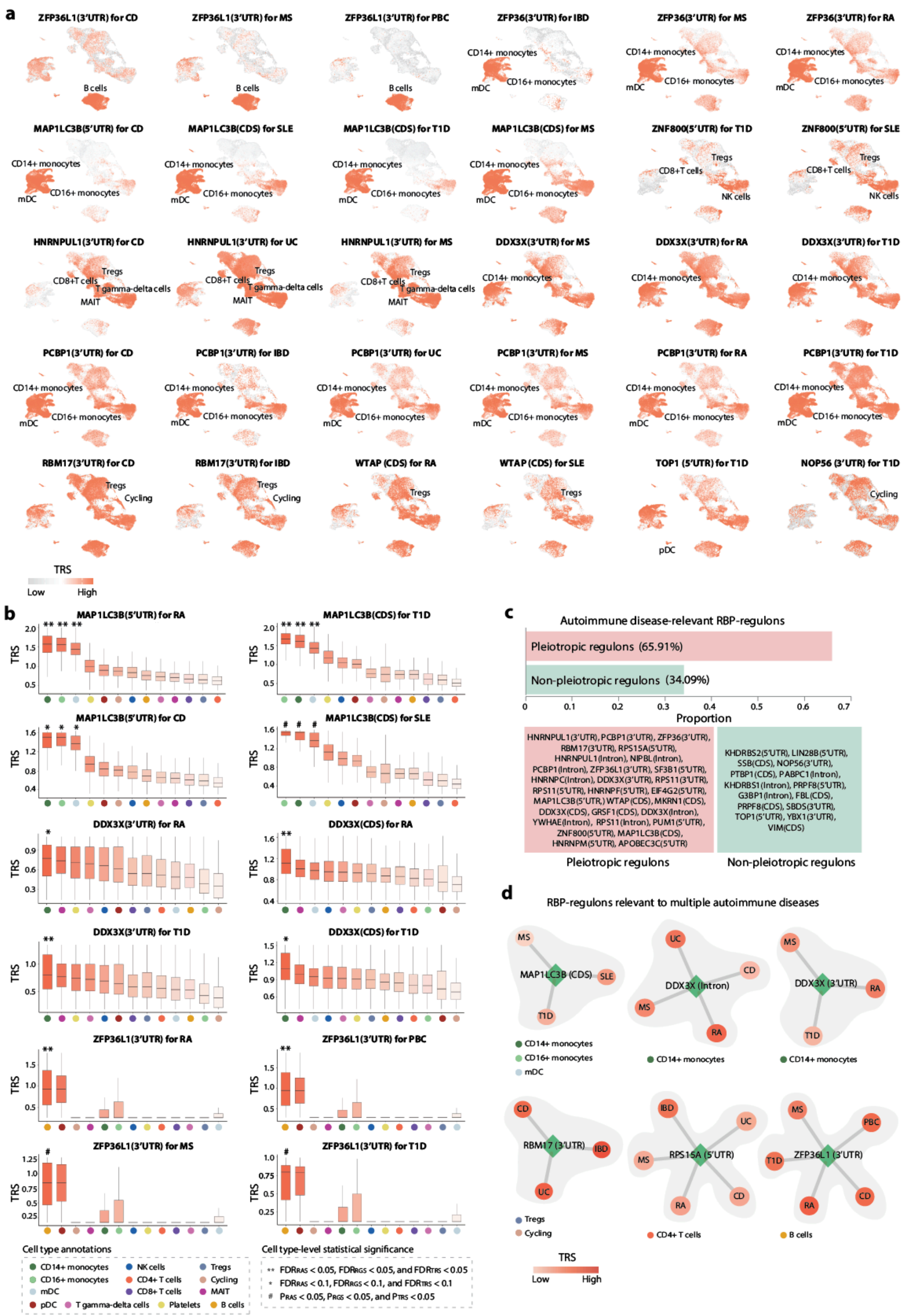

**Supplementary Fig. 15. Single-cell-resolved scRBP analysis highlights autoimmune-associated regulons enriched in high-TRS cells within their cognate immune populations.**

(a) Representative UMAP feature plots showing per-cell trait-relevance scores (TRSs) for autoimmune-associated RBP regulons. Examples illustrate strong cell type-specificity, with higher TRSs concentrated in the expected ("focal") immune populations relative to other cell types.

(b) Distributions of per-cell TRSs for autoimmune-associated regulons across immune cell types (boxplots; center line, median; box, interquartile range; whiskers, 1.5× IQR).

(c) Pleiotropy of scRBP-identified regulons across eight autoimmune diseases. The bar plot summarizes the fraction of regulons associated with two or more diseases (pleiotropic) versus those associated with a single disease (non-pleiotropic). Boxes list representative regulons from each category.

(d) Examples of regulon–disease sharing. Network diagrams depict six regulons associated with multiple autoimmune diseases, with nodes indicating diseases and edges indicating significant regulon–trait links; node color encodes TRS.

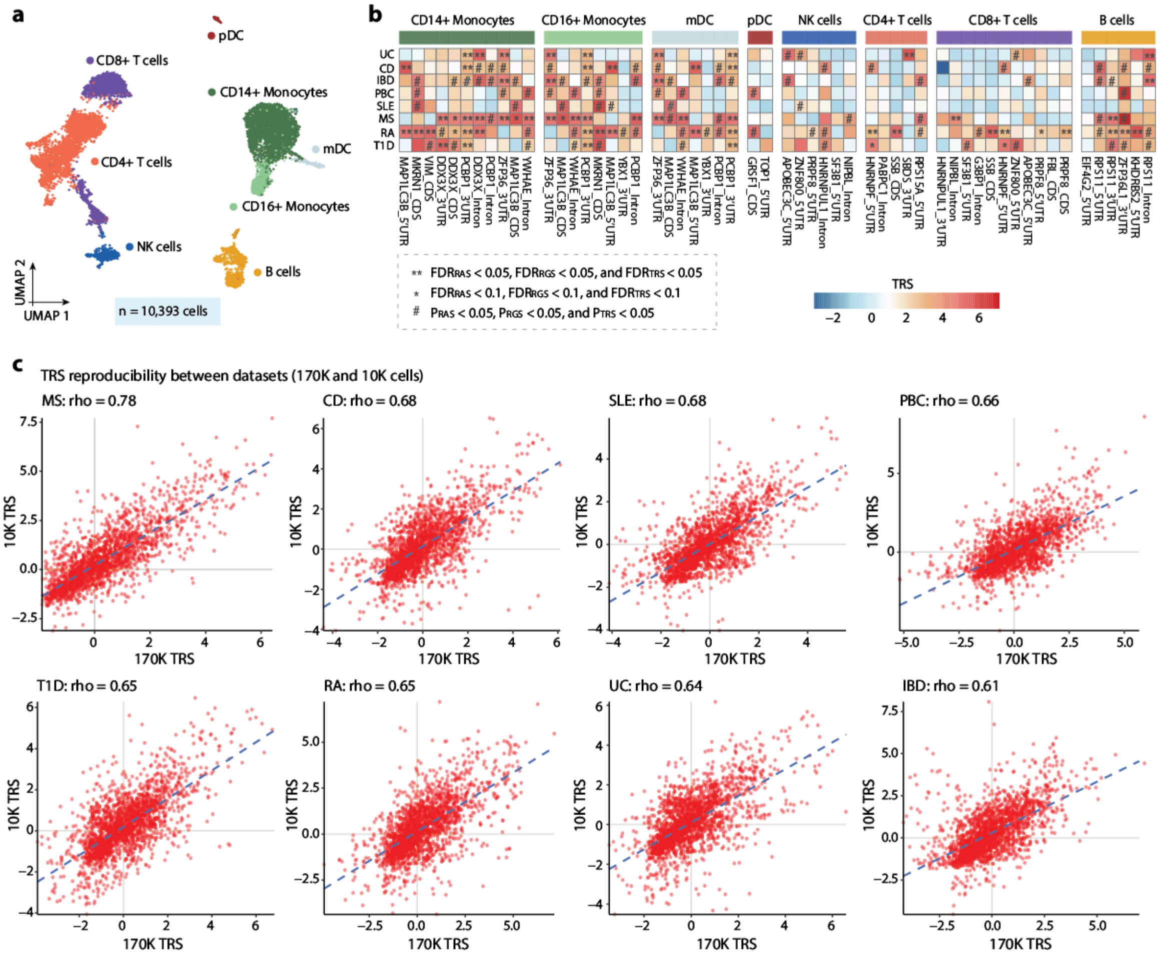


**Supplementary Fig. 16. Replication of scRBP-identified autoimmune-associated regulons in an independent PBMC scRNA-seq dataset.**

(a) UMAP of PBMC from the replication dataset, colored by annotated immune cell types (n = 10,393 cells; Supplementary Table 3).

(b) Summary heatmap of cell type-resolved RBP-regulons associated with eight autoimmune diseases across eight immune populations. Significance is reported for three complementary tests (RAS, RGS and TRS). ** indicates $FDR_{RAS} < 0.05$, $FDR_{RGS} < 0.05$, $FDR_{TRS} < 0.05$; * indicates FDR < 0.1 for all three scores; # indicates nominal significance ($P_{RAS}$, $P_{RGS}$, and $P_{TRS} < 0.05$). P values were obtained by Monte Carlo permutation (one-sided, upper tail; 1,000 iterations) and adjusted for multiple testing.

(c) Cross-dataset concordance of regulon-level TRSs for the eight autoimmune diseases. Scatter plots compare TRSs estimated in the discovery PBMC dataset (170K cells; x-axis) with those from the replication dataset (10K cells; y-axis). Pearson correlation coefficient is shown in each panel.

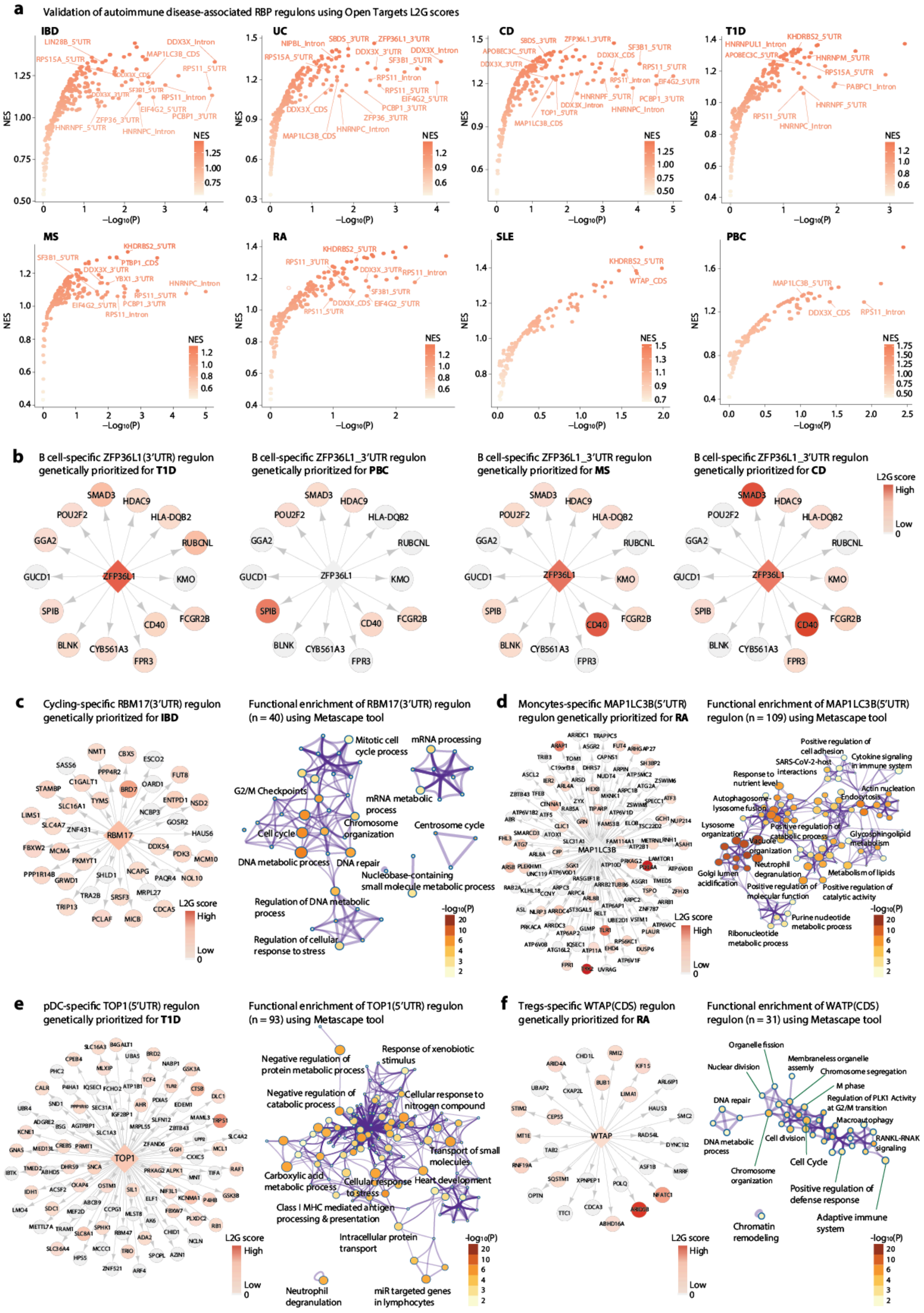

**Supplementary Fig. 17. Orthogonal genetic support for scRBP-identified regulons for eight autoimmune diseases.**

(a) Enrichment of scRBP-prioritized regulons in Open Targets locus-to-gene (L2G) rankings. For each disease, genes were ranked by L2G scores and each RBP regulon was tested for enrichment using GSEA. Points denote scRBP-prioritized regulons (x axis, $-\log_{10}$(P); y axis, normalized enrichment score (NES)).

(b) Example of a B cell–specific ZFP36L1 (3′UTR) regulon with genetic support across four additional autoimmune diseases (T1D, PBC, MS and CD). The regulator is shown as a diamond and target genes as circles; node color indicates Open Targets L2G score

for the corresponding disease.
(c) Example of a cycling-specific RBM17 (3′UTR) regulon associated with IBD. Right, regulon network (n = 40 target genes) with node colors denoting L2G scores and the corresponding GSEA significance indicated. Left, term–term similarity network of pathways enriched among RBM17 (3′UTR) targets (Metascape); nodes represent enriched terms, edges connect terms with high overlap, and node color encodes $-\log_{10}(P)$.
(d) Example of a monocyte-specific MAP1LC3B (5′UTR) regulon associated with RA. Right, regulon network (n = 109 target genes) with node colors denoting L2G scores and the corresponding GSEA significance indicated. Left, Metascape term–term similarity network for pathways enriched among MAP1LC3B (5′UTR) targets (color, $-\log_{10}(P)$).
(e) Example of a pDC-specific TOP1 (5′UTR) regulon associated with RA. Right, regulon network (n = 93 target genes) with node colors denoting L2G scores and the corresponding GSEA significance indicated. Left, Metascape term–term similarity network for pathways enriched among TOP1 (5′UTR) targets (color, $-\log_{10}(P)$).
(f) Example of a Tregs-specific WTAP (CDS) regulon associated with RA. Right, regulon network (n = 31 target genes) with node colors denoting L2G scores and the corresponding GSEA significance indicated. Left, Metascape term–term similarity network for pathways enriched among WTAP (CDS) targets (color, $-\log_{10}(P)$).

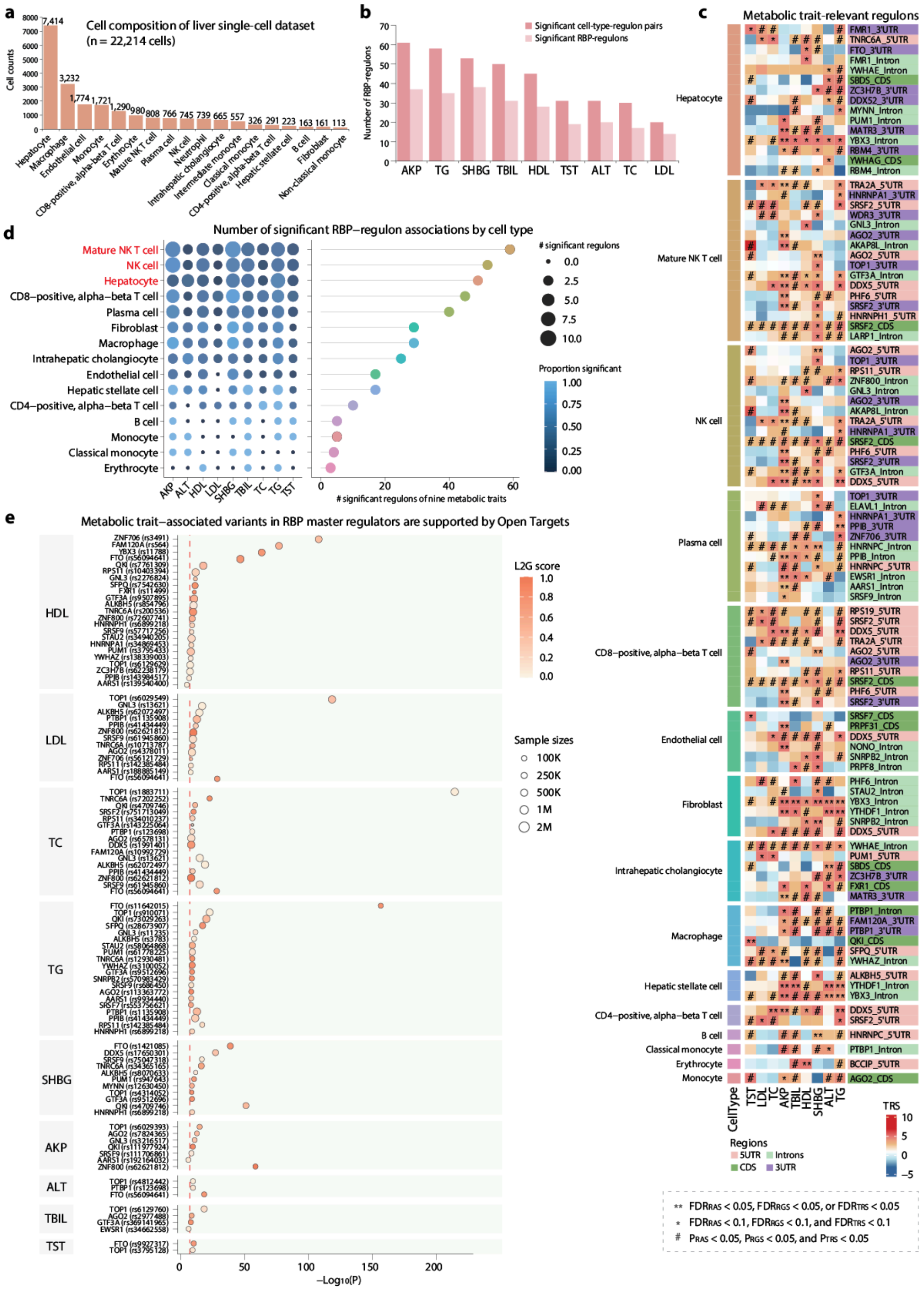


### Supplementary Fig. 18. scRBP identifies liver cell type–specific RBP regulons associated with nine metabolic traits.

(a) Cellular composition of the liver single-cell dataset used for scRBP analysis. Bar plot shows the number of cells for each annotated cell type, including hepatocytes, immune cell populations (e.g., NK cells, mature NK T cells), and other liver-associated cell types.

(b) Summary of significant RBP regulons associated with nine metabolic traits, including alkaline phosphatase (AKP), total cholesterol (TC), sex hormone–binding globulin (SHBG), total bilirubin (TBIL), high-density lipoprotein (HDL), testosterone (TST), alanine aminotransferase (ALT), triglycerides (TG), and low-density lipoprotein (LDL). Bars indicate the number of significant cell type–regulon pairs identified by scRBP. Dark segments represent significant cell type–regulon associations, whereas lighter segments denote significant RBP regulons.

(c) Heatmap summarizing metabolic trait–associated RBP regulons across cell types. Each row corresponds to an RBP regulon defined by both regulator and transcript region (5′UTR, CDS, intronic, and 3′UTR), and each column corresponds to a metabolic trait. Color intensity represents the regulon trait relevance score (TRS). ** indicates $FDR_{RAS} < 0.05$, $FDR_{RGS} < 0.05$, $FDR_{TRS} < 0.05$; * indicates FDR < 0.1 for all three scores; # indicates nominal significance ($P_{RAS}$, $P_{RGS}$, and $P_{TRS} < 0.05$). Statistical significance was assessed by MC permutation testing (one-sided, upper tail; 1,000 iterations; multiple-testing corrected). Region labels denote associated transcript regions (5′UTR, CDS, intronic, and 3′UTR).

(d) Distribution of significant RBP-regulon associations across cell types. Left: Bubble plot summarizing the number of significant regulons detected for each metabolic trait in each cell type. Bubble size reflects the number of significant RBP-regulon pairs, and color intensity represents the proportion of significant regulons within that cell type. Right: Lollipop plot ranking cell types by the total number of significant metabolic trait–associated regulons across all traits.

(e) Overlap between scRBP-identified regulons and genome-wide significant GWAS loci for nine metabolic traits. Points represent RBPs harboring genome-wide significant variants across metabolic traits. The x-axis shows $-\log_{10}(P)$ values derived from GWAS statistics; point color represents L2G scores from Open Targets genetics, and point size reflects GWAS sample size for each trait. Disease labels are indicated on the left.

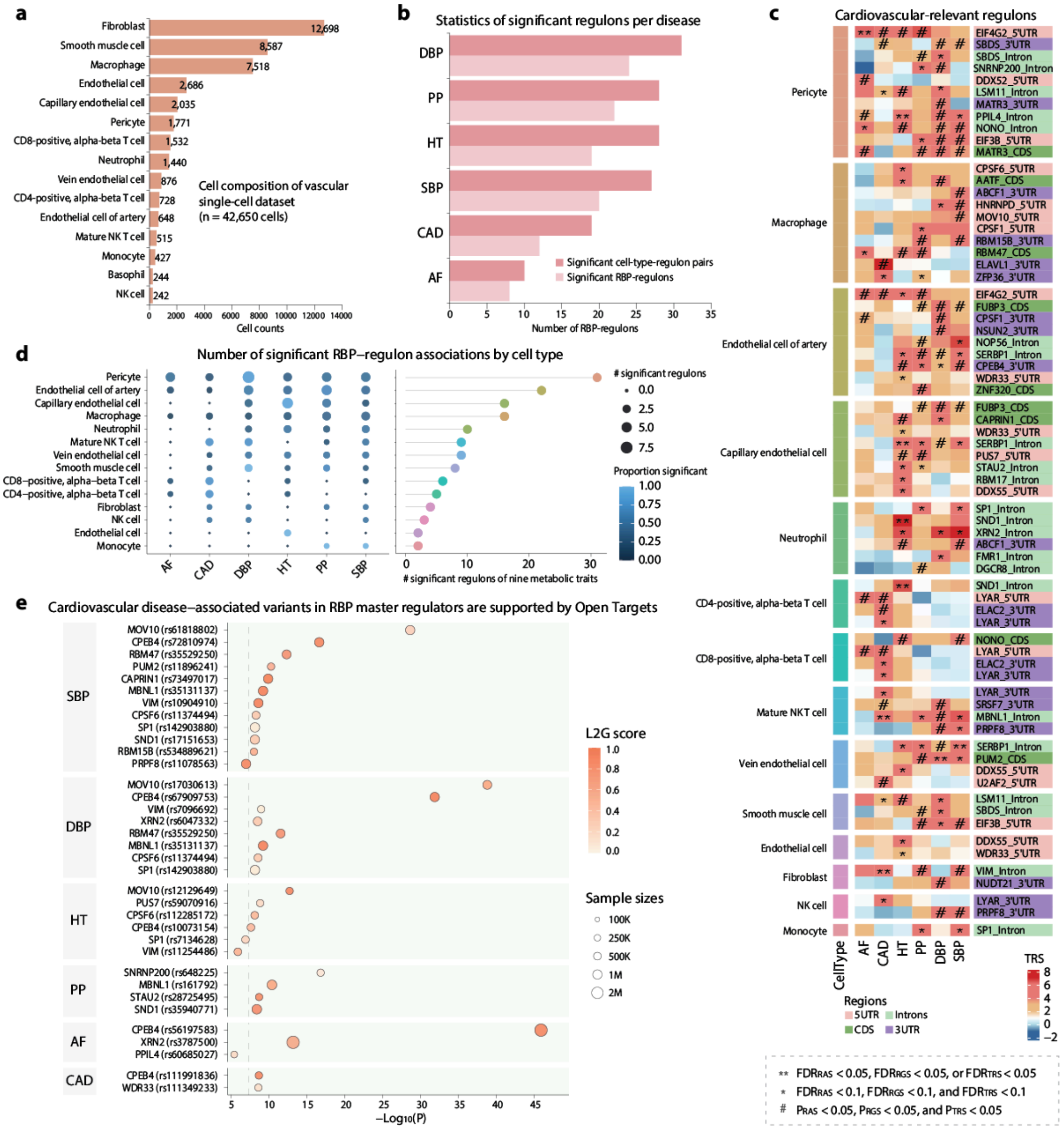


**Supplementary Fig. 19. scRBP identifies vascular cell type–specific RBP regulons associated with six cardiovascular diseases.**

(a) Cellular composition of the vascular single-cell dataset used for scRBP analysis. The bar plot shows the number of cells for each annotated vascular and immune cell type, including fibroblasts, smooth muscle cells, macrophages, endothelial subtypes, pericytes, and immune populations.

(b) Summary of significant RBP regulons associated with six cardiovascular traits and diseases, including diastolic blood pressure (DBP), pulse pressure (PP), hypertension (HT), systolic blood pressure (SBP), coronary artery disease (CAD), and atrial fibrillation (AF). Bars indicate the number of significant cell type–regulon pairs identified by scRBP. Dark segments represent significant cell type–regulon associations, while lighter segments denote significant RBP regulons.

(c) Heatmap summarizing cardiovascular disease–associated RBP regulons across vascular and immune cell types. Each row corresponds to an RBP regulon defined by both regulator and transcript region (5′UTR, CDS, intronic, and 3′UTR), and each column corresponds to a cardiovascular trait. Color intensity represents the regulon trait relevance score (TRS). ** indicates $FDR_{RAS} < 0.05$, $FDR_{RGS} < 0.05$, $FDR_{TRS} < 0.05$; * indicates FDR < 0.1 for all three scores; # indicates nominal significance ($P_{RAS}$, $P_{RGS}$, and $P_{TRS} < 0.05$). Statistical significance was assessed by MC permutation testing (one-sided, upper tail; 1,000 iterations; multiple-testing corrected). Region labels denote associated transcript regions (5′UTR, CDS, intronic, and 3′UTR).

(d) Distribution of significant RBP-regulon associations across vascular cell types. Left: Bubble plot summarizes the number of significant regulons detected for each cardiovascular trait in each cell type. Bubble size reflects the number of significant RBP-regulon pairs, and color intensity represents the proportion of significant regulons within that cell type. Right: Cell type ranking based on the number of significant cardiovascular trait–associated regulons. The Lollipop plot shows the total number of significant RBP regulons detected per cell type across all traits, highlighting vascular cell types with strong regulatory contributions.

(e) Overlap between scRBP-identified regulons and genome-wide significant GWAS loci. Points represent RBPs harboring genome-wide significant variants across six cardiovascular traits. The x-axis shows $-\log_{10}(P)$ values for variant associations; color indicates L2G scores and point size reflects GWAS sample size.

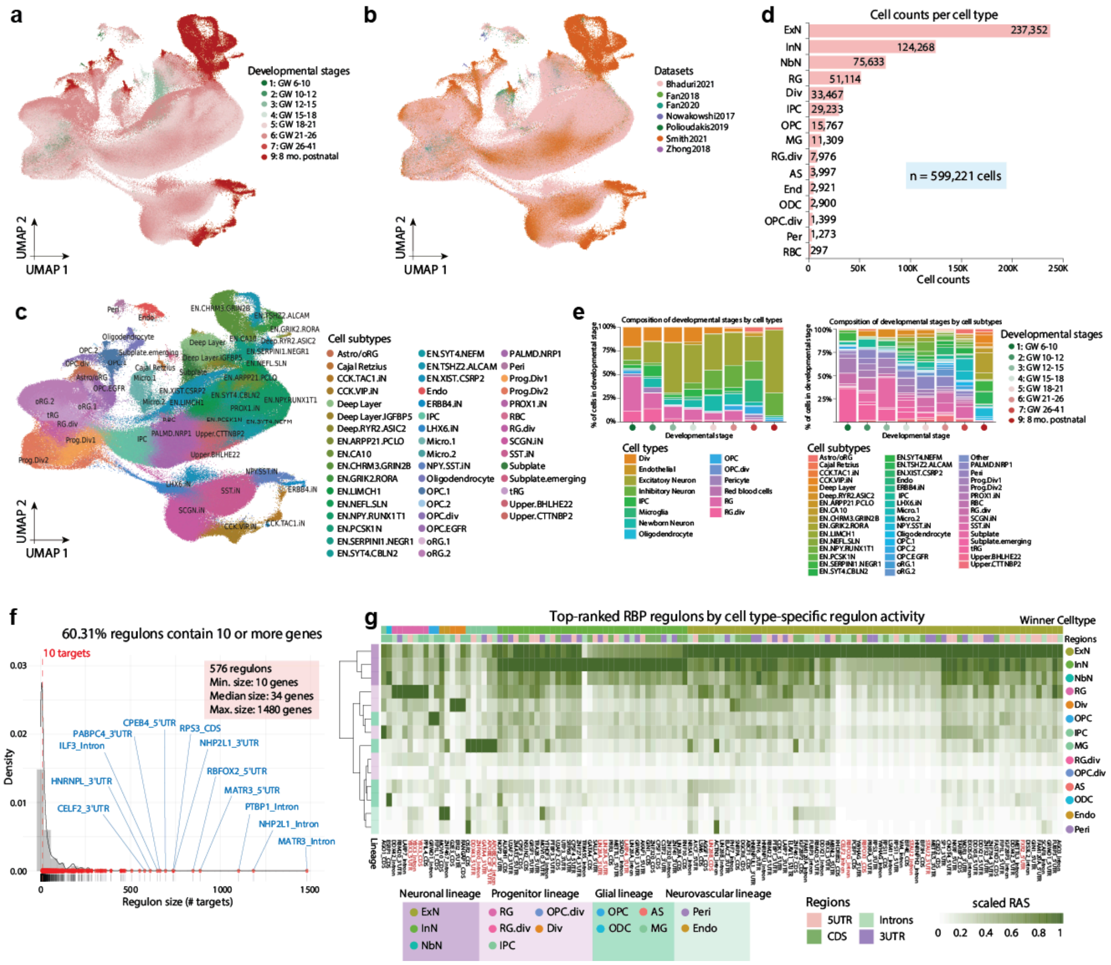


**Supplementary Fig. 20. Cell type-specific architecture of RBP regulons across the developing human cortex.**

(a) UMAP visualization of the integrated single-cell developmental brain atlas (n = 599,221 cells) colored by developmental stages.

Eight developmental stages within these individuals were delineated based on biological landmarks (Kang et al, *Nature* 2011).
(b) UMAP visualization of the integrated single-cell developmental brain atlas, colored by dataset sources. Seven independent single-cell datasets were included as shown in the plot.
(c) UMAP visualization of 47 neuronal and progenitor subtypes used for subtype-resolved regulon analysis.
(d) Distribution of cells across major developing human brain cell types. The integrated atlas comprises 15 annotated cell populations, including neuronal, progenitor, glial, vascular, and red blood cell (RBC) populations. As RBCs were rare (n = 297 cells), we excluded this population and retained 14 cell types for downstream analyses.
(e) Cell type composition across developmental stages. Stacked bar plots show the proportional distribution of major cell types (left) and refined cell subtypes (right) across developmental stages in the integrated human cortical atlas. Values represent the percentage of cells per stage.
(f) Size distribution of inferred RBP-regulons (number of target genes). The dashed vertical line indicates the minimum cutoff (≥10 target genes) used to define regulons for downstream analyses.
(g) Heatmap of top-ranked RBP regulons across major cell types based on regulon activity scores (RAS). Columns represent regulons and rows represent cell types grouped by lineage. Color indicates scaled RAS. Winner cell type annotations indicate the cell type with maximal regulon activity, and region labels denote associated transcript regions (5′UTR, CDS, intronic, and 3′UTR).

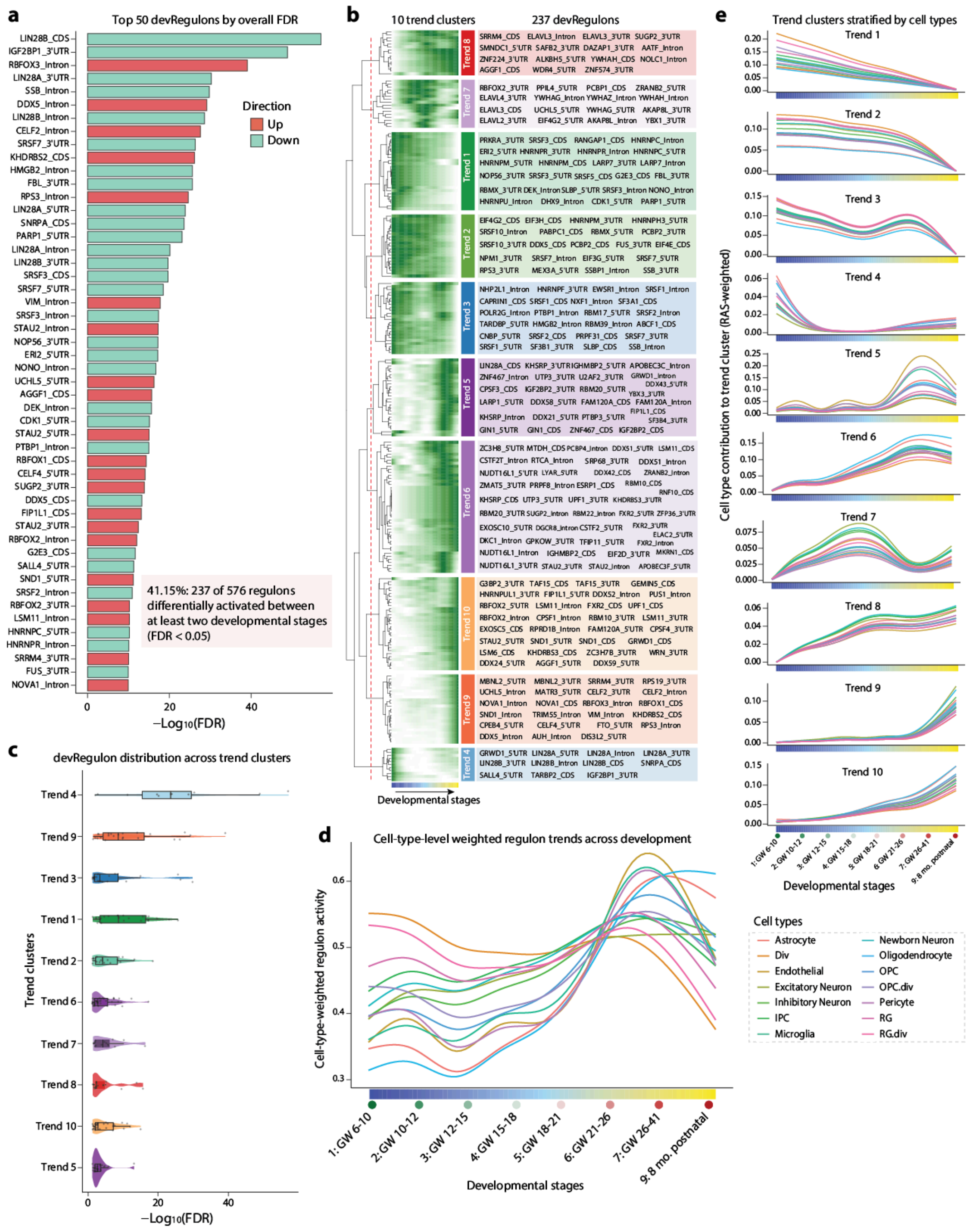

**Supplementary Fig. 21. Summary of devRegulons across 10 trend clusters.**

(a) Bar plot shows the top 50 devRegulons with significant differential activity across developmental stages, ranked by $-\log_{10}$(FDR). Colors indicate direction of activity change (upregulated or downregulated). Overall, 237 of 576 regulons (41.15%) exhibited significant stage-dependent activity (FDR < 0.05).

(b) Developmental trajectory clustering of RBP regulons. Heatmap shows activity patterns of 237 RBP regulons with significant developmental variation (FDR < 0.05) across developmental stages. Regulons were grouped into ten temporal trend clusters based on similarity of activity trajectories. DevRegulons for each trend cluster are indicated.

(c) Distribution of developmentally dynamic RBP-regulons across trend clusters. Violin plots show $-\log_{10}$(FDR) distributions of devRegulons within each temporal trend cluster.

(d) Cell type–level trajectories of RBP regulon activity across cortical development. Curves represent RAS-weighted average developmental trajectories of RBP regulons for each major cell type. Regulon trend profiles were derived from the global developmental trend matrix and weighted by cell type–specific RAS, normalized within each cell type to control for overall activity differences. The x-axis indicates developmental stages spanning gestational weeks to postnatal periods.

(e) Trend cluster–resolved cell type trajectories of RBP regulon activity. Curves show RAS-weighted developmental trajectories of cell type–specific regulon activity within each regulon trend cluster.

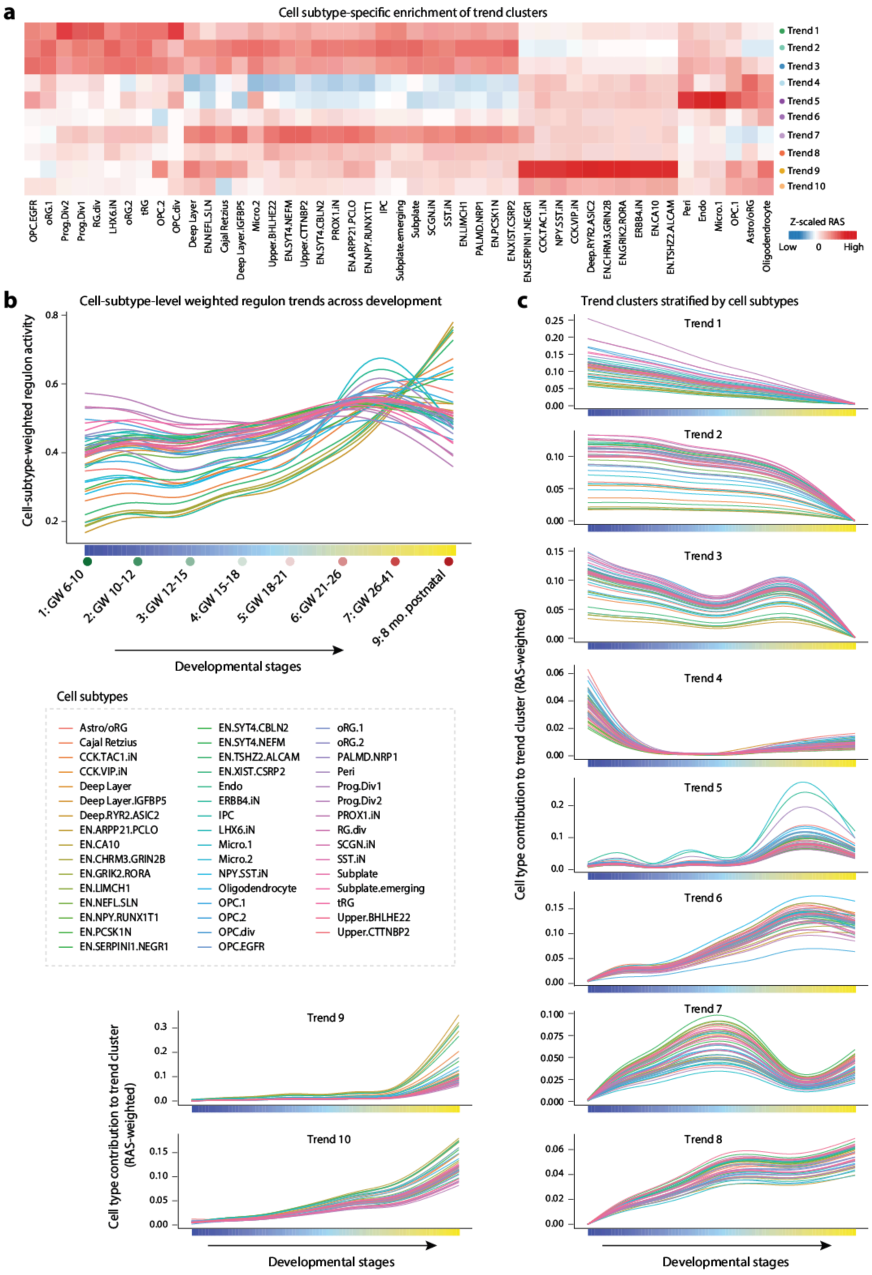


**Supplementary Fig. 22. Cell subtype-resolved analysis of developmental RBP regulon dynamics.**

(a) Heatmap showing cell subtype–resolved enrichment of developmental regulon trend clusters. For each cell subtype, enrichment was calculated by averaging regulon activity scores (RAS) across regulons belonging to the same trend cluster, followed by z-score normalization across clusters within each subtype. Columns represent 47 neuronal and progenitor subtypes, and rows represent 10 developmental regulon trends.

(b) Cell subtype–level trajectories of RBP regulon activity across development. Curves show RAS-weighted developmental trajectories of RBP regulons for each cell subtype. Trend profiles were derived from the global developmental trend matrix and weighted by subtype-specific RAS, normalized within each subtype. The x-axis denotes developmental stages from gestational weeks to postnatal periods.

(c) Trend cluster–resolved cell subtype trajectories of RBP regulon activity. Curves show RAS-weighted developmental trajectories of cell subtype–specific regulon activity within each regulon trend cluster.

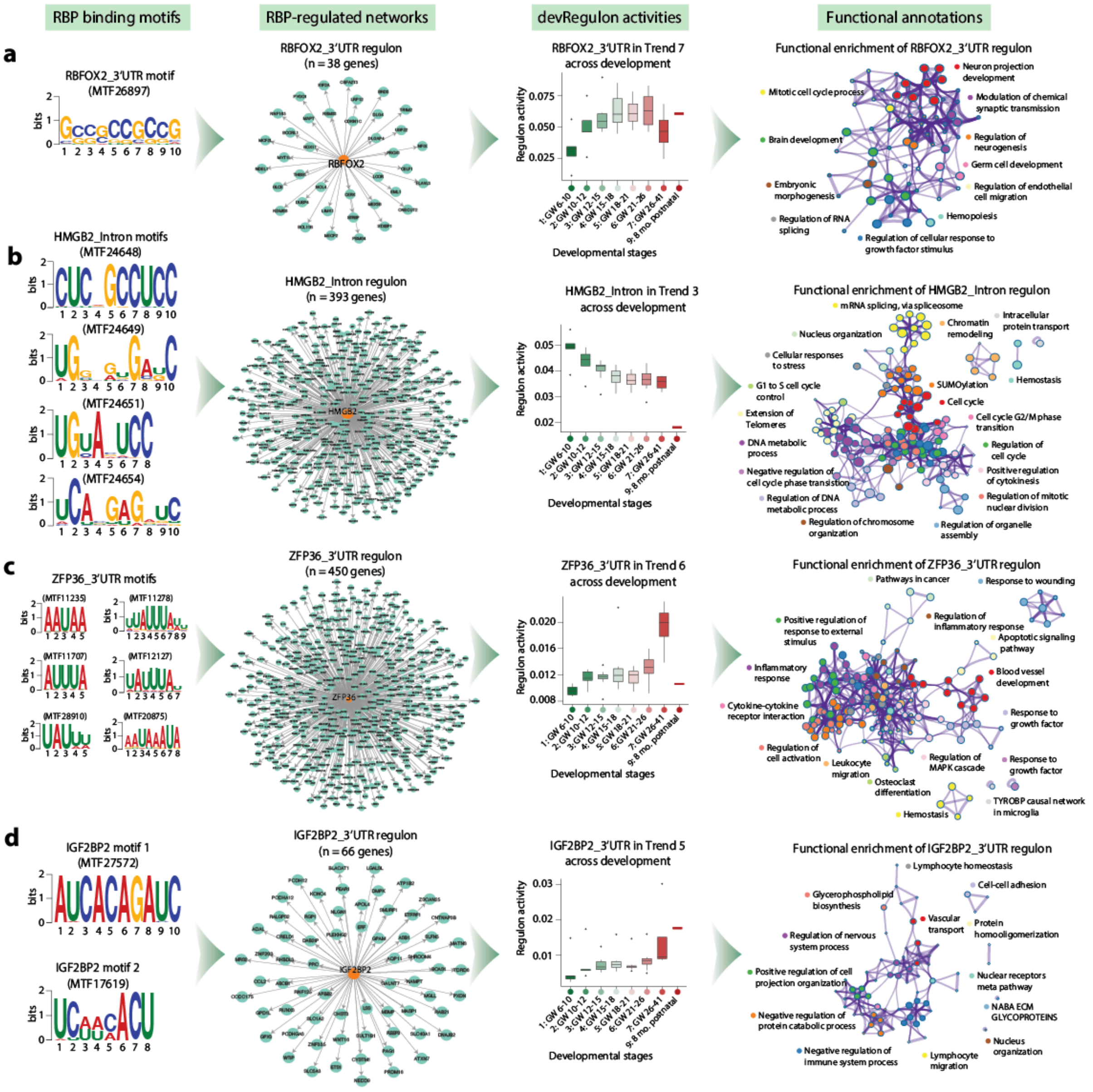


**Supplementary Fig. 23. Four representative devRegulons across cortical development.**

(a) RBFOX2 (3′UTR) regulon motif, target network, and developmental activity. Sequence logo of the canonical RBFOX2 RNA-binding motif identified in 3′UTR regions is shown together with a network representation of the corresponding RBFOX2 3′UTR regulon, comprising predicted target transcripts inferred from motif-based ranking and co-expression refinement. Boxplots depict the developmental trajectory of regulon activity across fetal-to-postnatal stages, illustrating stage-dependent variation in RBFOX2-associated post-transcriptional regulation (Trend 7). Functional enrichment analysis of the RBFOX2 (3′UTR) (n = 38 genes).

(b) HMGB2 (intronic) regulon motif, network, and developmental activity. Representative HMGB2 intronic binding motifs, corresponding regulon network, and developmental activity profile across cortical stages (Trend 3). Functional enrichment analysis of the HMGB2 (intronic) (n = 393 genes).

(c) ZFP36 (3′UTR) regulon motif, network, and developmental activity. ZFP36 3′UTR binding motifs, regulon target network, and developmental dynamics of regulon activity across stages (Trend 6). Functional enrichment analysis of the ZFP36 (3′UTR) (n = 450 genes).

(d) IGF2BP2 (3′UTR) regulon motif, network, and developmental activity. Representative IGF2BP2 3′UTR binding motifs, inferred regulon network, and developmental activity trajectory (Trend 5). Functional enrichment analysis of the IGF2BP2 (3′UTR) (n = 66 genes). The pathway enrichment analyses were performed by using Metascape (https://metascape.org/).

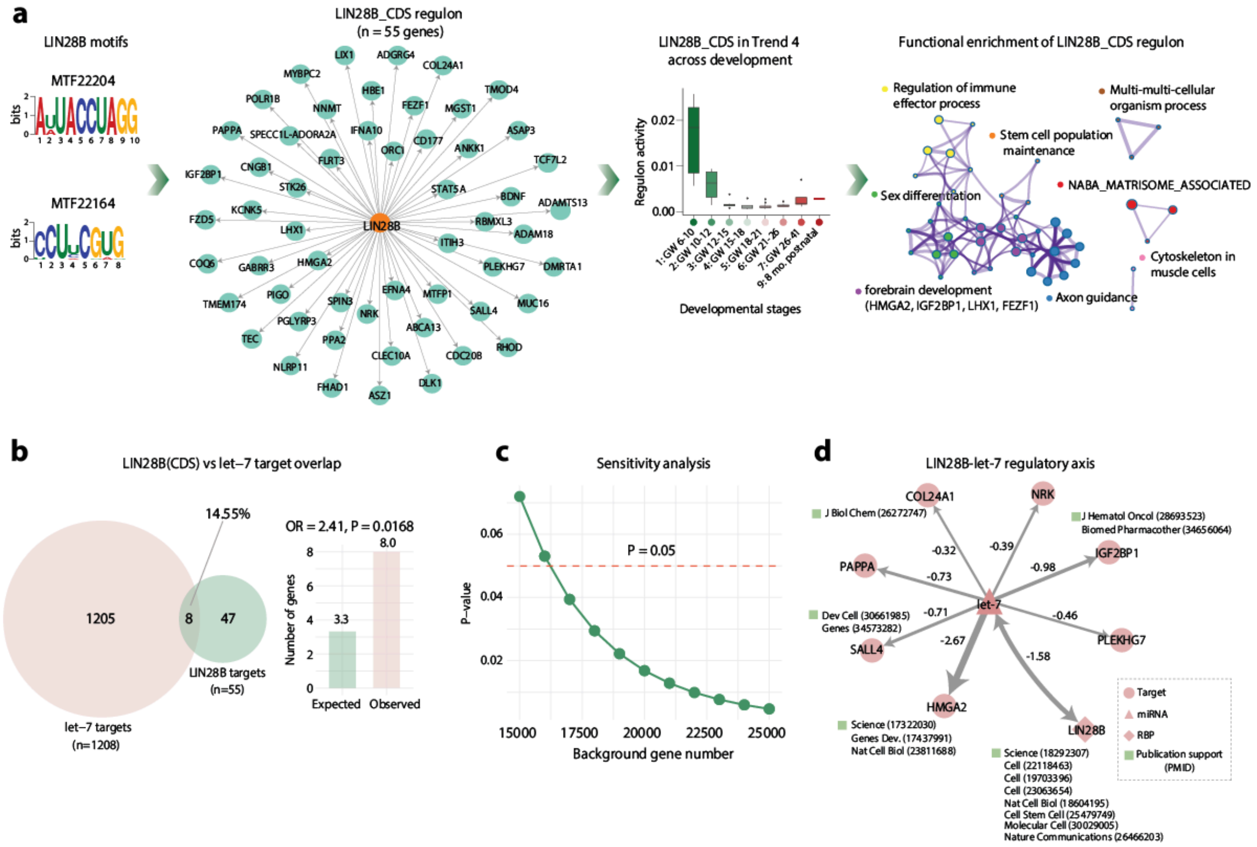


### Supplementary Fig. 24. LIN28B (CDS) regulon motif, network, developmental activity, and let-7 regulatory axis.

(a) Representative LIN28B (CDS) binding motifs, inferred regulon network, and developmental activity trajectory (Trend 5). Functional enrichment analysis of the LIN28B (CDS) (n = 55 genes). The pathway enrichment analysis was performed by using Metascape (https://metascape.org/).

(b) Overlap between predicted LIN28B (CDS) regulon targets and let-7 miRNA target genes. Left panel: The Venn diagram shows shared and unique targets between the LIN28B (CDS) regulon (n = 55) and let-7 targets (n = 1,208). Eight genes (14.55% of LIN28B targets) are common to both sets, suggesting potential interaction between LIN28B regulation and the let-7 regulatory axis. Right panel: The bar plot compares the expected versus observed number of let-7 target genes among LIN28B CDS regulon targets. A significant enrichment was observed (OR = 2.41, P = 0.0168), supporting a potential regulatory relationship between LIN28B and the let-7 pathway.

(c) Sensitivity analysis of let-7 target enrichment in the LIN28B (CDS) regulon. The plot shows enrichment P-values calculated using varying background gene set sizes. The dashed horizontal line indicates the significance threshold (P = 0.05). Enrichment remained significant across a range of background sizes, supporting the robustness of the observed LIN28B–let-7 regulatory association.

(d) Regulatory network illustrating the LIN28B–let-7 axis. Nodes represent predicted let-7 target genes within the LIN28B (CDS) regulon together with related regulatory components. Edges indicate TargetScan-predicted miRNA–target interactions, ranked by cumulative weighted context++ scores (more negative values indicate stronger predicted repression). Supporting literature evidence (PMIDs) is annotated where available. This network highlights the post-transcriptional regulatory relationships underlying the LIN28B–let-7 regulatory axis.

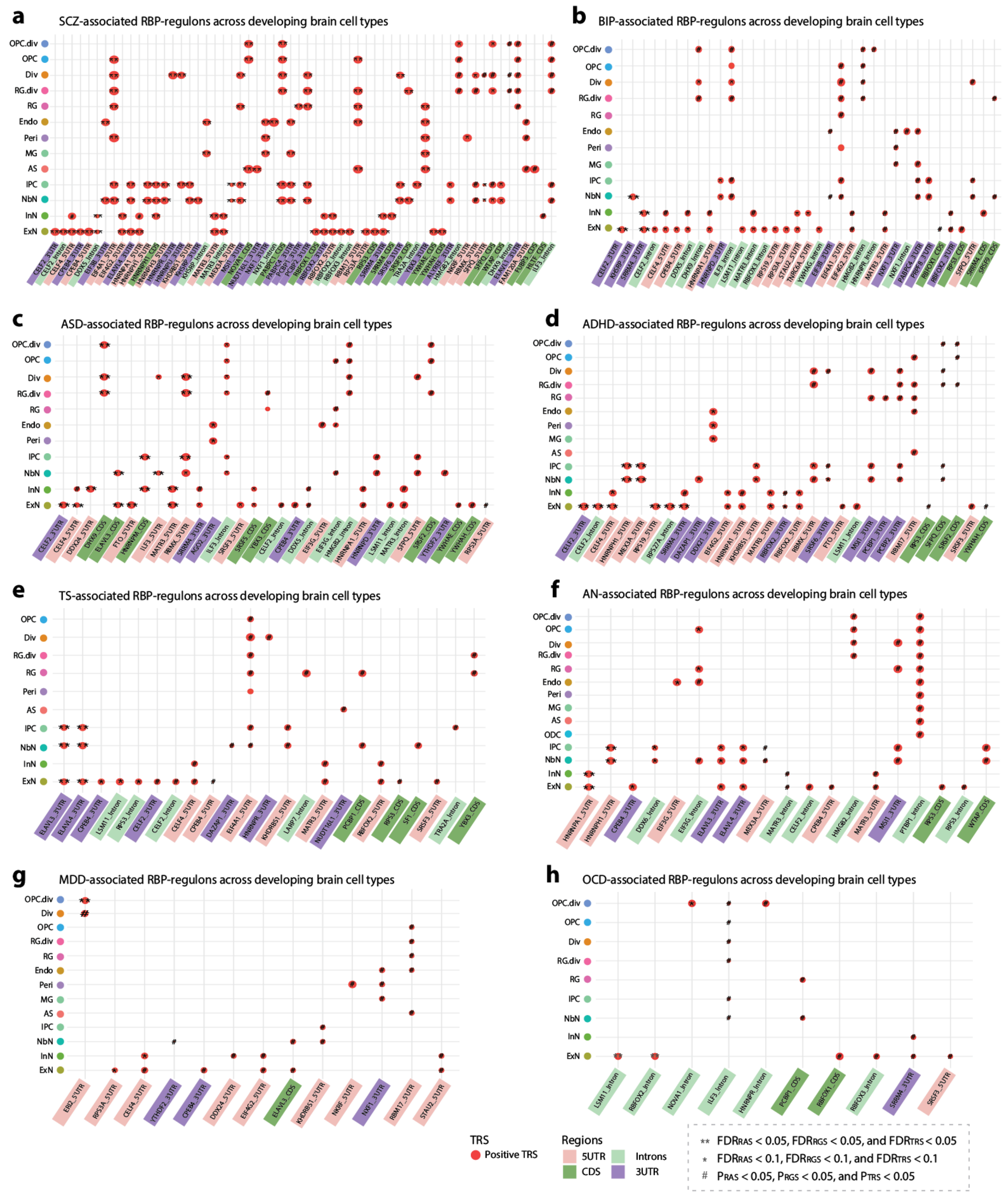


**Supplementary Fig. 25. scRBP identifies brain cell type-specific RBP regulons associated with eight neuropsychiatric disorders.**

Dot plots show the distribution of neuropsychiatric disorder-associated RBP regulons across major fetal brain cell types. Each column represents an RBP regulon (annotated by genomic binding region), and each row corresponds to a cell type. Red dots indicate the significant positive trait-relevant scores (TRS) for each disorder. ** indicates $FDR_{RAS} < 0.05$, $FDR_{RGS} < 0.05$, $FDR_{TRS} < 0.05$; * indicates FDR < 0.1 for all three scores; # indicates nominal significance ($P_{RAS}$, $P_{RGS}$, and $P_{TRS} < 0.05$). Statistical significance was assessed by Monte Carlo permutation testing (one-sided upper tail test; 1,000 permutations; multiple-testing corrected). Panels show results for (a) SCZ, (b) BIP, (c) ASD, (d) ADHD, (e) TS, (f) AN, (g) MDD, and (h) OCD.

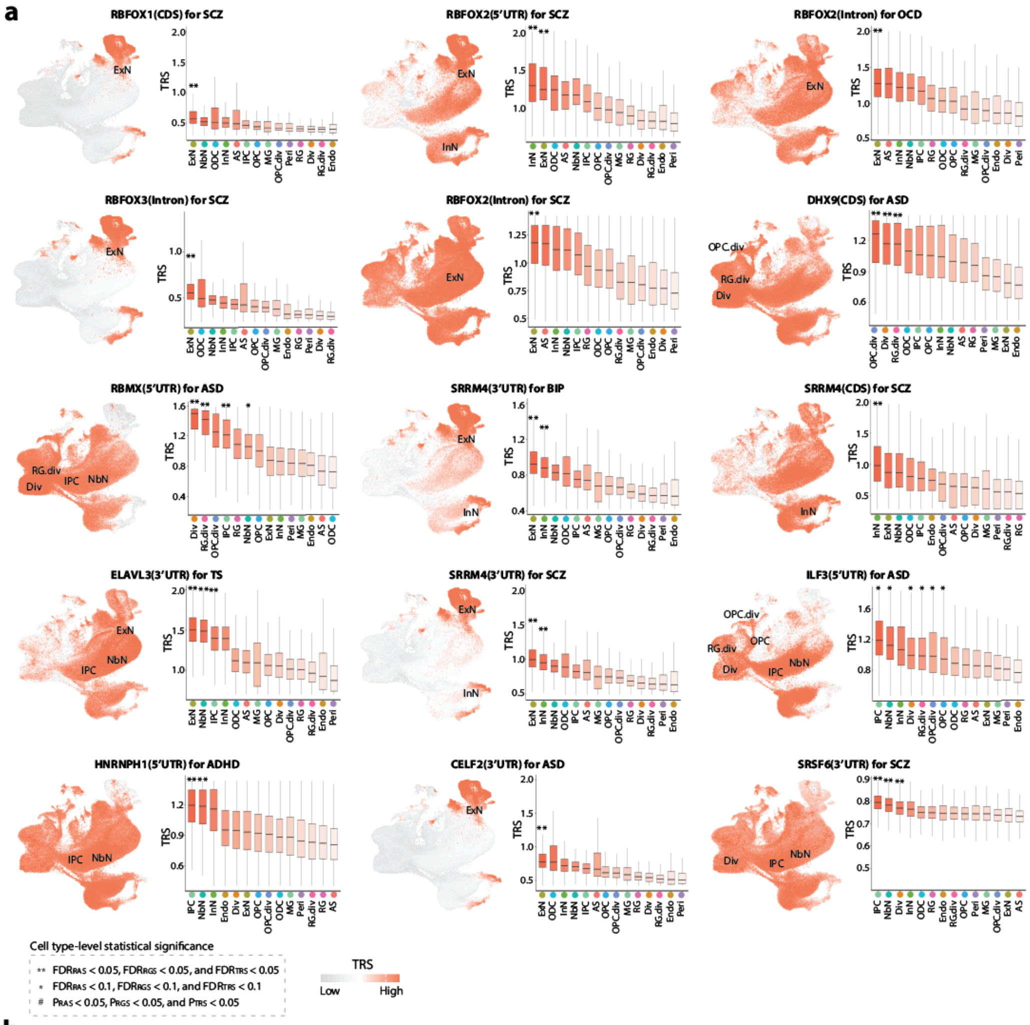


**b**

| Traits | Regulons | Cell types | RBPs | Regions | RAS | RGS | TRS | FDR_RAS | FDR_RGS | FDR_TRS | N_Null |
|---|---|---|---|---|---|---|---|---|---|---|---|
| AD | RPS11_5'UTR | ExN | RPS11 | 5'UTR | 2.07 | 7.37 | 4.16 | 0.065 | 0.142 | 0.071 | 1000 |
| AD | LSM11_Intron | ExN | LSM11 | Intron | 8.01 | 4.07 | 4.96 | 0.014 | 0.278 | 0.069 | 1000 |
| AD | HNRNPR_3'UTR | Div | HNRNPR | 3'UTR | 3.14 | 5.12 | 5.88 | 0.031 | 0.357 | 0.076 | 1000 |
| PD | SRSF2_CDS | Div | SRSF2 | CDS | 4.54 | 5.91 | 4.96 | 0.024 | 0.142 | 0.047 | 1000 |
| PD | EIF3B_3'UTR | NbN | EIF3B | 3'UTR | 4.44 | 7.65 | 4.85 | 0.010 | 0.153 | 0.102 | 1000 |
| PD | EIF3B_3'UTR | IPC | EIF3B | 3'UTR | 2.17 | 7.65 | 4.80 | 0.072 | 0.153 | 0.102 | 1000 |
| PD | CELF4_5'UTR | ExN | CELF4 | 5'UTR | 7.64 | 5.08 | 4.26 | 0.014 | 0.213 | 0.071 | 1000 |
| PD | SRSF9_5'UTR | NbN | SRSF9 | 5'UTR | 2.92 | 5.62 | 4.81 | 0.035 | 0.213 | 0.071 | 1000 |
| PD | AGO1_CDS | IPC | AGO1 | CDS | 5.00 | 3.38 | 5.24 | 0.016 | 0.213 | 0.071 | 1000 |
| PD | XRN2_CDS | NbN | XRN2 | CDS | 3.25 | 2.81 | 2.78 | 0.020 | 0.213 | 0.071 | 1000 |

**Supplementary Fig. 26. Single-cell-resolved scRBP analysis reveals neuropsychiatric disorder–associated regulons enriched in high-TRS fetal brain cells.**

(a) Representative disorder-associated RBP regulons shown at single-cell resolution. Left, UMAP feature plots displaying per-cell trait-relevant scores (TRSs), highlighting enrichment of regulon activity within specific fetal brain populations. Right, distributions of per-cell TRSs across major cell types for the corresponding regulons (boxplots; center line, median; box, interquartile range; whiskers, 1.5× IQR).

(b) Negative-control analysis for Alzheimer's disease (AD) and Parkinson's disease (PD). The table summarizes top-scoring regulon–cell-type associations and their corresponding scRBP metrics (RAS, RGS and TRS) with Monte Carlo permutation–based significance estimates (1,000 permutations). No regulons remained significant for AD or PD after multiple-testing correction across all three metrics at the indicated thresholds ($FDR_{RAS}$, $FDR_{RGS}$, and $FDR_{TRS}$< 0.05 or 0.1).

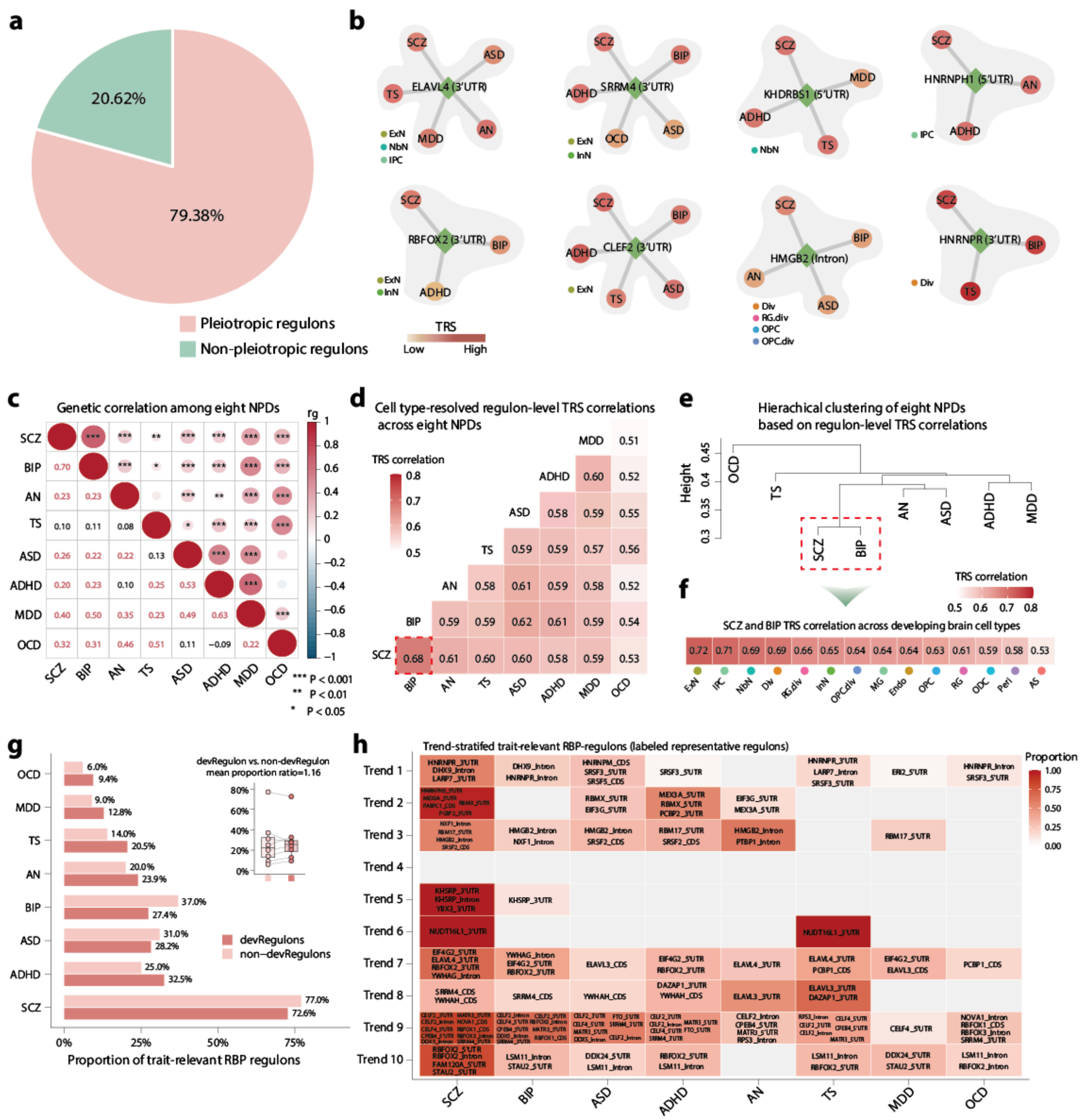


**Supplementary Fig. 27. Cross-disorder pleiotropy and developmental enrichment of psychiatric trait-relevant RBP regulons.**

(a) Distribution of psychiatric disorder–associated RBP regulons according to the number of disorders with significant trait-relevant scores (TRSs). Most trait-relevant regulons (79.38%) were associated with two or more disorders, indicating pervasive regulatory pleiotropy.

(b) Examples of highly pleiotropic RBP regulons shared across multiple neuropsychiatric disorders. Nodes represent disorders and edges denote significant regulon–trait associations; node color encodes trait-relevant scores (TRSs). Selected regulons, including ELAVL4 (3′UTR), SRRM4 (3′UTR), and CELF2 (3′UTR), were associated with three to five disorders.

(c) Genetic correlation among eight neuropsychiatric disorders (NPDs), calculated using cross-trait LD score regression (LDSC). Circle size and color intensity indicate the magnitude of genetic correlation. Asterisks denote statistically significant genetic correlations (* $P < 0.05$, ** $P < 0.01$, *** $P < 0.001$).

(d) Heatmap showing correlations of cell-type-resolved regulon TRSs across eight NPDs. The boxed region highlights strong concordance of trait relevance between SCZ and BIP.

(e) Hierarchical clustering of eight NPDs based on regulon-level TRS correlations, showing close clustering of SCZ and BIP and a broader neurodevelopmental cluster comprising ADHD, ASD, and MDD.

(f) Cell type–resolved TRS correlations between SCZ and BIP across developing brain cell types, with strongest concordance in neuronal and progenitor populations, particularly ExN.

(g) Proportion of trait-relevant RBP regulons among development-associated regulons (devRegulons) and non-development-associated regulons across eight NPDs (bar plots). Inset boxplot shows paired comparisons of trait-relevant regulon proportions between devRegulons and non-devRegulons across disorders, with connecting lines indicating matched disorder-level comparisons.

(h) Trend-stratified enrichment of trait-relevant RBP regulons across neuropsychiatric disorders. Rows denote developmental regulon trend clusters and columns indicate disorders. Color intensity reflects the proportion of trait-relevant regulons within each cluster. Representative top-ranked regulons are labeled.

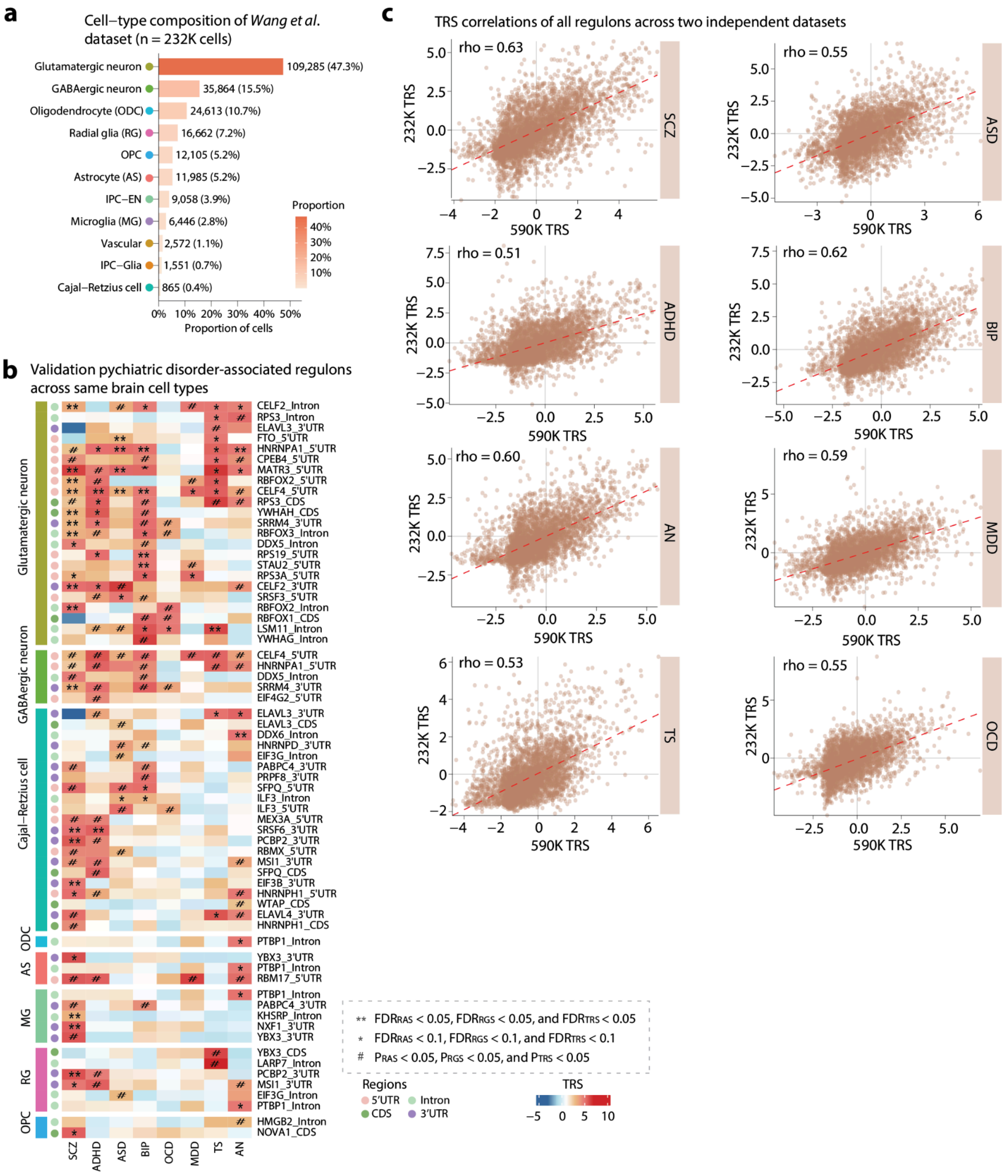


### Supplementary Fig. 28. Concordance of regulon-level trait-relevant scores between independent developmental brain single-cell datasets.

(a) Cell type composition of 11 major fetal brain cell types in the Wang et al. developing neocortex single-cell dataset (n = 232,328 cells).

(b) Heatmap showing cell type-resolved RBP-regulons associated with eight neuropsychiatric disorders (SCZ, BIP, ASD, ADHD, AN,

MDD, TS, and OCD) in the validation dataset. Statistical significance is reported for three complementary metrics (RAS, RGS and TRS). ** indicates $FDR_{RAS} < 0.05$, $FDR_{RGS} < 0.05$, $FDR_{TRS} < 0.05$; * indicates FDR < 0.1 for all three scores; # indicates nominal significance ($P_{RAS}$, $P_{RGS}$, and $P_{TRS} < 0.05$). P values were estimated by Monte Carlo permutation (one-sided, upper tail; 1,000 iterations) and adjusted for multiple testing. Region labels denote transcript regions (5′UTR, CDS, intronic, and 3′UTR).
(c) Cell type–resolved TRS correlations of all regulons between the discovery fetal brain dataset (n = 599,221 cells; 590K) and the independent developing neocortex validation dataset (n = 232,328 cells; 232K) across eight neuropsychiatric disorders. Pearson correlation coefficients are shown.

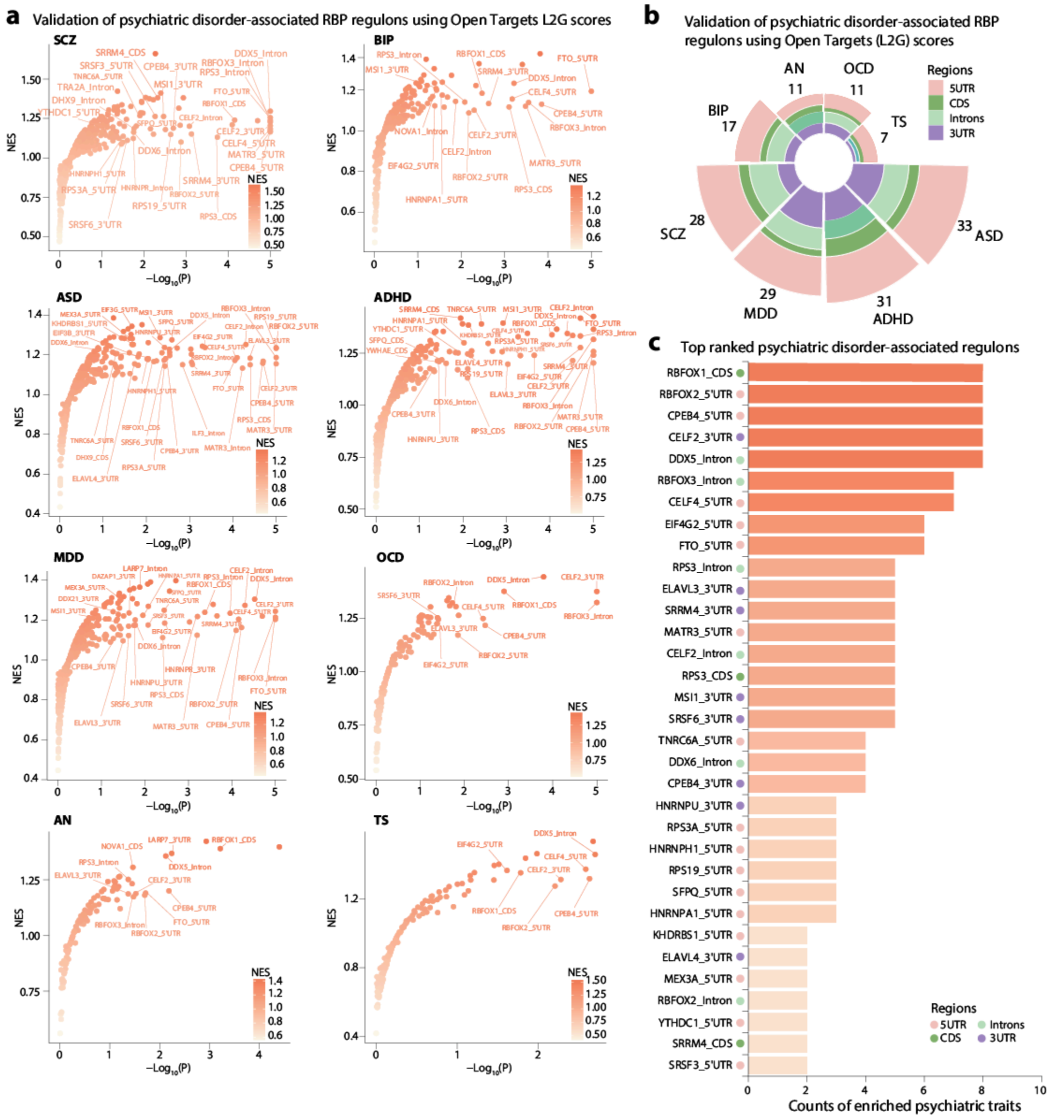


### Supplementary Fig. 29. Orthogonal validation of scRBP-prioritized regulons using neuropsychiatric disorder-specific genetic prioritization scores.

(a) Gene set enrichment analysis (GSEA) of scRBP-identified disorder-associated regulons using disorder-specific locus-to-gene (L2G) prioritization scores from the Open Targets platform. Genes were ranked by L2G scores for each disorder, and enrichment of regulon target genes was evaluated against these ranked lists.
(b) Radial stacked bar plot showing the number of neuropsychiatric disorder-associated RBP regulons across diseases, stratified by transcript regions (5’UTR, CDS, intronic, and 3’UTR).
(c) Examples of recurrently enriched regulons across multiple disorders. Selected top-ranked regulons, including RBFOX1 (CDS), RBFOX2 (5′UTR), CPEB4 (5′UTR), CELF2 (3′UTR) and RBFOX3 (intronic), exhibited enrichment across two or more disorders, consistent with shared post-transcriptional regulatory programs across neuropsychiatric traits.

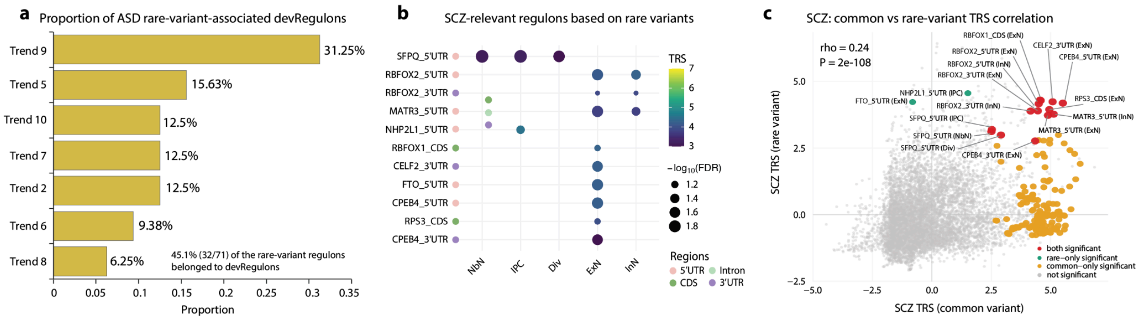


d Top 10 ranked regulons for each negative control trait (AD, PD, LDL-C) using scRBP rare-variant model

| Traits | Regulons | Cell types | RBPs | Regions | RAS | RGS | TRS | FDR_RAS | FDR_RGS | FDR_TRS | N_Null |
|---|---|---|---|---|---|---|---|---|---|---|---|
| AD | CAPRIN1_CDS | Inhibitory Neuron | CAPRIN1 | CDS | 1.92 | 4.31 | 4.65 | 0.317 | 0.278 | 0.139 | 1000 |
| AD | LIN28A_Intron | Inhibitory Neuron | LIN28A | Introns | 2.05 | 3.84 | 5.61 | 0.185 | 0.764 | 0.139 | 1000 |
| AD | LIN28A_Intron | RG | LIN28A | Introns | 1.74 | 3.84 | 4.90 | 0.195 | 0.764 | 0.139 | 1000 |
| AD | LIN28A_Intron | IPC | LIN28A | Introns | 1.61 | 3.84 | 4.70 | 0.296 | 0.764 | 0.139 | 1000 |
| AD | LIN28A_Intron | Newborn Neuron | LIN28A | Introns | 1.71 | 3.84 | 4.83 | 0.311 | 0.764 | 0.139 | 1000 |
| AD | SF3A3_Intron | Newborn Neuron | SF3A3 | Introns | 1.13 | 10.38 | 4.04 | 0.433 | 0.139 | 0.185 | 1000 |
| AD | ACIN1_CDS | OPC | ACIN1 | CDS | 0.77 | 5.00 | 3.27 | 0.583 | 0.278 | 0.278 | 1000 |
| AD | FIP1L1_CDS | OPC | FIP1L1 | CDS | 14.03 | 2.60 | 3.91 | 0.035 | 0.625 | 0.278 | 1000 |
| AD | FIP1L1_CDS | Oligodendrocyte | FIP1L1 | CDS | 10.87 | 2.60 | 4.51 | 0.035 | 0.625 | 0.278 | 1000 |
| AD | AGO2_CDS | Inhibitory Neuron | AGO2 | CDS | 1.50 | 3.81 | 4.52 | 0.528 | 0.347 | 0.278 | 1000 |
| PD | FASTKD2_CDS | Newborn Neuron | FASTKD2 | CDS | 3.03 | 4.11 | 4.70 | 0.027 | 0.284 | 0.142 | 1000 |
| PD | FASTKD2_CDS | Inhibitory Neuron | FASTKD2 | CDS | 2.19 | 4.11 | 4.92 | 0.248 | 0.284 | 0.142 | 1000 |
| PD | ZCCHC14_CDS | IPC | ZCCHC14 | CDS | 1.37 | 3.58 | 3.41 | 0.299 | 0.284 | 0.142 | 1000 |
| PD | FASTKD2_CDS | IPC | FASTKD2 | CDS | 0.82 | 4.11 | 2.77 | 0.296 | 0.284 | 0.199 | 1000 |
| PD | ALKBH3_CDS | Newborn Neuron | ALKBH3 | CDS | 2.35 | 3.14 | 4.43 | 0.057 | 0.378 | 0.142 | 1000 |
| PD | PCBP1_CDS | Newborn Neuron | PCBP1 | CDS | 1.78 | 2.89 | 3.84 | 0.119 | 0.378 | 0.142 | 1000 |
| PD | BCLAF1_5'UTR | Div | BCLAF1 | 5UTR | 2.97 | 4.46 | 4.91 | 0.071 | 0.567 | 0.071 | 1000 |
| PD | BCLAF1_5'UTR | IPC | BCLAF1 | 5UTR | 2.16 | 4.46 | 4.82 | 0.086 | 0.567 | 0.071 | 1000 |
| PD | BCLAF1_5'UTR | RG | BCLAF1 | 5UTR | 1.92 | 4.46 | 4.32 | 0.236 | 0.567 | 0.071 | 1000 |
| PD | BCLAF1_5'UTR | RG.div | BCLAF1 | 5UTR | 1.78 | 4.46 | 4.14 | 0.298 | 0.567 | 0.071 | 1000 |
| LDL-C | KHSRP_CDS | IPC | KHSRP | CDS | 4.11 | 4.00 | 4.73 | 0.016 | 0.227 | 0.071 | 1000 |
| LDL-C | KHSRP_CDS | Newborn Neuron | KHSRP | CDS | 3.52 | 4.00 | 4.62 | 0.020 | 0.227 | 0.071 | 1000 |
| LDL-C | FAM120A_CDS | Red blood cells | FAM120A | CDS | 2.88 | 3.79 | 5.75 | 0.340 | 0.227 | 0.071 | 1000 |
| LDL-C | SRSF5_CDS | RG | SRSF5 | CDS | 1.91 | 3.80 | 4.07 | 0.335 | 0.227 | 0.071 | 1000 |
| LDL-C | SRSF3_CDS | IPC | SRSF3 | CDS | 1.55 | 3.86 | 3.87 | 0.232 | 0.227 | 0.071 | 1000 |
| LDL-C | SRSF3_CDS | Newborn Neuron | SRSF3 | CDS | 1.64 | 3.86 | 3.96 | 0.138 | 0.227 | 0.071 | 1000 |
| LDL-C | FMR1_CDS | Excitatory Neuron | FMR1 | CDS | 2.20 | 3.48 | 4.45 | 0.231 | 0.227 | 0.095 | 1000 |
| LDL-C | KHSRP_CDS | Red blood cells | KHSRP | CDS | 2.82 | 4.00 | 5.59 | 0.355 | 0.227 | 0.095 | 1000 |
| LDL-C | SRSF5_CDS | OPC | SRSF5 | CDS | 2.43 | 3.80 | 4.51 | 0.122 | 0.227 | 0.142 | 1000 |
| LDL-C | SRSF3_CDS | Div | SRSF3 | CDS | 2.69 | 3.86 | 4.66 | 0.142 | 0.227 | 0.142 | 1000 |

**Supplementary Fig. 30. Developmental patterning and specificity of rare-variant-associated RBP regulons.**

(a) Distribution of ASD rare-variant-associated developmentally dynamic regulons across temporal trajectories. Of the 71 regulons associated with ASD rare-variant burden, 32 (45.1%) were classified as devRegulons. Bars show the proportion of these 32 regulons assigned to each developmental trajectory; only trajectories containing at least one ASD-associated devRegulon are shown.

(b) Cell type-resolved RBP regulons associated with SCZ rare-variant burden. The 16 significant regulon–cell type associations comprised 11 regulons from 9 RBPs across 5 cell types. Rows denote regulons defined by an RBP and its targeted transcript region, and columns denote cell types. Dots indicate associations significant across all three scRBP components ($FDR_{RAS}$, $FDR_{RGS}$ and $FDR_{TRS}$ < 0.1). Dot colour denotes the trait-relevance score (TRS), and dot size denotes $-\log_{10}$ of the maximum FDR across the three tests. Row annotations indicate the targeted transcript region.

(c) Relationship between common- and rare-variant TRSs across all SCZ regulon–cell type pairs tested (n=8,064). Points are classified as significant in both models (n = 14), in the rare-variant model only (n = 2), in the common-variant model only (n = 133) or in neither model. Spearman's correlation is shown (ρ = 0.24, two-sided $P = 2 \times 10^{-108}$). Regulons significant in both models are labelled.

(d) Top-ranked regulon–cell type associations for three comparison traits: Alzheimer's disease (AD), Parkinson's disease (PD) and LDL cholesterol (LDL-C). The ten highest-TRS associations for each trait are shown together with their regulon activity score (RAS), regulon genetic score (RGS), integrated TRS and FDR values. Scores were calibrated using 1,000 matched null regulons. No association met the joint significance criterion applied throughout the rare-variant analysis ($FDR_{RAS}$, $FDR_{RGS}$ and $FDR_{TRS}$ < 0.1).

Note: FDR values were adjusted using the Benjamini–Hochberg method. NbN, newborn neuron; IPC, intermediate progenitor cell; ExN, excitatory neuron; Div, dividing cell; RG.div, dividing radial glia; Endo, endothelial cell; OPC, oligodendrocyte precursor cell; RG, radial glia; Peri, pericyte; OPC.div, dividing oligodendrocyte precursor cell; InN, inhibitory neuron; MG, microglia; ODC, oligodendrocyte; AS, astrocyte.

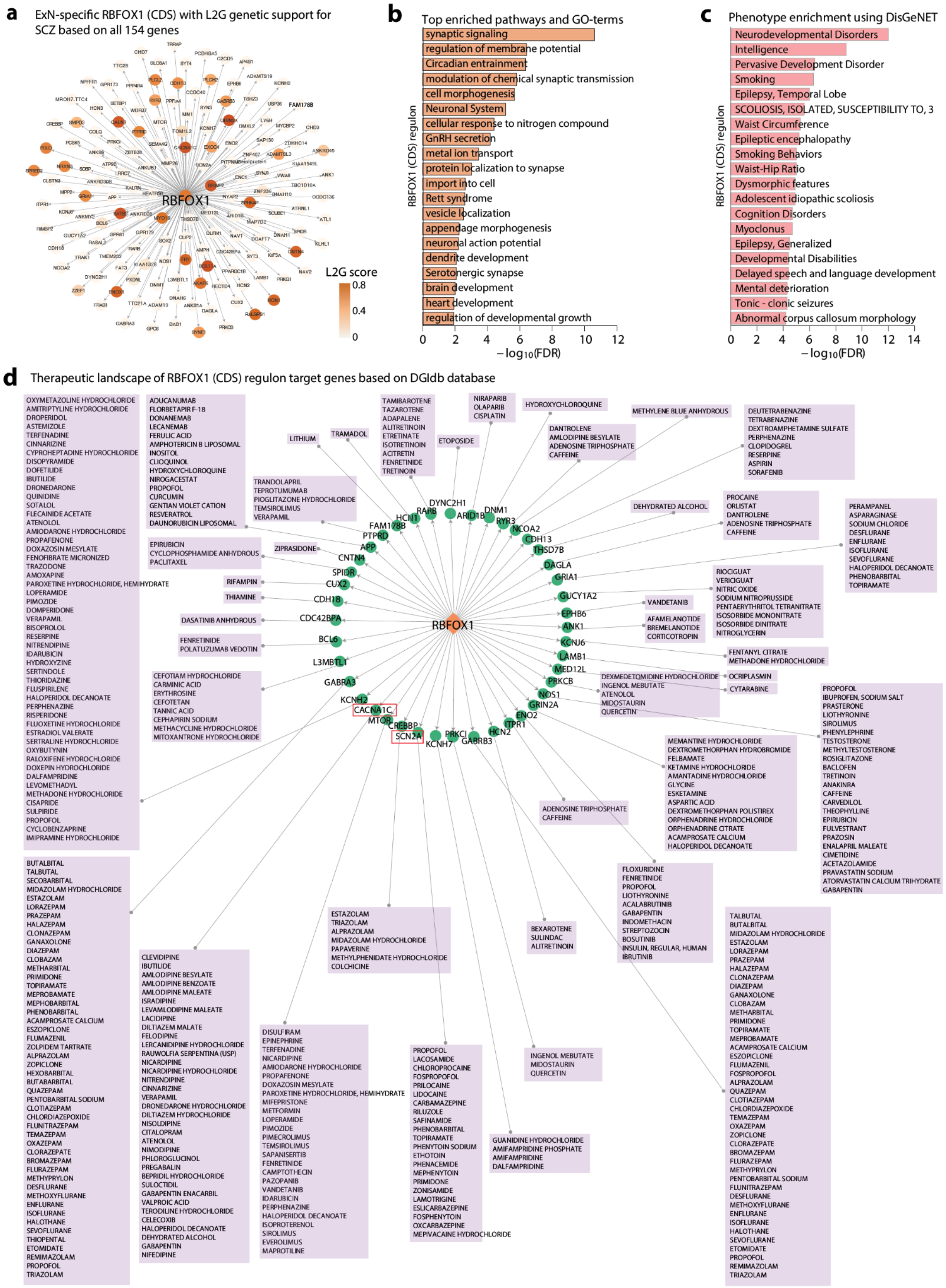


**Supplementary Fig. 31. RBFOX1(CDS)-centered regulatory program and functional characterization.**

(a) ExN-specific RBFOX1(CDS)-centered regulon identified by scRBP. Network visualization shows predicted RBFOX1 target genes containing (U)GCAUG motif matches within coding sequence (CDS) regions. Node color reflects genetic support based on L2G prioritization scores.

(b) Functional enrichment analysis of RBFOX1 regulon targets (n = 154 genes). Gene Ontology and pathway enrichment highlight neuronal and synaptic processes, performed by using Metascape (https://metascape.org/).

(c) Phenotype enrichment analysis of RBFOX1 (CDS) regulon targets by using DisGeNET (http://www.disgenet.org). Enriched phenotypic annotations include neurodevelopmental, cognitive and psychiatric disorder–related traits.

(d) Drug–gene interaction landscape of RBFOX1 (CDS) regulon targets based on DGIdb annotation (https://dgidb.org/). A substantial fraction of targets are linked to FDA-approved drugs, including sodium-channel modulators targeting SCN2A (e.g., lamotrigine and carbamazepine) and CACNA1C-associated agents (e.g., haloperidol decanoate and valproic acid) (Highlighted in red boxes). The orange diamond denotes the RBP regulator (RBFOX1), green circles indicate target genes with FDA-approved drug associations, and purple boxes represent corresponding drugs.
